# Altered interactions between circulating and tissue-resident CD8 T cells with the colonic mucosa define colitis associated with immune checkpoint inhibitors

**DOI:** 10.1101/2021.09.17.460868

**Authors:** Molly Fisher Thomas, Kamil Slowikowski, Kasidet Manakongtreecheep, Pritha Sen, Jessica Tantivit, Mazen Nasrallah, Neal P. Smith, Swetha Ramesh, Leyre Zubiri, Alice Tirard, Benjamin Y. Arnold, Linda T. Nieman, Jonathan H. Chen, Thomas Eisenhaure, Karin Pelka, Katherine H. Xu, Vjola Jorgji, Christopher J. Pinto, Tatyana Sharova, Rachel Glasser, Elaina PuiYee Chan, Ryan J. Sullivan, Hamed Khalili, Dejan Juric, Genevieve M. Boland, Michael Dougan, Nir Hacohen, Kerry L. Reynolds, Bo Li, Alexandra-Chloé Villani

**Affiliations:** Center for Immunology and Inflammatory Diseases, Department of Medicine, Massachusetts General Hospital (MGH), Boston, MA, USA; MGH Cancer Center, Boston, MA, USA; Broad Institute of Massachusetts Institute of Technology (MIT) and Harvard, Cambridge, MA, USA; Harvard Medical School (HMS), Boston, MA, USA; Division of Gastroenterology, Department of Medicine, MGH, Boston, Massachusetts, USA; Division of Infectious Disease, Department of Medicine, MGH, Boston, MA, USA; Division of Rheumatology, Immunology and Allergy, Department of Medicine, Massachusetts General Hospital, Boston, MA, USA; Division of Hematology and Oncology, MGH, Boston, Massachusetts, USA; Department of Pathology, MGH, Boston, MA, USA; Clinical Research Center, MGH, Boston, MA, USA; Department of Surgery, MGH, Boston, MA, USA

## Abstract

Therapeutic blockade of co-inhibitory immune receptors PD-1 and CTLA-4 has revolutionized oncology, but treatments are limited by immune-related adverse events (IRAEs). IRAE Colitis (irColitis) is the most common, severe IRAE affecting up to 25% of patients on dual PD-1 and CTLA-4 inhibition. Here, we present a systems biology approach to define the cell populations and transcriptional programs driving irColitis. We collected paired colon mucosal biopsy and blood specimens from 13 patients with irColitis, 8 healthy individuals, and 8 controls on immune checkpoint inhibitors (ICIs), and analyzed them with single-cell/nuclei RNA sequencing with paired TCR and BCR sequencing, multispectral fluorescence microscopy, and secreted factor analysis (Luminex). We profiled 299,407 cells from tissue and blood and identified 105 cell subsets that revealed significant tissue remodeling in active disease. Colon mucosal immune populations were dominated by tissue-resident memory (T_RM_) *ITGAE*-expressing CD8 T cells representing a phenotypic spectrum defined by gene programs associated with T cell activation, cytotoxicity, cycling, and exhaustion. CD8 T_RM_ and effector CD4 T cells upregulated type 17 immune programs (*IL17A, IL26*) and Tfh-like programs (*CXCL13, PDCD1*). We also identified for the first time an increased abundance of two *KLRG1* and *ITGB2*-expressing CD8 T cell populations with circulatory cell markers, including a *GZMK* T_RM_-like population and a *CX3CR1* population that is predicted to be intravascular. These two populations were more abundant in irColitis patients treated with dual PD-1/CTLA-4 inhibition than those receiving anti-PD-1 monotherapy. They also had significant TCR sharing with PBMCs, suggesting a circulatory origin. In irColitis we observed significant epithelial turnover marked by fewer *LGR5*-expressing stem cells, more transit amplifying cells, and upregulation of apoptotic and DNA-sensing programs such as the cGAS-STING pathway. Mature epithelial cells with top crypt genes upregulated interferon-stimulated pathways, *CD274* (PD-L1), anti-microbial genes, and MHC-class II genes, and downregulated aquaporin and solute-carrier gene families, likely contributing to epithelial cell damage and absorptive dysfunction. Mesenchymal remodeling was defined by increased endothelial cells, both in irColitis patients and specifically in patients on dual PD-1/CTLA-4 blockade. Cell-cell communication analysis identified putative receptor-ligand pairs that recruit CD8 T cells from blood to inflamed endothelium and positive feedback loops such as the CXCR3 chemokine system that retain cells in tissue. This study highlights the cellular and molecular drivers underlying irColitis and provides new insights into the role of CTLA-4 and PD-1 signaling in maintaining CD8 T_RM_ homeostasis, regulating CD8 T recruitment from blood, and promoting epithelial-immune crosstalk critical to gastrointestinal immune tolerance and intestinal barrier function.

## Introduction

Therapeutic monoclonal antibodies against co-inhibitory immune receptors PD-1 and CTLA-4 to treat human malignancies have revolutionized immuno-oncology. Since 2011, a total of seven immune checkpoint inhibitor (ICI) therapies targeting CTLA-4, PD-1, and PD-L1 have been approved by the Food and Drug Administration to treat cancer. Unfortunately, these treatments are frequently limited by the development of immune-related adverse events (IRAEs), which are inflammatory side effects that can affect any organ system and range in severity from mild to potentially lethal. IRAEs occur in over 85% of patients on these therapies, leading to treatment discontinuation in 10-40% of patients (Wolchok et al. 2017; Postow, Sidlow, and Hellmann 2018).

ICI-related colitis (i.e. irColitis) is the most common, severe IRAE and develops in approximately 10-20% of patients on anti-CTLA-4 therapy, 1-2% of patients on anti-PD-1 therapy, and 15-25% of patients on dual CTLA-4 and PD-1 blockade (Wolchok et al. 2017; Postow, Sidlow, and Hellmann 2018; Beck et al. 2006; Naidoo et al. 2016). Histopathologically, ICI-associated colitis (irColitis) presents with increased intraepithelial lymphocytes, neutrophilic infiltrates, epithelial apoptosis, and/or isolated lymphocytic colitis (Chen et al. 2017; Verschuren et al. 2016). To date, little is known about the mechanisms leading to IRAEs, in part because mice treated with ICI-blocking antibodies do not develop significant IRAEs (Elsas, Hurwitz, and Allison 1999; Curran et al. 2010; Heul and Stappenbeck 2018). Correlational studies have suggested roles for circulating T regulatory cells, CD21^lo^ B cells, increased tumor necrosis factor alpha (TNF-α) and IL-17α, microbial dysbiosis, and early T cell receptor (TCR) diversification in blood (Perez-Ruiz et al. 2019; Callahan MK S. Tandon et al. 2011; Shahabi et al. 2013; Das et al. 2018; Chaput et al. 2017; Oh et al. 2017). irColitis is commonly treated with steroids as first line therapy and infusion biologics targeting TNF-α or the integrin α_4_β_7_ for steroid-refractory disease (Puzanov et al. 2017; Haanen et al. 2017). Currently there are no prospective trials to guide the management of irColitis, and it is unclear what long-term effects these immunosuppressive treatments have on cancer treatment and likelihood of progression, though some studies suggest that using high dose steroids to treat IRAEs blunts ICI anti-tumor efficacy (Faje et al. 2018; Arbour et al. 2018). An improved understanding of IRAE pathogenesis is urgently needed to facilitate the early diagnosis of IRAEs and to inform new therapeutic targets that preserve anti-tumor responses.

We leverage single-cell RNA-sequencing (scRNA-seq) and single-nuclei RNA-sequencing (snRNA-seq) of colon mucosal biopsies and paired blood specimens collected from a well-characterized cohort of irColitis cases and controls to reveal the tremendous cellular heterogeneity that underlies this immune-mediated disease (Papalexi and Satija 2017; Gomes, Teichmann, and Talavera-López 2019) and define cell subsets that could ultimately be targeted for clinical diagnostics and tailored therapeutic intervention.

## Results

### Experimental Design

To define cellular populations associated with irColitis, we assayed the transcriptomes and T-cell and B-cell receptor repertoire of cells from endoscopic colon biopsies and matched peripheral blood specimens. We obtained samples from three complementary patient cohorts: 13 patients with irColitis, 8 patients on ICI therapy without colitis, and 8 healthy controls (**Fig. 1A-B**). In the majority of patients with irColitis, the disease, like ulcerative colitis, involves the rectum and extends proximally to affect the whole colon (Abu-Sbeih et al. 2018; Marthey et al. 2016). Because disease severity can differ between the ascending and descending colon, biopsies for this study were taken from the descending colon, sigmoid, and rectum and analyzed together (Geukes Foppen et al. 2018; Wright et al. 2019). Most patients on ICI therapy had melanoma or lung cancer, were treated with PD-1 blockade (either as monotherapy or in combination with anti-CTLA-4), and were biopsied within two weeks of symptom onset (**Fig. 1B, Table S1**). At the time of biopsy, 3 patients were on chronic, low-dose prednisone and 1 control patient was on concomitant chemotherapy (**Table S1**). Of the 8 control patients on ICI therapy, 4 had ICI-related gastritis and/or enteritis with sparing of the colon.

**Figure 1.**
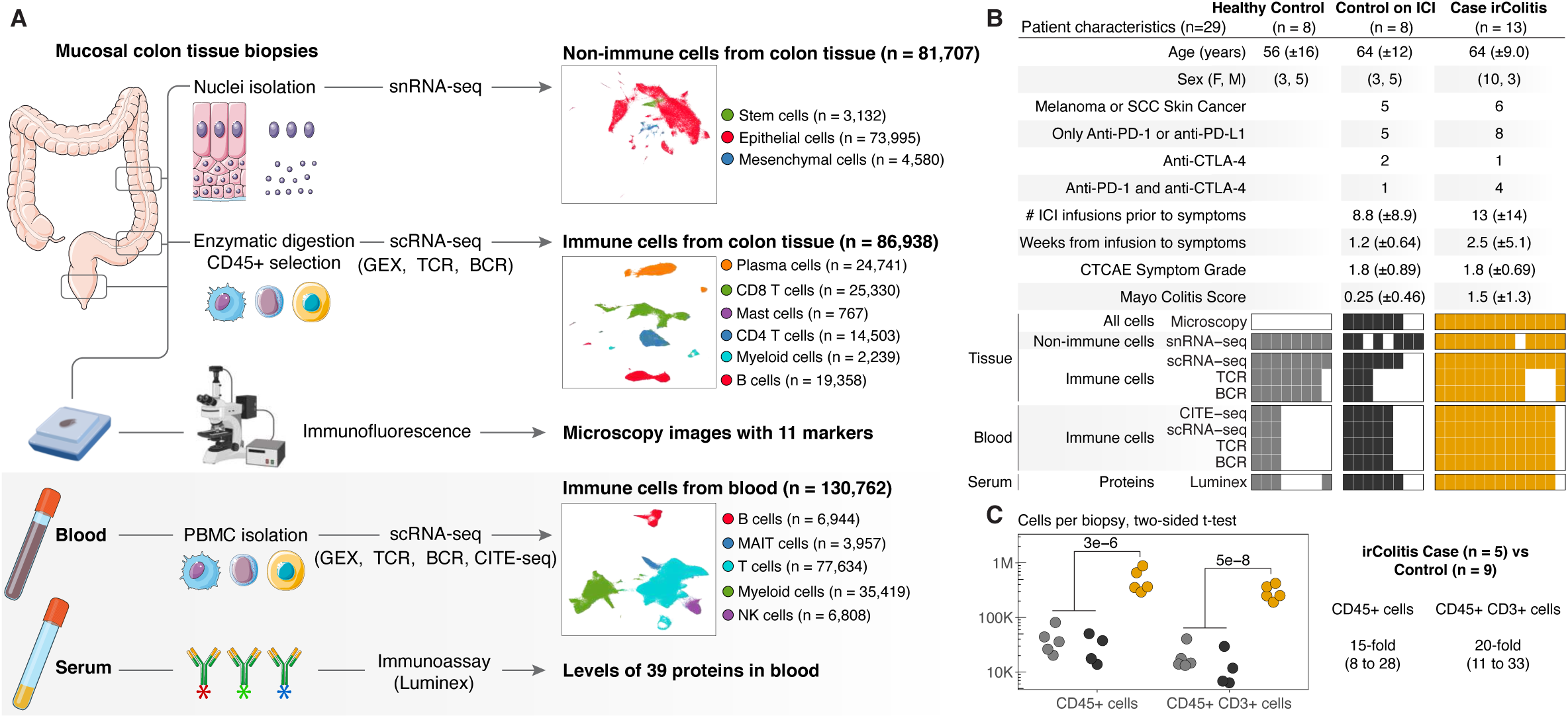
Overview of checkpoint colitis cohort. **A.** Sample processing pipeline overview. Mucosal colonic biopsies along with paired serum and blood specimens were collected from every patient at the time of diagnosis. From biopsies, single-nuclei RNA-sequencing (snRNA-seq) from 81,707 nuclei that passed quality control (QC) and single-cell RNA-sequencing (scRNA-seq) with paired T-cell receptor (TCR) and B-cell receptor (BCR) sequencing data from 86,938 cells that passed QC. Secreted factors were measured by Luminex ProcartaPlex in the serum, and scRNA-seq with paired TCR, BCR, and CITE-seq data from 130,762 cells that passed QC were generated from peripheral blood mononuclear cells (PBMCs) isolated from blood specimens. CITE-seq: Cellular Indexing of Transcriptomes and Epitopes by Sequencing. **B.** Cohort overview reporting the breakdown of the samples from 29 patients per clinical meta-data and per experimental approach used to analyze the cohort. ICI: immune checkpoint inhibitor; irColitis: immune-related colitis; CTCAE: Common Terminology Criteria for Adverse Events; SCC: Squamous cell carcinoma. **C.** Number of immune cells (defined by CD45 protein marker) and T cells (defined by CD3 protein marker) by fluorescence-activated cell sorting (FACS) analysis across healthy controls in grey, controls on ICI in black, and irColitis cases in orange. Two-sided Wilcoxon rank sum test p-values and 95% confidence intervals for fold-changes.

We processed several endoscopic colon mucosal biopsies from each patient in parallel to enable multiple readouts (**Fig. 1A-B; Methods**). For most patients, processing steps included enzymatic digestion of fresh tissue for scRNA-seq, nuclei isolation from flash-frozen tissue for snRNA-seq, and tissue fixation for microscopy. Paired serum and blood specimens were collected at the same time as the biopsies for secreted factor analysis (Luminex) and scRNAseq profiling of peripheral blood mononuclear cells (PBMCs) with paired TCR and BCR sequencing and 197 surface proteins (**Fig. 1; Methods**).

We generated single-cell transcriptomes from colon immune cells (86,938 cells from 27 patients), colon epithelial and mesenchymal cell nuclei (81,707 nuclei from 26 patients), and circulating PBMCs (130,762 cells from 21 patients) (**Fig. 1A-B**; **Table S1; Fig. S1**). We used canonical genes to group immune cells into broad cell types including B cells (*MS4A1*, *JCHAIN*), T cells (*CD3D, CD8A, CD4*), and myeloid cells (*LYZ*) (**Fig. S1).** Colon mucosal single nuclei were broadly identified as deriving from epithelial cells (*EPCAM, CDH1*), epithelial stem cells (*LGR5, SMOC2*), or mesenchymal cells (*VIM, COL3A1*) (**Fig. S1**). All clusters captured cells (or nuclei) from most patients, despite patient differences in ICI exposure and underlying cancer status (**Fig. S1**). We observed minimal differences in cell abundance and differential gene expression across cell lineages between the two different control groups (i.e. healthy controls and controls on ICI) (**Fig. 1C; Fig. S5-6,8-13,16**). These two control groups were therefore aggregated together for all remaining analyses.

### Cytotoxic CD8 T cells and CD4 Tregs cells are significantly more abundant in the colon mucosa of irColitis patients

Consistent with prior reports, patients with irColitis have significantly increased colon mucosal immune cells including T cells. (**Fig. 1C**) (Chen et al. 2017; Verschuren et al. 2016). Through flow cytometry analysis, we observed 15-fold (*P*=5e-6, 95% CI 8 to 28) more CD45^+^ immune cells and 20-fold (*P*=5e-8, 95% CI 11 to 33) more CD45^+^ CD3^+^ T cells in biopsies from irColitis cases than healthy controls or controls on ICI therapy (**Fig. 1C**). To further characterize these immune cell types and states, we clustered immune cell scRNAseq data into major cell lineages defined by canonical markers (i.e. CD8 T and innate cytotoxic lymphocytes, CD4 T cells, B cells, and myeloid cells) and assessed abundance differences between cases and controls (**Fig. 2; Fig. S1**).

**Figure 2.**
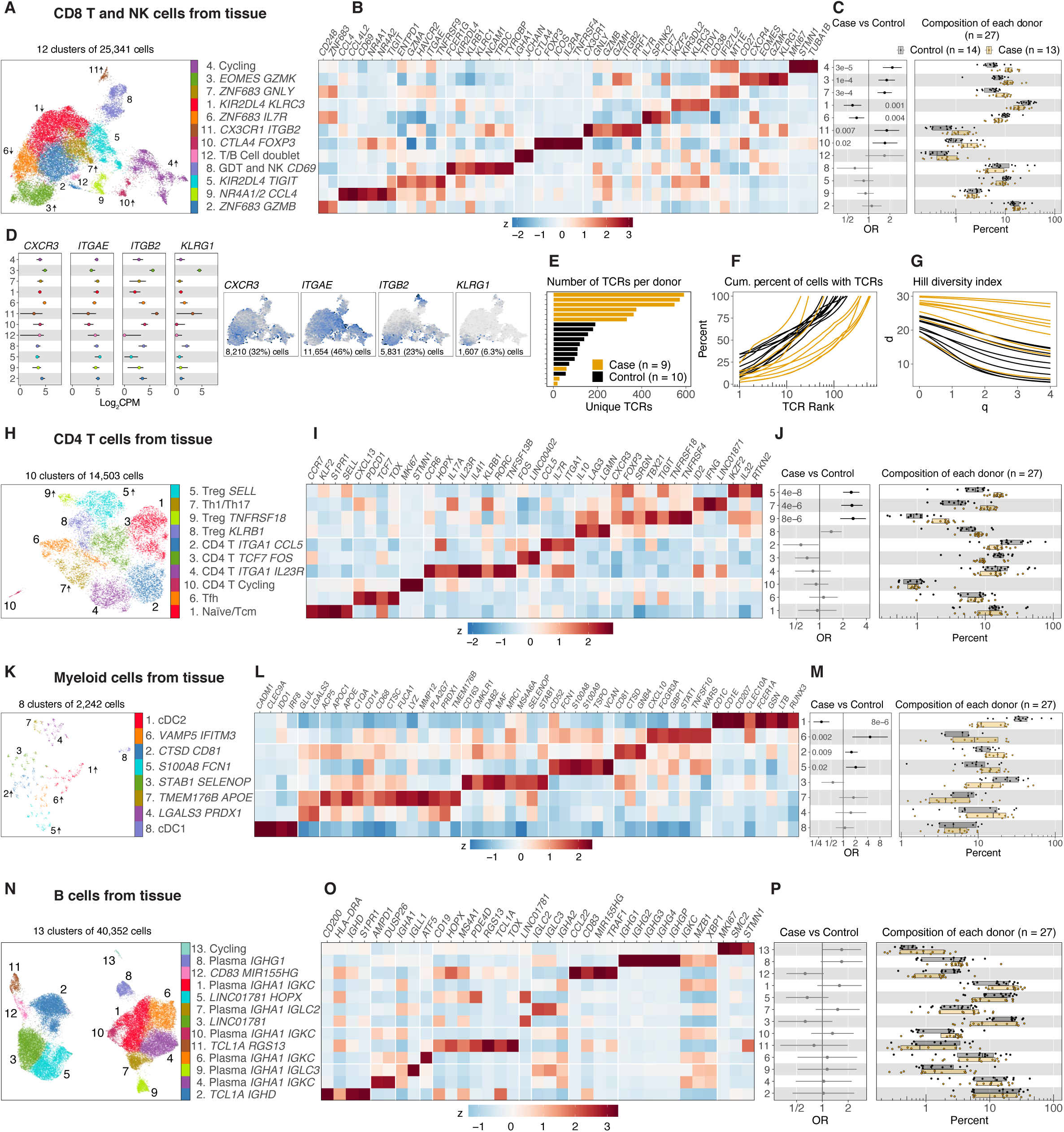
Analysis of cell type abundance and TCR clonality of immune cell subsets from colon tissue. **A, H, K, N.** UMAP embedding of **(A)** 25,341 CD8 T, gamma delta T (!”/GDT), natural killer (NK) cells, **(H)** 14,503 CD4 T cells, **(K)** 2,242 myeloid cells, and (**N**) 40,352 B cells. Colors indicate cell cluster identities. **B, I, L, O.** Normalized gene expression (mean zero, unit variance) for selected genes, showing relative expression across cell clusters from **(B)** CD8 T/GDT/NK, **(I)** CD4 T, **(L)** myeloid and **(O)** B cells. Rows of the heatmap are aligned to every cluster defined in the UMAP. **C, J, M, P.** Cell subset abundance differences between cases in orange (n=13) and controls in grey (n=14) across all (**C)** CD8 T/GDT/NK, **(J)** CD4 T, **(M)** myeloid and **(P)** B cell subsets. Boxplots show patient cell type compositions where each dot represents a patient. Composition of each patient is reported as the percent of cells from a patient in each cell cluster. Error-bars indicate logistic regression odds ratio (OR) for differential abundance of case cells for each cell cluster, and unadjusted likelihood ratio test p-values are shown. **D.** Gene expression level defined by Log_2_CPM of selected genes across CD8 T, GDT, and NK cell subsets is reported by cluster number and by feature plots where color indicates gene expression (Log_2_CPM) on the UMAP embedding. **E, F, G. (E)** Number of unique T cell receptor (TCR) clones per donor, **(F)** cumulative percent of cells with each TCR clone, ranked in order of abundance, and **(G)** Hill diversity index reported across cases in orange (n=9) and controls in black (n=10).

We analyzed 25,341 immune cells expressing *CD3D* and *CD8A* (**Fig. S1D**) and defined 12 clusters capturing a spectrum of cytotoxic lymphocytes including CD8 T cell subsets, gamma delta T cells (!”), and natural killer (NK) cells (**Fig. 2A; Fig. S2A-B; Table S2**). These cytotoxic cell types and states were annotated through marker genes specifically expressed in each cluster (**Methods**) and by cross-referencing those genes with published immune transcriptional signatures and legacy cell markers (**Fig. 2B; Tables S2-S3**) (Smillie et al. 2019; James et al. 2020; Corridoni et al. 2020; Boland et al. 2020). Most CD8 T cells captured in clusters 1, 2, 3, 5, 6, and 7 have a T_RM_ transcriptional phenotype (*ITGA1*, *ITGAE* or *KLRG1*, *CD69*) and share similarities with long-lived *ITGAE^lo^ KLRG1*^Hi^ *ITGB2*^Hi^ *GZMK*^Hi^ and *CD103*^Hi^ *KLRG1*^lo^ *IL7R*^Hi^ long- lived T_RM_ subsets identified human intestinal transplant studies (**Fig. 2B,D; Table S3**) (Bartolomé-Casado et al. 2019; FitzPatrick et al. 2021). These six T_RM_-like clusters could be further subsetted based on the gene expression of either (i) *KLRG1, ITGB2, GZMK, and EOMES* (cluster 3); (ii) *ITGAE and KIR2DL4* (clusters 1, 5); (iii) *ITGAE and ZNF683* (clusters 6, 2, 7). *KIR2DL4* encodes a killer-cell immunoglobulin-like receptor with both activating and inhibitory functions (Faure and Long 2002). The expression of *KIR2DL4*, *HAVCR2*, *ENTPD1*, and *TIGIT* genes are all enriched in CD8 T cluster 5 and comprise a dysfunctional CD8 T cell gene program described in melanoma (Li et al. 2018). *ZNF683* encodes the Blimp-1 homolog Hobit that is upregulated in long-lived effector CD8 T cells (Vieira Braga et al. 2015; Szabo, Miron, and Farber 2019).

We found seven CD8 T cell subsets with significant differential abundance in cases (patients with irColitis) versus controls (**Fig. 2C; Table S4**). Two types of resting T_RM_ cells expressing *IL7R and TCF7* were relatively depleted in irColitis cases: *KIR2DL4* T_RM_ cells (cluster 1, OR = 0.59, 95% CI 0.44 to 0.78) and *ZNF683* T_RM_ cells (cluster 6, OR = 0.67, 95% CI 0.53 to 0.86). In contrast, irColitis cases were enriched for *EOMES*, *GZMK-*expressing cytotoxic CD8 T cells (cluster 3, OR = 1.8, 95% CI 1.4 to 2.4), *ZNF683*, *GZMB*-expressing cytotoxic CD8 T cells (cluster 7, OR = 1.7, 95% CI 1.3 to 2.1), *MKI67*, *STMN1-*expressing cycling CD8 T cells (cluster 4, OR = 2.2, *P* = 3e-05), and a mixed population of *CD4*^Hi^ *CD8A/B*^Hi^ T cells that express *FOXP3*, *CTLA4,* and *ICOS* and thus may represent a regulatory T cell population (cluster 10, OR = 1.7, *P* = 0.02). We also observed an overabundance of CD8 T cells (cluster 11, OR = 1.8, 95% CI 1.2 to 2.7) with a circulating, cytotoxic phenotype (*CX3CR1*, *KLF2*, *S1PR1*, *GZMB*, *PRF1*, *GNLY*) that resemble previously described human *CX3CR1*^Hi^ cytolytic effector memory CD8 T cells confined to blood (**Fig. 2C-D**) (Buggert et al. 2020).

ICI therapy significantly remodels the TCR repertoire of circulating and tumor infiltrating CD8 T cells, and CD8 TCR clonotype diversity correlates positively with anti-tumor ICI responsiveness (Kidman et al. 2020). Since the impact of irColitis on colon TCR repertoire diversity was unknown, we performed single cell TCR-αβ profiling to further define the clonal and functional interrelationship of unique CD8 T cell subsets (**Fig. 2E-G; Fig. S3A-B; Methods**). Sequenced CD8 TCR clones were highly diverse with the majority of T cells expressing unique TCR-αβ pairs (**Fig. 2E**). Compared to controls, patients with irColitis demonstrated strikingly increased CD8 TCR clonal diversity with larger numbers of unique TCR sequences (**Fig. 2F-G**).

Sub-clustering analysis was performed on 14,503 CD4 T cells. Of the 10 clusters observed (**Tables S2-3**), 3 were increased in irColitis (**Fig. 2H-J; Fig. S1D, S2C-D; Table S4)**. All 3 *FOXP3*- expressing CD4 T regulatory (Treg) cell clusters were overabundant in irColitis relative to controls without colitis. These included: *SELL* expressing follicular-like Tregs (cluster 5, OR = 2.6, 95% CI 2 to 3.3), immunosuppressive-like Tregs expressing *KLRB1*, *LAG3*, and *IL10* (cluster 8, OR = 1.4, 95% CI 1 to 1.9), and Tregs expressing *TNFRSF4* and *TNFRSF18* (cluster 9, OR = 2.7, 95% CI 1.8 to 3.9). Effector CD4 T cells expressing transcriptional features of Th1-like cells (*CXCR3*, *IFNG*, *TBX21*) and Th17-like cells (*IL17A*) were also overabundant in irColitis cases (cluster 7, OR = 2.6, 95% CI 1.9 to 3.6) (**Table S4**). As observed in CD8 T cells, CD4 T cells with a resting T_RM_-like phenotype were relatively depleted in irColitis, including the cells from cluster 2 expressing *ITGA1*, *CCL5*, *IL7R* (OR = 0.57, 95% CI 0.34 to 0.97) and cluster 3 expressing *TCF7* (OR = 0.69, 95% CI 0.48 to 0.99). However, not all T_RM_-like CD4 T cell populations were depleted — a Th17-like T_RM_ population expressing *ITGA1*, *IL7R*, *IL23R*, and *IL411* had a similar abundance in irColitis cases and controls (cluster 4, OR = 0.82, 95% CI 0.48 to 1.4). CD4 TCR clonotype analysis showed no difference in TCR diversity between cases and controls **(Fig. S3C-E**).

### Inflammatory myeloid cells and cycling B cells are enriched in irColitis colon tissue

Given the important contribution of intratumoral myeloid cells to the efficacy of anti-PD-1 therapy (Garris et al. 2018; Chow et al. 2019; Strauss et al. 2020), we next assessed whether the colon mucosal mononuclear phagocyte (MP) compartment was altered in patients with irColitis compared to controls. Sub-clustering analysis of 2,242 myeloid cells identified eight distinct clusters, including two dendritic cell (DC) populations: *CLEC9A* expressing cDC1 cells (cluster 8), *CD1C* expressing cDC2 cells (cluster 1), and six MP populations that expressed genes associated with macrophages (C*D68*, *C1QA, C1QB, C1QC*) and monocytes (*CD14*, *FCG3RA*) (**Fig. 2K-M; Fig. S1D, S2E-F; Tables S2-S3**). Of these, three MP populations were enriched in colitis, including monocyte-like cells expressing *S100A8, FCN1, VCAN, CCR2* (cluster 5, OR = 2, 95% CI 1.2 to 3.5), and two populations co-expressing *CD68* and *C1QA* cells: cluster 6 cells (OR = 4.6, 95% CI 1.7 to 12) expressing a strong interferon-stimulated gene (ISG) signature (e.g., *IFITM3*, *GBP1*, *STAT1*, *CXCL10*), and cluster 2 cells (OR = 1.6, 95% CI 1.1 to 2.3) expressing several cathepsin genes (*CTSD*, *CTSB*, *CTSZ*), which are known to be upregulated by inflammatory macrophages in inflammatory bowel disease (IBD) (Menzel et al. 2006) (**Table S4**). In contrast, both cluster 1 cDC2 (expressing *CLEC10A, CD207, FCER1A; OR = 0.29, 95% CI 0.19 to 0.46)* and cluster 3 MP cells expressing *STAB1*, *CD163*, and *MRC1* (OR = 0.56, 95% CI 0.29 to 1.1) were found to be underrepresented in cases versus controls.

Subclustering analysis of 40,352 B cells identified 13 distinct clusters, including plasma B cells expressing *MZB1* and *XBP1* (clusters 1, 4, 6, 7, 8, 9, 10), *CD19* and *TNFRSF13B* expressing memory-like B cells (clusters 3 and 5), activated germinal center-like B cells expressing *CD19* and *CD83* (cluster 12), follicular B cells expressing *CD19* and *TCL1A* (clusters 2 and 11), and cycling B cells expressing *MKI67* (cluster 13) (**Fig. 2N-P; Fig. S1D, S2G-H; Tables S2-3**). None of these cell clusters were significantly enriched in irColitis cases (**Fig. 2P; Table S4**). In patients with ulcerative colitis, class switching to IgG plasma B cells has been observed along with an associated increase in the IgG to IgA plasma cell ratio. (Baklien and Brandtzaeg 1975; Scott et al. 1986). While *IGHG*-expressing plasma B cells were increased in irColitis patients, this finding was not significant (Wilcoxon Rank Sum Test *P* = 0.33), so we did not observe a difference in the ratio of IgG-to-IgA plasma cells (**Fig. 2P; Fig. S3H**). We did not find any differences in BCR diversity between irColitis cases and controls (**Fig. S3F-G, I**). Notably, irColitis developed in one patient receiving anti-PD-1 therapy for lymphoma and who had been treated with B-cell depleting CAR-T cell therapy (**Table S1**, patient C14*). The presence of expanded CD8 T cells expressing *GZMB* and *IFNG* and the complete absence of B cells in this patient (**Fig. S4**) suggest that tissue B cells are not necessary for the development of irColitis in all patients. Given the subtle proportional differences in B cell subsets and the ability of mucosal CD8 T cells to expand in the absence of B cells (in one patient), it is likely that B cells are not primary drivers of irColitis pathology.

### Colon mucosal T cells in irColitis are defined by the transcriptional co-regulation of genes implicated in T cell receptor signaling, cytotoxicity, and cell migration

To identify dominant gene expression programs associated with irColitis pathogenesis across CD4 and CD8 T cell populations, we performed differential gene expression analysis in irColitis cases versus controls at the level of cell lineage (CD4, CD8) and individual clusters (**Fig. 3**). Among the 12 CD8 T cell/cytotoxic lymphocyte clusters defined in colon tissue, 5 (i.e. clusters 1, 2, 4, 5, and 7) were enriched for differentially expressed genes (**Fig. 3A**) across several biological themes, including ISGs, nucleic acid sensors, T cell activation, antigen presentation, TCR signaling, apoptosis, and metabolism (**Fig 3**; **Fig. S5; Table S5**). Notably, NK and GDT cells (cluster 8) had few differentially expressed genes compared to CD8 T cells expressing TCR alpha/beta suggesting that innate cytotoxic lymphocytes undergo less dynamic transcriptional regulation in disease (**Fig. 3A**).

**Figure 3.**
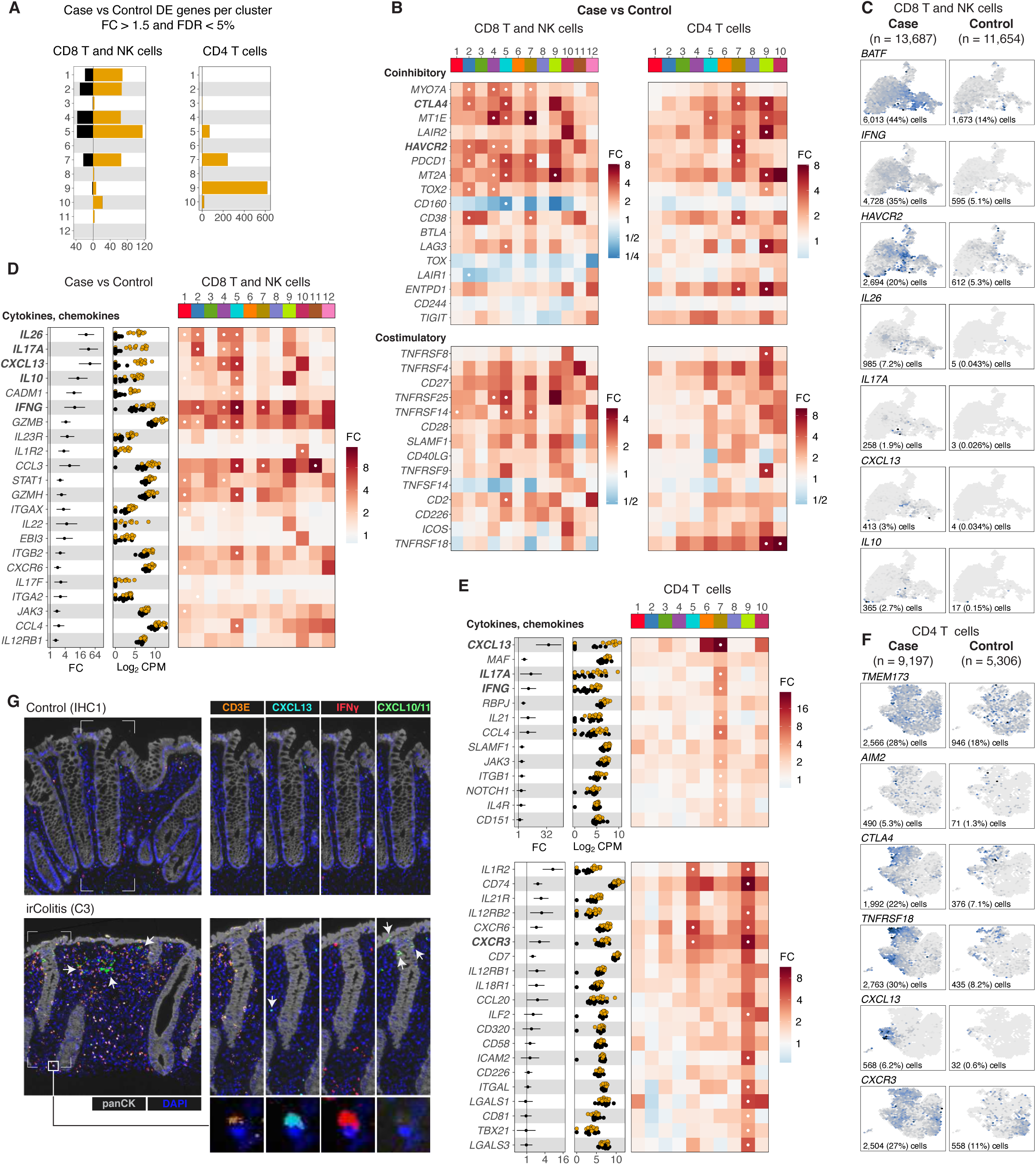
irColitis case versus control differential gene and protein expression across CD8 and CD4 T cell subsets. **A.** Bar plot of the number of differentially expressed (DE) genes per CD8 and CD4 T cell cluster (fold change (FC) greater than 1.5 and false discovery rate (FDR) less than 5%). **B.** Heatmap of fold change differences between irColitis cases and controls across CD8 T/GDT/NK and CD4 T cell subsets defined in Figure 2, showing selected co-inhibitory and co-stimulatory genes. White dot indicates FDR <5%. **C, F**. Feature plots illustrating gene expression level of selected genes between cases (left column) and controls (right column) in **(C)** CD8 T/GDT/NK cells and **(F)** CD4 T cells. Feature plots use color to indicate gene expression (Log_2_CPM) in the UMAP embedding. Number and percentage of cells (from cases or from controls) with expression are reported at the bottom of each feature plot. **D, E.** Fold change (FC) and gene expression (log_2_CPM) for cases (orange) and controls (black) is reported for cytokine and chemokine genes across **(D)** CD8 T/NK cell subsets and **(E)** CD4 T cell subsets. Heatmap color indicates fold-change difference between irColitis cases and controls across **(D)** CD8 T/GDT/NK and **(E)** CD4 T cell subsets defined in Figure 2 for selected co-inhibitory and co-stimulatory genes. White dot indicates FDR <5%. **G.** 6-color multispectral fluorescence panel depicting the RNA expression level of *CD3E* (orange), *CXCL13* (aqua), *IFNG* (red), and *CXCL10/11* (green), protein immunofluorescence of panCK (grey), and DAPI (blue) across HC1 control subject in the upper panel and C3 irColitis case in the bottom panel. Bottom box highlights co-expression of *CD3E*, *CXCL13*, and *IFNG* in the same T cell observed in the C3 irColitis case.

Many genes differentially expressed from irColitis patients were broadly expressed across most CD8 T cell subclusters, highlighting shared transcriptional programs common to diverse CD8 T cell subtypes. For example, DGE analysis showed that most CD8 T cells from irColitis patients all upregulated genes mediating cytotoxicity (*GZMB*, *IFNG*, and *EBI3* ), cytokine signaling (*STAT1*, *JAK3*, *IL23R*, *IL12RB1*), and CD8 T cell homing (*ITGA2*, *ITGB2*, *ITGAX*, *CADM1*, *CXCR6*) (**Fig. 3B-D, Fig. S5, Table S5**). In contrast, other differentially expressed genes were cluster-specific (**Table S5**).

DGE analysis in CD8 T cells revealed that *ITGAE*^Hi^ CD8 T_RM_ and T_RM_-related effectors upregulated different transcriptional programs in irColitis than the two *ITBG2* and *KLRG1*- expressing CD8 T cell populations with high circulatory marker expression (clusters 3 and 11; **Fig. 2A-D; Table S5**). CD8 T_RM_ cluster 1 expressing *KIR2DL4* (**Fig. 2A-B**) and T_RM_-related effectors expressing (CD8 T clusters 2, 7, 5, and 4; **Fig. 2A-B**) all showed significant numbers of differentially expressed genes in irColitis (**Fig. 3A**). Strikingly, all of five of these CD8 T populations upregulated the Th17 genes *IL17A*, *IL26*, and *IL23R* and the T follicular helper (Tfh) chemokine CXCL13, which participates in lymphoid structure organization by acting as a chemoattractant for immune cells expressing CXCR5 or CXCR3 (**Fig. 3C-D**). *IL17A*, *IL26*, and *CXCL13* expression was virtually absent from the *ITBG2* and *KLRG1*-expressing CD8 T cell populations (clusters 3 and 11), which had few differentially expressed genes (**Fig. 3A-C; Table S5**). *BATF*, which encodes a transcription factor that controls Th17 differentiation, was highly upregulated in CD8 T cells from irColitis patients (**Fig. 3C**) and thus was identified as a possible transcriptional regulator of the type 17 transcriptional program in CD8 T cells; however, given its broad expression across diverse CD8 T cell populations, *BATF* expression is likely not sufficient to upregulate *IL17A* and *IL26*. The *HAVCR2*^Hi^ *TIGIT*^Hi^ CD8 T cell population 5 additionally showed strong induction of secreted immune effector molecules including the chemokines *CCL3/4*, cytotoxic genes (*IFNG*, *GZMB*, *GZMH*), and the immunoregulatory genes (*IL10, IL22*) (**Fig. 3C-D; Table S5**). These gene programs were also present to a lesser extent across other CD8 T cells including *ZNF683 GZMB*-expressing cluster 2 and cycling cluster 4.

Candidate DGE analysis showed that many co-inhibitory/T cell exhaustion genes (*HAVCR2, CTLA4, PDCD1, TOX2, CD38, LAIR2*) were the most strikingly upregulated in the *ITGAE*^Hi^ T_RM_-related CD8 T effectors in clusters 2, 4, 5, and 7 (**Fig. 3B-C**) and to a lesser extent across other CD8 T and innate cytotoxic lymphocyte clusters. In contrast, TCR co-stimulatory genes (*TNFRSF14, CD27, TNFSF4, TNFRSF4, TNFRSF25, TNFRSF8, SLAMF1, CD2*) were significantly upregulated across several CD8 T cell subsets (**Fig. 3B**). Together these data reveal that *ITGAE*^Hi^ CD8 T effectors undergo dramatic transcriptional regulation in irColitis and strongly upregulate Th17 and Tfh-like genes including *IL17A*, *IL26,* and *CXCL13*. In contrast, *KLRG1*^Hi^ *ITGB2*^Hi^ CD8 T cells related to circulatory CD8 T cells showed almost no differentially expressed genes, suggesting that their increased abundance in irColitis may be driven more by relocalization to tissue and/or bystander activation.

In contrast to CD8 T cells, only three clusters of CD4 T cells had a significant number of differentially expressed genes, including two Treg cell subsets — *SELL*-expressing Treg cells (cluster 5) and *TNFRSF18-*expressing Treg cells (cluster 9) — as well as Th1/17 cells (cluster 7) (**Fig. 3A; Fig. S6; Table S5**). All three of these clusters were expanded in irColitis (**Fig. 2J**). Notably, an additional Treg subset that was not significantly expanded in irColitis (cluster 8, OR = 1.4, 95% CI 1 to 1.9) had higher levels of inhibitory receptors *KLRB1* and *LAG3* and no differentially expressed genes in irColitis. Since most of the gene expression differences seen within the CD4 T cell compartment were accounted for by Th1/17 cluster 7 and *TNFRSF18* cluster 9 (**Fig. 3A**), we performed differential gene expression within each of these respective subclusters (**Fig. 3E-F**; **Fig. S6G-J**). Genes jointly upregulated by clusters 7 and 9 (**Fig. 3F; Fig. S6, Table S5**) included interferon-stimulated genes and nucleic acid sensors (*TMEM173, GBP1, IFI6, BST2*), MHC class II genes (*HLA-DRA*, *HLA-DPA1*), genes implicated in cytokine and chemokine signaling (*EBI3, GZMB, CCR5, TNFRSF18*), and genes involved in cellular metabolism, transcription, apoptosis and autophagy (*MT1E, MT2A, CREM, BATF, CASP7, PYCARD, ATG16L1*). Compared to Treg cluster 9, Th1/Th17 cells (cluster 7) uniquely upregulated Th17- related genes *IL17A* and *IL21*, Th1 gene *IFNG*, and Tfh gene *CXCL13*, which was also upregulated in Tfh cluster 6 (**Fig. 3E-F, Fig. S6G-H; Table S5**). *TNFRSF18*-expressing Treg cells (cluster 9) had the largest number of differentially expressed genes of any CD4 T cell population (N = 725 genes, FC > 1.5 and FDR < 5%), suggesting strong transcriptional remodeling in irColitis. Some of the genes upregulated in cluster 9 included canonical Th1 genes such as *TBX21*, *IL12RB1*, *IL12RB2*, and *CXCR3*. Notably, in irColitis *CXCR3* was induced across most CD4 T cells (**Fig. 3E-F; Fig. S6I-J; Table S5**), which may represent a shared mechanism of CD4 T cell retention in inflamed tissue. Other upregulated genes in cluster 9 included those responding to Th17 cytokines (i.e. *IL21R*), those that induce Th17 trafficking (*CCL20*), as well as genes impacting CD4 trafficking (*CXCR6, ITGAL*), TCR co-stimulation (*ICOS, TNFRSF14*), TCR co-inhibition or exhaustion (*TOX2, LAG3*), autophagy (*ATG101, ATG16L2*), apoptosis (*CASP2, CASP8AP2*), and metabolism (*IDH2, FABP5, FUCA*) (**Fig. 3E-F; Fig. S6I-J; Table S5**). Of note, many genes upregulated by CD4 T cells were also seen upregulated in CD8 T cells, including co-stimulatory, co-inhibitory, T cell exhaustion, interferon-stimulation, and cytotoxic genes in addition to *CXCL13* (**Fig. 3B-C; Fig. S5**). We confirmed through multispectral RNA *in* situ hybridization (ISH) that T cells co-expressing *CXCL13* and *IFNG* were strongly enriched in irColitis patients and present throughout the lamina propria (**Fig. 3G; Fig. S7**). In summary, CD4 and CD8 T cells from irColitis patients show dynamic transcriptional regulation of genes modulating TCR signaling and implicated in Th1, Th17, and Tfh biology.

### Mononuclear phagocytes in irColitis colon contribute to ISG signatures observed disease

Colon mucosal MP subsets, particularly cluster 6, also showed marked upregulation of several differentially expressed genes (**Fig. S8; Table S5).** Most of the differentially expressed genes were interferon-stimulated and included the CXCR3 chemokine ligands (*CXCL9*, *CXCL10*, and *CXCL11*) and *STAT1*, which is required for the induction of many ISGs, including the CXCR3 chemokines ligands. While *CXCL9* was significantly upregulated by all myeloid clusters, ISG^Hi^ cluster 6 MP cells showed the highest fold change in *CXCL10* and *STAT1* induction and the highest expression of *CXCL11*. RNA-ISH confirmed that cells co-expressing the epithelial marker panCK and cells in the lamina propria that are panCK negative express high *CXCL10/11* (**Fig. 3G**, **Fig. S7**). We predict based on our scRNA-seq data that MP cluster 6 has the highest *CXCL10/11* expression of any tissue immune population. Other secreted factors upregulated by MPs in irColitis included the neutrophil chemoattractant *CXCL1* and the inflammatory cytokines *IL27* and *IL32 (***Fig. S8G**).

Pseudobulk DGE analysis of all MP subsets (**Fig. S8; Table S5**) showed upregulation of inhibitory MHC-I binding leukocyte immunoglobulin-like receptors (LILRs) and MHC class I genes *HLA-A* and *HLA-B* (**Fig. S8G**). MHC class II genes were significantly upregulated only in ISG-expressing MP cells from cluster 6, *APOE-*expressing MP cells (cluster 7), and cDC1 cells (cluster 8). All MP subsets upregulated the PD-1 ligand, *CD274* (i.e. PD-L1) in irColitis along with varying levels of the TCR co-stimulatory genes *CD80* and *CD40*, which overall reflect highly activated myeloid cell states.

Since irColitis patients likely have increased absolute numbers of IgG plasma cells in irColitis (though not altered plasma cell IgG to IgA ratios; **Fig. S3H**), we also sought to determine if Fc gamma receptor (FcγRs) genes were differentially expressed in MP cells from irColitis patients. In IBD, B cells may exert pathologic effects related to elevated IgG plasma cells and induced expression of activating FcγRs, which can opsonize commensal-IgG immune complexes that activate the NLRP3 inflammasome and interleukin-1β (Castro-Dopico et al. 2019). To determine if the myeloid Fc!R repertoire is altered in irColitis, we examined the relative expression of activating (*FCGR2A*, *FCGR3A*, *FCGR3B*) and inhibitory receptors (*FCGR2B*) across all MP subsets and noted that the activating FcγRs *FCGR1A* and *FCGR3A* were both significantly upregulated in irColitis cases without significant changes in the activating *FCG2RA* receptor or the inhibitory *FCGR2B* receptor (**Fig. S8G**). While *NLRP3* was not upregulated across MP clusters, several components of the NLRP3 inflammasome pathway, including *CASP1*, *PYCARD*, *GSDMD*, and *IL18*, were upregulated in three *CD68*-expressing MP clusters (i.e., clusters 3, 6, and 7; **Table S5**), suggesting potential dynamic regulation of inflammasome-related pathways in some of the MP subpopulations that are both enriched (i.e. cluster 6) and depleted (i.e. cluster 3) in irColitis patients.

In contrast to MP and T cell subsets, B cells had very few differentially expressed genes, which included the B cell chemoattractant *CCL25* and two ISGs (*STAT1* and *PSME2*) (**Fig. S9, Table S5**). Plasma IgA B cell cluster 1 had the largest number of DE genes (**Fig. S9B**) that defined a metabolically quiet cell state in irColitis characterized by the downregulation of genes participating in cell growth and differentiation (A20 binding protein *TNFAIP3*), nucleic acid binding and regulation (*PDCD4, PRDM2*), intracellular transport (*LAMP1*), and metabolism (*MTR, COX14, GLS*) (**Table S5**).

### Phenotypic relationship and TCR clonal dynamics of colon mucosal and circulating immune cells in irColitis

We next sought to determine if any mucosal immune cell populations enriched in the colon mucosa of irColitis patients could be readily detected in circulation. In some patient populations receiving anti-PD-1 therapy for solid tumors, anti-tumoral CD8 T cells have been reported to originate from circulatory cells populations (Yost et al. 2019). The contribution of peripheral immune cells to tissue-specific IRAEs, like irColitis, is not well-understood. At the time of colonoscopy, we collected peripheral blood matched to the same patients from whom tissue scRNA-seq and snRNA-seq libraries were generated (**Fig. 1A; Table S1; Methods**). Isolated PBMCs were processed for scRNA-seq with paired TCR and BCR sequencing as well as CITE-seq measurement. The resulting 130,762 cells that passed quality control (QC) filters (**Methods**) were further sub-clustered into CD8 T cells, CD4 T cells, B cells, and MP cell lineages containing 39 unique cellular subsets (**Fig. 4; Fig. S1E-G, S10-13; Tables S2-S3**).

**Figure 4.**
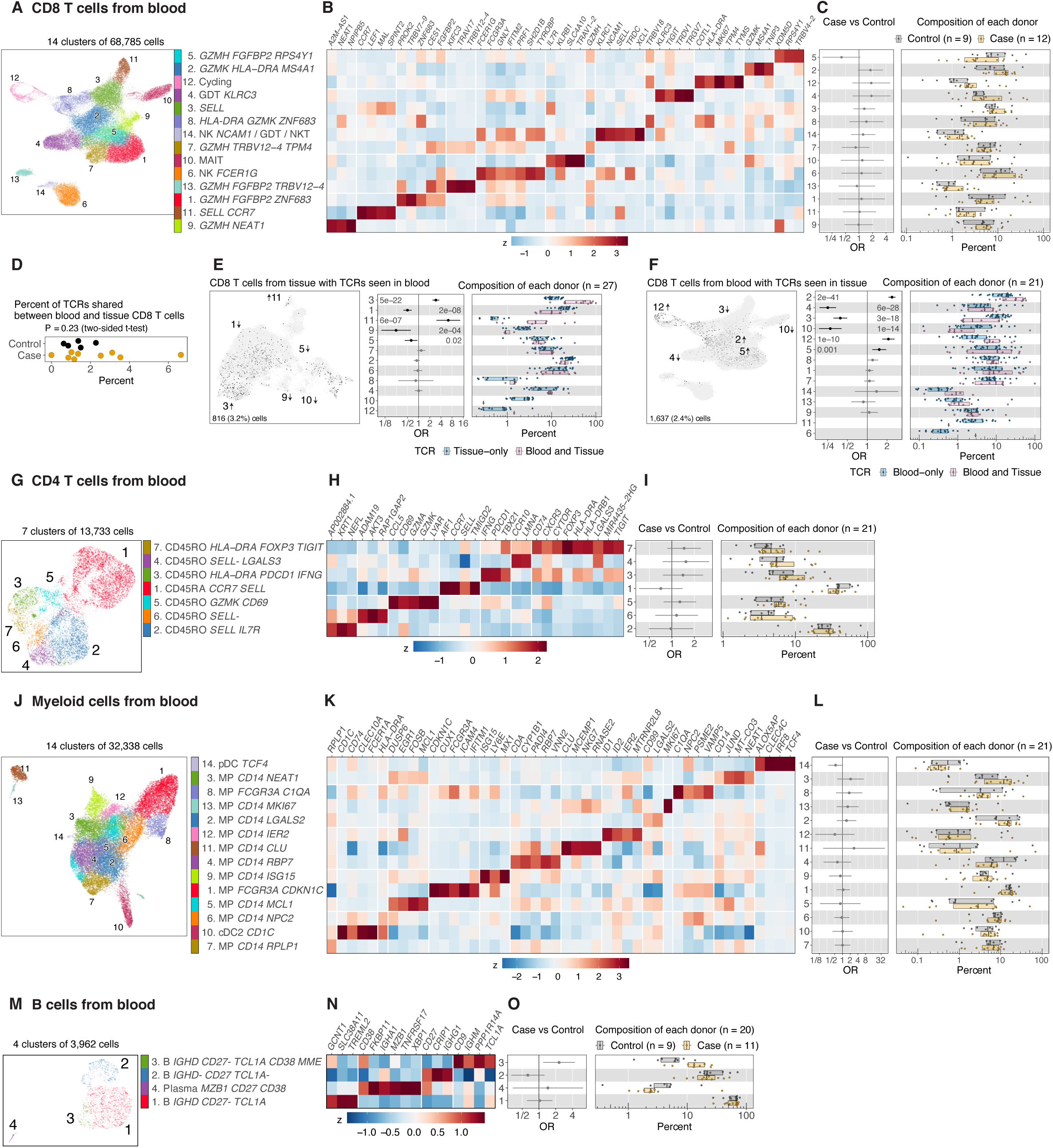
Analysis of cell type abundance and TCR clonality of immune cell subsets from paired blood samples. **A, G, J, M.** UMAP embedding of **(A)** 68,785 CD8T, GDT, and NK cells, **(G)** 13,733 CD4 T cells, **(J)** 32,338 myeloid cells, and (**M**) 3,962 B cells. Colors indicate cell cluster identities. **B, H, K, N.** Normalized gene expression (mean zero, unit variance) for selected genes, showing relative expression across cell clusters from **(B)** CD8 T/GDT/NK, **(H)** CD4 T, **(K)** myeloid and **(N)** B cells. Rows of the heatmap are aligned to every cluster defined in the UMAP. **C, I, L, O.** Cell subset abundance differences between cases in orange (n=13) and controls in grey (n=14) across all (**C)** CD8 T/GDT/NK, **(I)** CD4 T, **(L)** myeloid and **(O)** plasma/B cell subsets. Boxplots show patient cell type compositions, and each dot represents a patient. Composition of each patient is reported as the percentage of cells from a patient in each cell cluster. Error-bars indicate logistic regression odds ratio (OR) for differential abundance of cells from irColitis cases for each cell cluster, and unadjusted likelihood ratio test *p-*values are shown. **D.** Percent of CD8 T cells TCR clones shared between paired blood and tissue for each individual patient. Dots represent patients. irColitis case versus control, two-sided t-test (*p* = 0.23). **E.** Left panel shows CD8 T cells from colon tissue; color indicates cells with TCR clones observed in paired blood. Middle panel shows logistic regression odds ratio (OR) for differential abundance of TCR clones observed in tissue only or in both blood and tissue. Likelihood ratio test *p*-values are shown for clusters with FDR < 5%. Right panel shows the composition of each patient as percent of cells from each patient in each cell cluster. **F.** Left panel shows CD8 T cells from blood, color indicates cells with TCR clones observed in paired tissue. Middle panel shows logistic regression odds ratio (OR) for differential abundance of TCR clones observed in blood only or in both blood and tissue. Likelihood ratio test *p*-values are shown for clusters with FDR < 5%. Right panel shows the composition of each patient as percent of cells from each patient in each cell cluster.

The only circulating immune cell subset that might be differentially abundant in irColitis patients compared to controls is a population of B cells (cluster 3, OR = 2.3, 95% CI 1.2 to 4.4) expressing *IGHD*, *TCL1A*, *CD38*, and *MME* (**Fig. 4O; Table S4**) that is phenotypically similar to previously reported immature transitional B cells that represent an intermediate stage between immature bone marrow B cells and mature splenic B cells. Transitional B may represent an early checkpoint associated with the survival of autoreactive B cells (Dieudonné et al. 2019). Similarly to the tissue results, we did not find any difference in BCR diversity between irColitis cases patients and controls in circulation (**Fig. S14**).

Notably, none of the T cell or MP populations were significantly enriched or depleted from the blood of patients with irColitis compared to controls (**Table S4**). In contrast to cells from the tissue, there were no significantly differentially expressed genes in any T, B, or MP blood cell subsets between irColitis patients and controls (**Fig. S10-13; Table S5**). Three CD8 T cell subsets might have abundance differences in irColitis patients (**Fig. 4A-C; Table S4**), including: (1) a relative depletion in irColitis cases of *GZMH and FGFBP2-*expressing cells (cluster 5, OR = 0.41, 95% CI 0.15 to 1.1), and a relative enrichment in irColitis cases of (2) cycling cells (cluster 12, OR = 1.9, 95% CI 0.72 to 4.9) and (3) *GZMK* expressing cells (cluster 2, OR = 1.8, 95% CI 0.94 to 3.4). Cells from both clusters 2 and 5 share transcriptional features with populations previously described as circulating CD8 T effector memory cells, including an intravascular population of cytotoxic effector memory cells expressing *CX3CR1, GZMB, and PRF1* that do not recirculate (transcriptionally similar to cluster 5) and a population of effector memory cells expressing *CD27, TCF7, and GZMK* that recirculate in lymphatic and non-lymphatic tissue (transcriptionally similar to cluster 2) (Buggert et al. 2020). Among the seven CD4 T cell subsets from blood, irColitis might have slightly increased abundance of Tregs (cluster 7, OR = 1.5, 95% CI 0.87 to 2.5) and Th1- like CD4 effector cells (cluster 3, OR = 1.4, 95% CI 0.76 to 2.6), similar to the CD4 T effector cells observed in colon tissue (**Fig. 4G-I; Table S4**).

Paired single-cell TCR sequencing analysis enabled us to explore if there were any clonal relationships between mucosal and circulating CD8 T cells in irColitis patients and controls. Unlike in colon tissue, there was no association between irColitis status and CD8 TCR diversity, CD4 TCR diversity, or BCR diversity in human blood (**Fig. S14**). Examination of TCR sharing revealed that an average of 2% of a patient’s CD8 T cell clones were observed in both the colon mucosa and also in circulation (**Fig. 4D; Table S6**). This proportion of shared TCRs between colon tissue and blood was not significantly different in irColitis relative to controls (two-sided t-test *P*=0.23) (**Fig. 4D**). Two *KLRG1* and *ITGB2*-expressing tissue CD8 T subsets enriched in the tissue of irColitis patients (*CX3CR1-*expressing cluster 11 and *EOMES* and *GZMK*-expressing cluster 3) expressed circulatory markers (e.g. *KLF2*, *S1PR1*, *SELL*) and were thus predicted to traffic from circulation into colon tissue (**Fig. 2A-B; Table S3**). To determine if these two CD8 T cell subsets showed increased TCR clone-sharing with any CD8 T blood populations, we examined the relative frequency of tissue CD8 TCR detection in blood. Cluster 3 and 11 cells indeed had significantly increased TCR sharing with blood CD8 T cells (**Fig. 4E-F**). In contrast, some of the colon CD8 T cell subsets – including *KIR2DL4-*expressing T_RM_ cluster 1 cells, *KIR2DL4* and *TIGIT* expressing cluster 5 cells, and activated *CD69-*expressing cluster 9 cells – had significantly less TCR sharing with blood. Among the circulating CD8 T cell subsets, cycling cluster 12 cells, *GZMH* and *FGFBP2*-expressing cluster 5 cells, and *GZMK-*expressing cluster 2 cells all shared more TCRs with colon mucosal CD8 T cells in irColitis patients. Blood cluster 5 CD8 T cells were the most phenotypically similar to colon mucosal *CX3CR1-*expressing cluster 11 cells, which are predicted to be intravascular T effector memory cells (Buggert et al. 2020). Blood CD8 T cluster 2 cells were most phenotypically similar to tissue CD8 T *EOMES* and *GZMK* cluster 3 cells that have been described as recirculating, non-cytotoxic effector memory cells (Buggert et al. 2020). In contrast, blood naïve *SELL*-expressing cluster 3 cells, GDT cells in cluster 4, and MAIT cells in cluster 10 had relatively fewer CD8 T cell clones shared with tissue, suggesting that naïve and innate-like CD8 T cells from blood have little crossover with colon mucosal CD8 T cells in irColitis patients.

We additionally sought to determine if any specific secreted factors detected in blood were differentially associated with irColitis status. At the time of colonoscopy, serum was collected from 12 irColitis patients and 10 controls (4 healthy controls and 6 controls on ICI therapy) (**Fig. 1A**). 39 secreted factors were analyzed via the Luminex platform. These included factors upregulated in irColitis by colon mucosal T cells (CXCL13, IFN-γ, IL-7, IL-10, IL-17α, IL22, CCL3, CCL4), B cells (CCL25), and epithelial and/or myeloid cells (GM-CSF, CXCL1, CXCL2, CXCL8, CXCL9, CXCL10, IL-1α, IL-1β, IL-1RA, IL15, IL27) (**Tables S5, S7**). None of these 39 secreted factors were differentially expressed in irColitis cases versus controls (**Fig. S15**), suggesting that while these genes are upregulated in the colon tissue of irColitis patients, the respective increase in gene products could not be detected in circulation.

Together these blood data support that two blood CD8 T cell populations including a likely intravascular *CX3CR1* and a likely re-circulating *GZMK* CD8 T cell subset are detected at increased frequency in the colon mucosa of irColitis patients. Furthermore, the striking CD8 TCR diversity that defines the CD8 TCR repertoire in inflamed colon tissue is not reflected in blood. These findings are consistent with the prediction that most expanded CD8 T cell subsets in the colon mucosa of irColitis patients display a T_RM_ phenotype and thus are not predicted to recirculate.

### In irColitis, epithelial and mesenchymal cells show a strong interferon-induced gene signature associated with epithelial apoptosis, turnover, and absorptive defects

Because epithelial damage is a known histologic feature of irColitis (Verschuren et al. 2016; Oble et al. 2008; Berman et al. 2010; García-Varona, Odze, and Makrauer 2013), we next sought to define the epithelial and mesenchymal transcriptional programs that define active irColitis. Enzymatic digestion of endoscopic colon mucosal biopsies from patients with irColitis yielded dramatically fewer epithelial cells than biopsies from healthy control patients, potentially due to the relatively larger number of expanded immune cells and epithelial apoptosis (*data not shown*). To overcome this technical challenge, we employed a single nuclei RNA-seq (snRNA-seq) protocol that allowed for the isolation of intact single nuclei and associated ribosome-bound mRNA from frozen tissue samples (Drokhlyansky et al. 2020) (**Methods**). With this technique, we successfully sequenced 81,707 epithelial and mesenchymal nuclei that passed QC filters from 26 individuals (N=12 irColitis, N=8 healthy controls, N=6 controls on ICI therapy (**Fig. 1; Fig. 5A-D; Fig. S1I-K, S16A-B; Table S1)**. The majority of nuclei captured (∼93%) were derived from *EPCAM*-expressing epithelial cells, while the remainder were *VIM*-expressing mesenchymal cells (**Fig. S1I-K; Fig. 5A, D; Table S2)**. Using a combination of uniquely expressed genes (AUC ≥ 0.7) and known markers (Kanke et al. 2021; Smillie et al. 2019; Parikh et al. 2019), 23 predicted cell populations were identified (**Fig. 5A-D; Tables S2-3**) and classified as either immature epithelial cells (*LGR5, MKI67*), absorptive epithelial cells (high expression level of *EPCAM,* low expression of *SELENOP, CEACAM7, CEACAM1*, and *BEST4* enterocytes), mature absorptive epithelial cells (high expression of *EPCAM*, *SELENOP, CEACAM7, CEACAM1*, *AQP8*), secretory cells (*MUC2* goblet, *CHGA* enteroendocrine, *IL17RB* and *SH2D6* tuft cells), mesenchymal cells (*PECAM1* expressing endothelial cells, fibroblasts expressing high levels of *COL3A1* and low level of *CCL21*, inflammatory fibroblasts expressing high levels of *COL3A1* and *CCL21*), a mixed goblet and glial population (*MUC2* and *NRXN1* expressing cluster 17 cells), and a mixed immune and epithelial population (*EPCAM* and *PTPRC* expressing cluster 12 cells).

**Figure 5.**
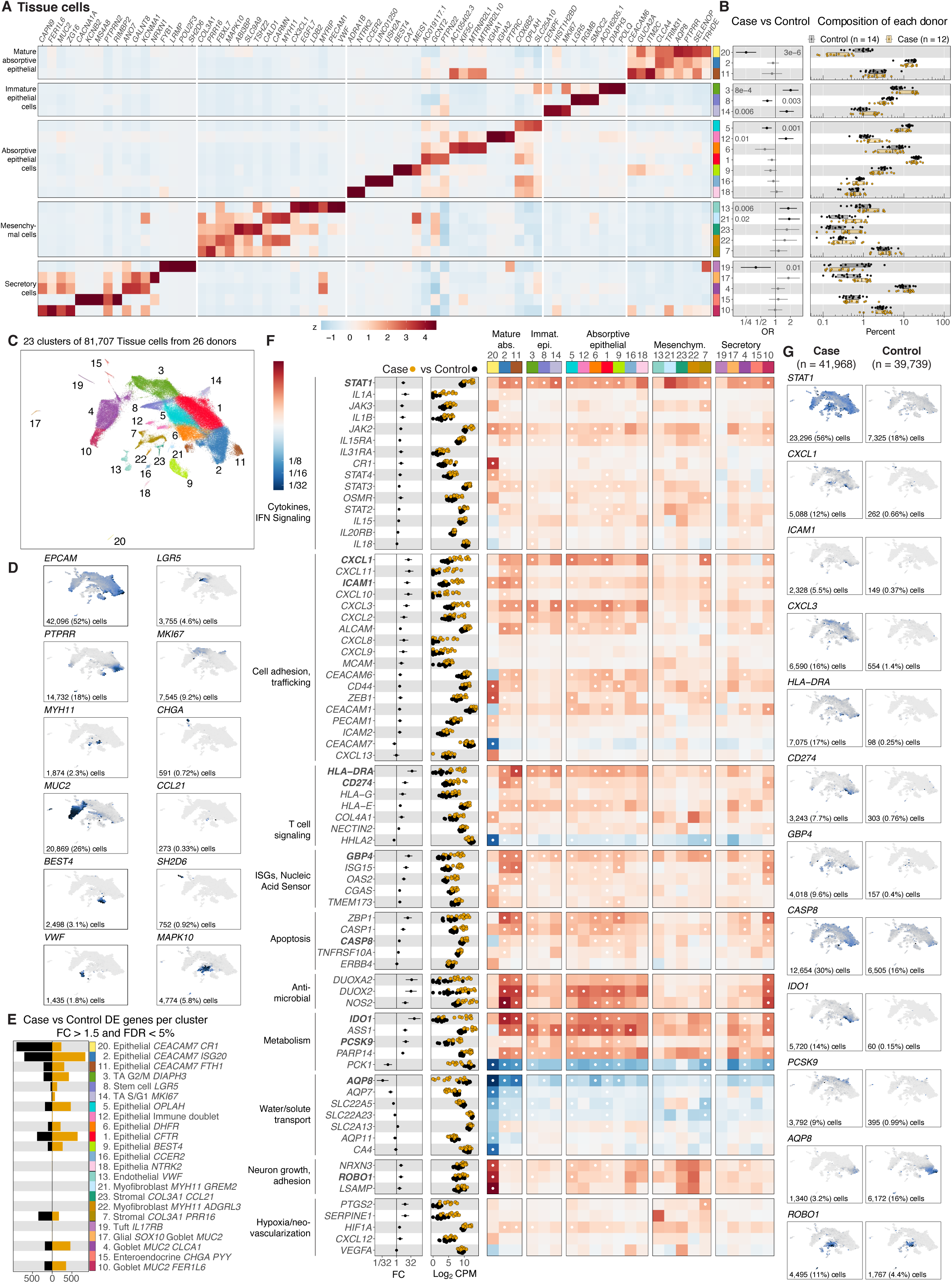
Analysis of cell type abundance of epithelial cell subsets from colon tissue. **A.** Normalized gene expression (mean zero, unit variance) for selected genes, showing relative expression across cell clusters of epithelial and mesenchymal cells. Rows of the heatmap are grouped into five families. **B.** Cell subset abundance differences between cases in orange (n=12) and controls in grey (n=14) across all non-immune cell subsets. Boxplots show patient cell-type compositions, and each dot represents a patient. Composition of each patient is reported as the percent of cells from a patient in each cell cluster. Error-bars indicate logistic regression odds ratio (OR) for differential abundance of case cells for each cell cluster, and unadjusted likelihood ratio test *p*-values are shown. **C.** UMAP embedding of 81,707 non-immune cells from 26 donors, color indicates one of 23 cell clusters. (See legend in panel **A** or **E**). **D.** Feature plots where color indicates gene expression (log_2_CPM) in the UMAP embedding. The selected lineage markers were used to guide initial cell subset annotation. **E.** Bar plot depicting number of differentially expressed (DE) genes per non-immune cell cluster (fold change (FC) greater than 1.5 and false discovery rate (FDR) less than 5%) **F.** Fold change (FC) and gene expression (log_2_CPM) for cases (orange) and controls (black) is reported for a set of genes organized across 10 biological themes defined on the left side of the heatmap. Heatmap color indicates fold change difference between irColitis cases and controls. White dot indicates FDR < 5%. **G.** Feature plots show gene expression (log_2_CPM) in cells from cases (left column) and controls (right column). Color shows gene expression in the UMAP embedding. Number and percentage of cells expressing each gene are reported at the bottom of each feature plot.

Abundance analyses assessing shifts in cellular population proportions between irColitis cases and controls (**Fig. 5B; Table S4**) revealed that there was a marked decline in colon epithelial stem cells (cluster 8, OR = 0.68, 95% CI 0.54 to 0.86), cluster 5 and 6 absorptive epithelial cells (cluster 5 OR = 0.67, 95% CI 0.53 to 0.84; cluster 6, OR = 0.61, 95% CI 0.38 to 0.96), mature epithelial expressing high levels of *CEACAM7* and *CR2* (cluster 20, OR = 0.25, 95% CI 0.16 to 0.4), and tuft cells (cluster 19, OR = 0.4, 95% CI 0.2 to 0.8). Meanwhile, irColitis samples had an increased abundance of transit amplifying epithelial cells in cluster 3 (OR = 2, 95% CI 1.4 to 2.9) and 14 (OR = 1.7, 95% CI 1.2 to 2.3) (**Fig. 5B; Table S4**). Most mesenchymal populations including *VWF* expressing endothelial cells (cluster 13, OR = 1.8, 95% CI 1.2 to 2.7) and *MYH11* expressing myofibroblasts (cluster 21, OR = 1.9, 95% CI 1.2 to 3.1; cluster 22, OR = 1.5, 95% CI 0.87 to 2.7) were more abundant in irColitis patients. Inflammatory fibroblasts expressing *CCL19* and *CCL21* were also more expanded in irColitis cases compared to controls though this was not significant (cluster 23, OR = 1.7, 95% CI 0.95 to 3). Cluster 23 fibroblasts were transcriptionally similar to inflammatory fibroblasts expressing *CCL19*, *CCL20* and *TNFSF14* that are enriched in ulcerative colitis patients (Smillie et al. 2019; Kinchen et al. 2018). Collectively these cellular abundance analyses support irColitis-induced remodeling of epithelial and mesenchymal cell compartments marked by a shift away from *LGR5*-expressing stem cells and towards cycling transit-amplifying cells, endothelial cells, myofibroblasts, and inflammatory fibroblasts.

We next performed DGE analysis to further characterize the impact of irColitis on epithelial and mesenchymal cell populations (**Fig. 5E-G; Fig. S16C-F; Table S5**). This analysis identified 8,185 differential expression results (FDR < 5%, fold-change > 1.5) across all 23 cell clusters (**Fig. 5E**). Several of the transcriptional gene modules significantly differentially expressed in irColitis cases comprised critical signaling pathways related to crosstalk between epithelial and mesenchymal cells with immune cells and comprised signaling programs predicted to impact epithelial barrier function (**Fig. 5F-G; Table S5**).

Some of the differentially-expressed immune signaling pathways in irColitis are related to increased inflammatory cytokine signaling, including cytokine receptors such as the IL-31 receptor heterodimer (*IL31RA*, *OSMR*), autocrine signaling loops (*IL15*, *IL15RA*), JAK-STAT signaling downstream of cytokine receptors, ISGs (*ISG15*, *GBP4, OAS2)*, cytosolic DNA sensors (*ZBP1, TMEM173*), and several components of the NRLP3 inflammasome pathway (*CASP1*, *IL18*, *IL1B*) (**Fig. 5F-G; Table S5**). *OSMR* was upregulated primarily in fibroblasts (cluster 7), which has also been observed in IBD (Smillie et al. 2019; Liu et al. 2015; Hegazy et al. 2017). Epithelial and mesenchymal cells both upregulated genes that modulate different components of immune function, such as antigen recognition (*HLA-G*, *HLA-E*, *HLA-DRA*, *CD74*), TCR signaling (*CD274/*PD-L1, *NECTIN2*, collagen genes), and immune trafficking including *ICAM1*, the CXCR3 chemokine ligands *CXCL10* and *CXCL11*, and neutrophil chemoattractants (*CXCL1*, *CXCL2*, *CXCL3*, *CXCL8*) (**Fig. 5F-G; Fig. S16C-D; Table S5**). Significant upregulation of genes with dual functions in cellular metabolism and suppression of T cell activation were also observed in irColitis cases (**Table S5**). These included: (1) *PCSK9*, which maintains cholesterol homeostasis and suppresses T cell activity by decreasing TCR recycling to the plasma membrane (Poirier et al. 2008; Yang et al. 2016; Yuan et al. 2021) (2) the tryptophan-metabolizing enzyme *IDO1*, which is induced by IFN-γ and TNF-α, participates in epithelial turnover, and exerts T cell suppressive activity in *IDO1*^+^ tumor cells (Yasui et al. 1986; Robinson, Hale, and Carlin 2005; Alvarado et al. 2019; Pflügler et al. 2020; Munn and Mellor 2016); and (3) *DUOX2* and *NOS2,* which, along with *IDO1*, comprises a transcriptional gene program that modulates microbial homeostasis at the colon luminal interface (Alvarado et al. 2019; Sarr et al. 2018). The upregulation of these varied immune transcriptional programs supports that in irColitis, interferon signaling mediated by nucleic sensing and inflammasome pathways likely induce both immune cell chemo-attractants and factors that attenuate immune responses such as the TCR co-inhibitor ligand *CD274* (PD-L1) and both MHC class II and non-canonical MHC class I genes that respectively mediate interactions with CD4 T cells and both CD8 T cells and innate cytotoxic lymphocytes. Collectively, these data support that PD-1 pathway inhibition may impair ISG-induced immune tolerance programs in irColitis.

We additionally identified several irColitis-specific gene programs impacting epithelial barrier function. These included the upregulation of multiple *CEACAM* adhesion molecules across epithelial cells and the significant down-regulation of aquaporin and small solute carrier gene families, which may directly contribute to impaired colonic water absorption associated with disease morbidity (**Fig. 5F-G; Table S5**). As predicted by previous histologic studies (Chen et al. 2017; Verschuren et al. 2016), colon epithelial cells from irColitis patients also displayed a range of apoptotic gene signatures. For example, *TNFRSF10A*/TRAILR-1 was markedly upregulated across multiple epithelial clusters along with *CASP1*, which comprises part of the NLRP3 inflammasome pathway, and *CASP8*, which contributes to TRAIL-mediated apoptosis. These results paralleled *TNFSF10* (TRAIL) upregulation in tissue CD8 T, CD4 T, and myeloid cells from irColitis patients (**Table S5**) and suggest a pathway by which infiltrating immune cells may impair epithelial cell survival.

Several irColitis-related gene programs induced in epithelial and mesenchymal cells showed variable expression along the epithelial crypt axis. Mature epithelial cell clusters 2 and 20 expressing *CEACAM7* had the largest number of DE genes and were predicted to reside towards the top of colonic crypts (based on expression of *SELENOP, CEACAM6*, *CEACAM7*) (**Fig.5F-G; Table S5**) (Parikh et al. 2019). Other mature absorptive epithelial populations (clusters 1, 5, 6, 9, 11) and both goblet populations (clusters 4 and 10) also had large numbers of differentially expressed genes in irColitis and were predicted to reside in the middle and top of crypts (based on intermediate *CEACAM6* and *SELENOP* expression) (Parikh et al. 2019). In contrast, undifferentiated epithelial stem cells (cluster 8) and TA cells (clusters 3 and 14) located towards the bottom of the crypt had relatively fewer differentially expressed genes in cases. In irColitis cases, ISGs (i.e. *ISG15*, *OAS1, STAT1, IDO1*, *CD274*/PD-L1) were most strongly upregulated in mature epithelial clusters 2 and 11, which are predicted to reside at the top of epithelial crypts based on the observed high expression level of *SELENOP*, *CEACAM6*, and *CEACAM7* (Parikh et al. 2019). Cluster 20 (*TNFAIP3 AQP8*) also expressed top crypt markers (*SELENOP*, *CEACAM7*) but was distinct from clusters 2 and 11 as it expressed fewer ISGs and more mesenchymal markers (*VIM*, *COL3A1)*.

Among the five mesenchymal cell clusters (**Fig. 5A**), only the *COL3A1*-expressing fibroblast cluster 7 showed differentially expressed genes in irColitis cases and expressed top crypt fibroblast markers, including *PDGFRA* and BMP ligands *BMP4/5/7* (Brügger et al. 2020). In irColitis patients, cluster 7 cells upregulated genes implicated in fibroblast differentiation (*FGF7*), epithelial differentiation (*BMP7*), ISGs (*STAT1*, *GBP1*), neutrophil chemoattractants (*CXCL1*), and prostaglandin synthesis (*PTGS1*, *PTGS2*), and downregulated multiple solute carrier genes associated with inflammatory bowel disease (IBD) in GWAS studies (*SCL22A5*, *SLC22A23*) (**Table S5**) (Leung et al. 2006; Serrano León et al. 2014). Multiple mesenchymal cell clusters from irColitis patients upregulated *ROBO1*, which may activate autophagy in intestinal stem cells and participate in epithelial-mesenchymal transition (Xie et al. 2020; Tan et al. 2020). While we noted that endothelial cells were more abundant in irColitis patients (**Fig. 5B; Table S4**), they had no significantly differentially expressed genes in irColitis cases vs controls (**Fig. 5E**). Together these results support that top-crypt epithelial cells display a strong ISG signature in irColitis marked by absorptive defects and that fibroblasts may contribute to epithelial cell dysfunction through the upregulation of immune chemoattractants, epithelial differentiation genes, and prostaglandins.

### Distinct cellular gene signatures associated with anti-PD-1 versus dual anti-PD-1/CTLA-4 therapy in irColitis

While dual PD-1 and CTLA-4 blockade has superior efficacy in treating many types of solid tumors (Rotte 2019), combination therapy is associated with a 5 – 10 times increased incidence of irColitis compared to anti-PD-1 monotherapy (Wolchok et al. 2017; Dougan 2017). In melanoma, compared to anti-PD-1 monotherapy, dual anti-PD-1/CTLA-4 therapy is preferentially associated with the expansion of activated, terminally-differentiated effector CD8 T cells and Th1-like CD4 T effectors, and the upregulation of CD8 T cell genes involved in mitosis and interferon signaling (Wei et al. 2019; Fairfax et al. 2020).

To better understand molecular drivers of irColitis in these different therapeutic contexts, we examined colon mucosal immune cell abundance and gene expression differences in irColitis patients on anti-PD-1 monotherapy (*N*=8) and dual PD-1/CTLA-4 blockade (*N*=4) compared to controls (*N*=12) (**Fig. 6; Fig. S17;Tables S4,S8**). Two *ITGB2* and *KLRG1*-expressing CD8 T cell subsets – *CX3CR1* cluster 11 and *EOMES GZMK* cluster 3 – were significantly more enriched in patients on dual therapy compared to those on anti-PD-1 monotherapy (**Fig. 6A; Table S4**). Both populations are unique in expressing genes associated with circulating cells (*KLF2*, *S1PR1*, *SELL*), in addition to displaying increased TCR sharing with circulating CD8 T cells (**Fig. 4E-F**), suggesting that these clusters may derive from circulating CD8 T cells. Very few genes were significantly differentially expressed between patients on ICI mono- and dual-therapy at a cell lineage pseudobulk or per-cluster basis across tissue immune cells (**Fig. 6G**; **Fig. S17; Table S8**). We did observe, however, that CD8 T cells from patients on dual therapy upregulated genes reflecting increased TCR signaling strength (*CD28*, *TNFRSF4*), homing to inflamed tissue (*SELL*, *ITGB2*), cytotoxicity (*GZMK*), and *LINC00861*, which is positively correlated with PD-1, PD-L1, and CTLA-4 levels in human cancers (**Fig. 6B-C; Table S8**) (Jiang et al. 2021; Hu et al. 2021). We additionally observed that patients on dual therapy had increased expression of *PAX*, which mediates CD8 T_RM_ retention in tumors via an interaction with the epithelial-binding integrin CD103 (Gauthier et al. 2017). Several of these genes were specifically upregulated in CD8 T cell cycling cluster 4 lymphocytes (*GZMK, LINC00861, PXN, SELL*) (**Fig. 6B-C; Table S8**), suggesting dynamic, therapy-specific gene expression programs in cycling cells. No significant abundance or gene expression differences were observed across other immune cell lineages in tissue (**Table S8**).

**Figure 6.**
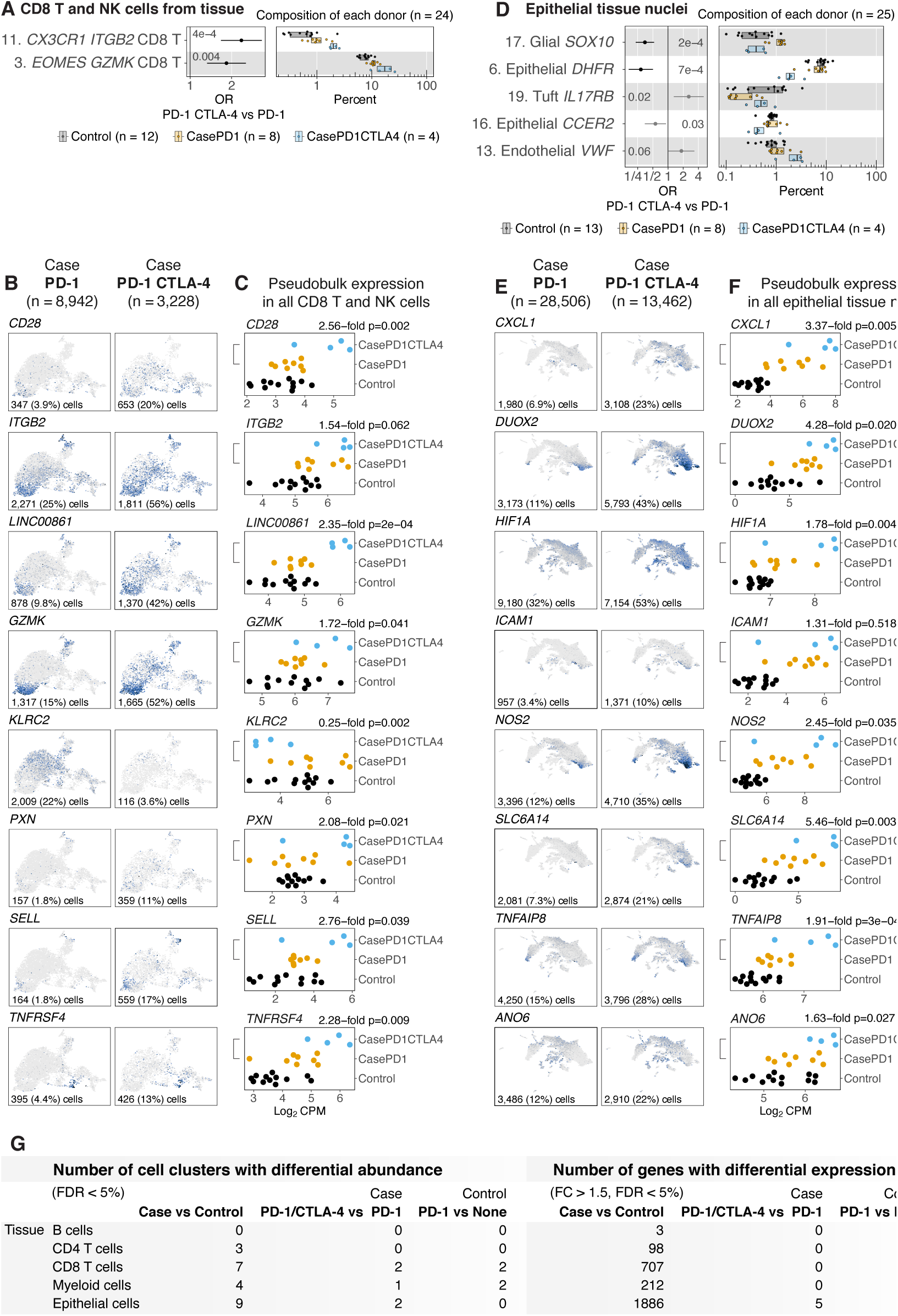
ICI drug analysis comparing cell subset abundance and differential gene expression across ICI treatment groups. **A, D.** Abundance analysis of (**A**) CD8 T/GDT/NK cell subsets and (**D**) epithelial/mesenchymal cell subsets. Controls in grey (n=12), irColitis cases on anti-PD-1 therapy in orange (n=8) and irColitis cases on anti-PD-1/CTLA-4 combination therapy in blue (n=4). Each dot represents a patient. Composition of each donor is reported as percent of cells from each patient in each cluster. Error-bars indicate logistic regression odds ratio (OR) for differential abundance of cells from patients on anti-PD-1/CTLA-4 versus patients on anti-PD-1 for each cell cluster, and unadjusted likelihood ratio test *p-*values are shown. **B, E.** Feature plots with gene expression (log_2_CPM). Selected genes with differential expression between cases on anti-PD-1 therapy (left column) and cases on anti-PD-1/CTLA-4 combination therapy (right column) across **(B)** CD8 T/GDT/NK cells and (**E**) across epithelial/mesenchymal cells in colon tissue. Color indicates gene expression (log_2_CPM) in cells from cases on anti-PD-1 or anti-PD-1/CTLA-4 therapy. Number and percentage of cells expressing each gene is at the bottom of each feature plot. **C, F.** Pseudobulk expression level for **(D)** all CD8T/GDT/NK cells and **(E)** all epithelial/mesenchymal cells from colon tissue for selected genes, stratified across 3 categories: irColitis cases on anti-PD-1/CTLA-4 combination therapy in blue, irColitis cases on anti-PD1 therapy in orange, controls in black. **G.** Number of cell clusters with differential abundance (FDR < 5%) for each cell type, for each contrast (i.e., Case vs Control, PD-1/CTLA-4 vs PD-1 within Cases, PD-1 vs None within Controls). Number of genes with differential expression (FDR < 5%, FC > 1.5, percent of cells with expression > 1%).

ICI therapy-specific differences in cell abundance and transcriptional signatures were also observed across epithelial and mesenchymal cell populations (**Fig. 6D-G; Fig. S18; Tables S4, S8**). For example, endothelial cells might be more abundant in dual therapy patients (cluster 13, OR = 1.8, 95% CI 1 to 3.3), while absorptive epithelial *DHFR*-expressing (cluster 6, OR = 0.29, 95% CI 0.17 to 0.5) and *CCER2*-expressing (cluster 16, OR = 0.56, 95% CI 0.35 to 0.91) cells were less abundant (**Fig. 6D; Table S4**). In contrast, compared to controls and irColitis patients on PD-1 blockade, irColitis patients on dual anti-PD-1/anti-CTLA-4 therapy a displayed proportional reduction in *SOX10* glial-containing cells (cluster 17, OR = 0.35, 95% CI 0.23 to 0.53) and increase in *IL17RB* tuft cells (cluster 19, OR = 2.6, 95% CI 1.3 to 5.1) (**Fig. 6D**). Differential gene expression analysis showed that irColitis patients on dual therapy preferentially upregulated genes implicated in cell growth and metabolism (*LCN2*), transcription (*LUCAT1*), cell adhesion (*CDH3*), immune cell homing to inflamed tissue (*CXCL1*), neutrophil homing and function (*CSF3*, *CXCL1*, *CXCL3*), and anti-microbial function *(NOS2*) (**Fig. 6E-F; Table S8**). Mature epithelial cluster 2 (*CEACAM7*, *SELENOP*, *AQP8,* ISG^Hi^ signature), and epithelial cluster 1 (*CFTR*) had the largest number of differentially expressed genes between irColitis patients on dual vs anti-PD-1 monotherapy (**Fig. S18B**). In irColitis patients on combination therapy, cells from these two clusters upregulated genes implicated in cell adhesion (*CD44, ITGA2*) and solute transport (*SLC6A14*). Cluster 1 epithelial cells showed particular upregulation of genes involved in anti-microbial responses (*DUOX, DUOXA1*) and NFKB signaling (*TNIP3, IRAK3*) (**Fig. 6E-F**; **Table S8**). In contrast, cluster 2 epithelial cells showed upregulation of genes involved in epithelial protection (*HIF1A*, *ANXA1*, *TNFAIP8*) (**Table S8**). Many of these differentially expressed genes – including *DUOX*, DUOXA2*, ICAM1*, and *CXCL1* – are ISGs, suggesting that increased interferon signaling may sustain irColitis in combination therapy. A summary of DGE across different cell lineages between cases/controls and between mono-/dual-therapy patients is summarized in **Fig. 6G**. Collectively, these results support that compared to irColitis in patients on anti-PD-1 monotherapy, irColitis in the setting of dual PD-1/CTLA-4 inhibition is marked by increased tissue recruitment of *ITGB2*-expressing CD8 T cells predicted to come from circulation, which may be mediated in part by increased interferon-stimulated immune homing ligands such as the ITGB2 ligand *ICAM1*.

### irColitis alters interactions between epithelial, myeloid, and T cells in colon tissue

We hypothesized that CD8 T cells may drive tissue damage in irColitis through direct interactions with epithelium and altered interactions with other immune populations. Almost all patients in our irColitis cohort were treated with anti-PD-1 inhibitors (12 of 13; **Fig. 1B**). Given the central role of PD-1 blockade in the pathogenesis of irColitis and the importance of PD-1/PD-L1 interactions in maintaining immune tolerance (Keir et al. 2008; Sharpe and Pauken 2018), we first sought to map interactions between all *PDCD1*/PD-1-expressing cells and cells expressing the PD-1 ligands *CD274*/PD-L1 and *PDCD1LG2*/PD-L2.

Consistent with a previous report (Nakazawa et al. 2004), T, epithelial, and mesenchymal cells from control patients showed low levels of *PDCD1*, *CD274*, and *PDCD1LG2* expression, though PD-1 and PD-L1 could not be detected by protein immunofluorescence (IF) in control tissue specimens (**Fig. 7A-B; Fig. S19,** *data not shown*). In contrast, *PDCD1* expression was upregulated in multiple activated CD4 and CD8 T cell subtypes (**Figs. 3 and 7A, Table S5**) in colon tissue from irColitis cases, and both PD-L1 and PD-1 could be readily detected by protein IF in irColitis tissue (**Fig. 7B; Fig. S19**). More specifically, PD-1 protein was detected in both CD8^+^ and CD8-^−^ T cells, the latter of which were presumed to be CD4 T cells based on our scRNAseq data showing comparably high *PDCD1* expression in only CD8 and CD4 T cells (*data not shown*). Although *PDCD1* could be detected at low expression levels in *FOXP3*-expressing CD8 and CD4 T cells (*data not shown*, **Fig. 2I**), we did not detect abundant FOXP3^+^ PD-1^+^ cell populations by protein IF microscopy (**Fig. 7B; Fig. S19**).

**Figure 7.**
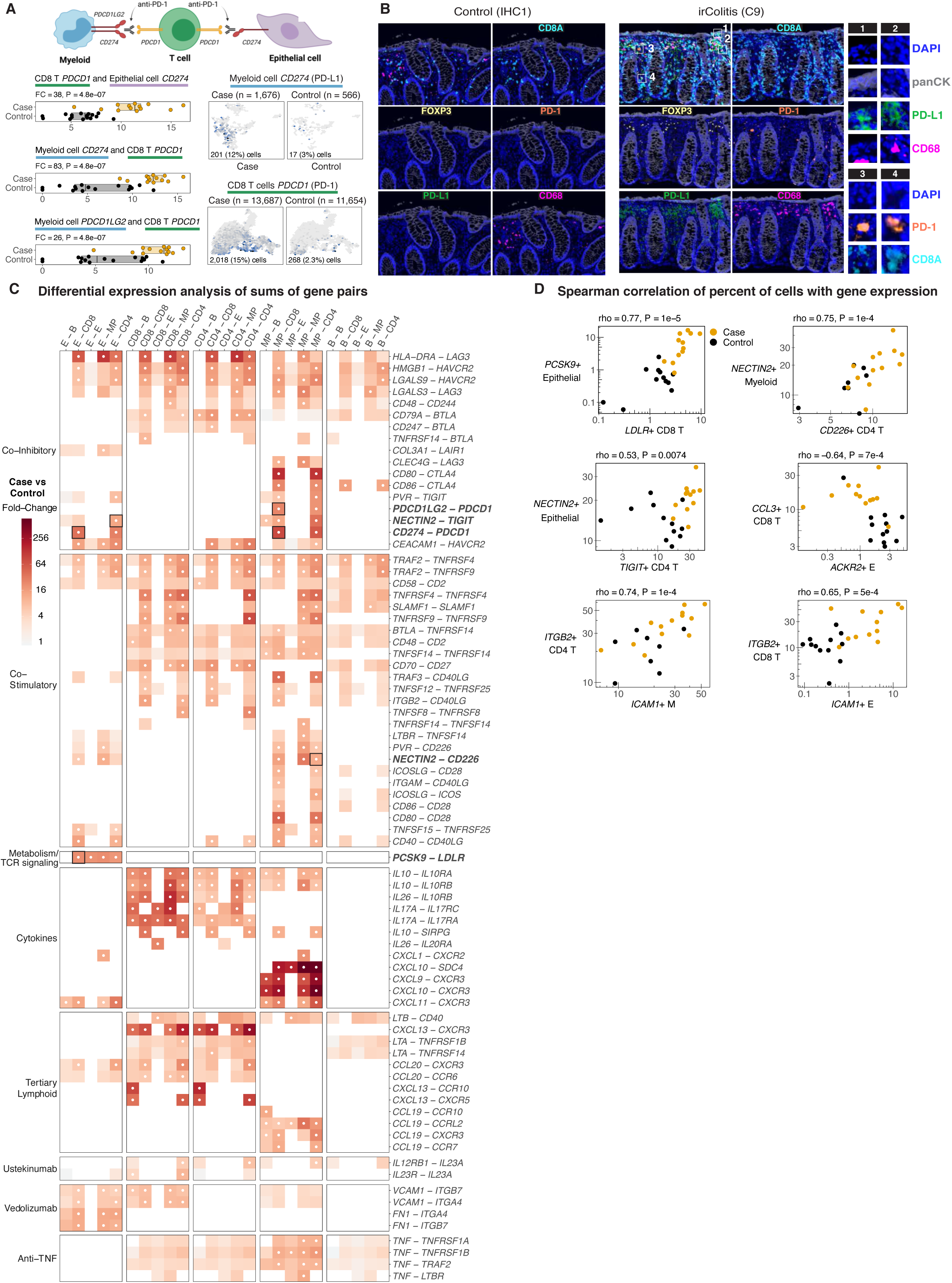
Cell-cell communication analysis. **A.** Schematic of a myeloid cell, T cell, and epithelial cell interacting via proteins encoded by *PDCD1* (PD-1), *CD274* (PD-L1), *PDCD1LG2* (PD-L2). The sum of log_2_CPM of *PDCD1* in CD8 T cells and log2CPM of *CD274* in Epithelial cells is 38-fold higher in irColitis cases relative to controls (P = 4.8e-7). Next, *CD274* in Myeloid cells and *PDCD1* in CD8 T cells, and *PDCD1LG2* in Myeloid cells and *PDCD1* in CD8 T cells. Top UMAP panel shows hexagonal bins with mean expression of *CD274* in Myeloid cells, stratified by irColitis case and control. Bottom UMAP panel shows *PDCD1* in CD8 T cells. **B.** Results from multispectral immunofluorescence staining of fixed colon mucosal tissue from HC1 control (left panel) and C9 irColitis case (right panel) with a 7-color panel that included DAPI (blue), panCK (grey), CD8A (aqua), FOXP3 (yellow), PD-1 (orange), PD-L1 (green), and CD68 (pink). Far right panels highlight co-detection of PD-L1 +/- CD68 and PD-L1 +/- panCK in individual cells (1, 2), and co-detection of PD-1 +/- CD8A in individual cells (3, 4). **C.** Differential expression analysis of sums of gene pairs. Heatmap color depicts irColitis case versus control fold-change for pairs of genes. Labels on the left indicate eight biological themes for the gene pairs. Columns indicate pairs of different cell lineages (E = epithelial, B = B cell, MP = myeloid cell, CD8 = CD8 T cell, CD4 = CD4 T cell). Pairs of genes in boldface are shown in panel **D**. White dots indicate FDR <5%. **D.** Spearman correlation of percent of cells with gene expression for pairs of genes depicted in panel **C**. Dots indicate patients. Axes indicate percent of cells in the lineage with expression of the gene. Spearman correlation and *p-*value is reported for each gene pair. **E.** Relative gene expression of candidate genes encoding drug targets for inflammatory bowel disease presented across five biological themes labeled on the left. Columns indicate immune, epithelial, and mesenchymal cell types. Color indicates scaled expression across cell types.

Analysis of PD-1/receptor gene pair products identified significant interactions between *PDCD1*-expressing CD8/CD4 T cells and both *CD274*-expressing epithelial and myeloid cells and *PDCD1LG2*-expressing myeloid cells (**Fig. 7A**). Across epithelial and mesenchymal populations, *CD274* was the most strongly upregulated gene in top-crypt epithelial clusters 2 and 11 (**Fig. 5F; Table S5**) and MP cluster 6 (**Table S5**). This is consistent with the high ISG signature (i.e. *ISG15*, *STAT1*, *GBP4*) observed in all three of these clusters as *CD274* is known to be induced by interferons (Kim et al. 2005). Protein IF confirmed that PD-L1 could be detected in top crypt epithelial cells; however, most of the PD-L1 signal in the colon mucosa was derived from cells in the lamina propria, including CD68^+^ cells located near the top of the crypt (**Fig. 7B**). Collectively, these results suggest that in irColitis patients, PD-1 inhibition may preferentially disrupt PD-1/PD-L1 interactions at the top of epithelial crypts, potentially leading to unmitigated cytotoxic T cell responses and interferon release at the luminal side of the crypt axis.

We next sought to map other T cell co-inhibitory and co-stimulatory receptor ligands across immune, epithelial, and mesenchymal populations. In irColitis patients, most interactions between T cell co-inhibitory/stimulatory receptors and their ligands were predicted between T cells and other T cell or myeloid populations, rather than between T cells and epithelial or B cells (**Fig. 7C; Tables S9-10**). In addition to *PDCD1*, several TCR co-inhibitory receptors were predicted to interact with more than one ligand, including LAG3 (*LAG3-LGALS3*, *LAG3-CLEC4G*), TIM-3 (*HAVCR2-HMGB1*, *HAVCR2-LGALS9*), BTLA (*BTLA-CD79A*, *BTLA-CD247*, *BTLA-TNFRSF14*), and TIGIT (*TIGIT-PVR*, *TIGIT-NECTIN2*). In turn, multiple ligands (*e.g. CD80*, *CD86*, *NECTIN2*) are predicted to have opposite effects on TCR signaling depending on the receptor they bind. For example, irColitis patients upregulated *NECTIN2-TIGIT* and *NECTIN2-CD226* gene pairs that are predicted to have co-inhibitory and co-stimulatory effects, respectively, on TCR signaling (**Fig. 7C-D;Tables S9-10**).

Additional analyses (**Methods**) predicted other unique cell-cell interactions in irColitis that may mediate tissue damage (**Fig. 7C-D; Fig. S20; Tables S9-10**). One of the strongest ligand/receptor pairs induced in epithelial and immune cells, respectively, from irColitis patients was *PCSK9*-*LDLR*, which may have a T cell suppressive effect (Benjannet et al. 2004; Lagace et al. 2006; Yang et al. 2016; Yuan et al. 2021). Several secreted ligand and receptor interactions were also identified to be significantly upregulated in irColitis cases compared to controls (**Fig. 7C; Fig. S20; Tables S9-10**). For example, T cell-specific cytokines including IL-26 and IL-17α are predicted to bind to receptors (IL10R2/IL20R1 and IL17RA/IL17RC, respectively) expressed across diverse cell types including myeloid cells, B cells, epithelial/mesenchymal cells, and different T cell populations expanded in irColitis cases. In contrast, in irColitis patients CXCR3 chemokine ligands *CXCL9*, *CXCL10*, and *CXCL11* were predominantly induced in MPs and predicted to interact with only immune cells based on expression of the canonical CXCL9/10/11 receptor *CXCR3* and the non-canonical receptor *SDC4*. We further observed that both T cells and MPs from irColitis cases over-expressed cytokines and chemokines capable of inducing tertiary lymphoid structures (*LTA, LTB*, *CXCL13*, *CCL19, CCL20*).

Putative interacting cell-cell interactions were also assessed using a complementary approach via Spearman correlations of up- and down-regulated receptor and ligand pairs across different cell types in irColitis patients and controls (**Methods**). This analysis detected several of the same cell-cell interactions predicted by differential expression of gene pairs (e.g. *PCSK9-LDLR*, *NECTIN2-TIGIT*, *NECTIN2-CD226*), in addition to identifying putative interactions between *ITGB2*-expressing CD8 T cells and *ICAM1*-expressing epithelial cells (**Fig. 7D; Fig. S20**). Because *ITGB2* is most highly expressed in CD8 T cell populations that may come from circulation (*CD8 T cell* clusters 3 and 11; **Fig. 2A, D**) and ICAM1 is most highly expressed in endothelial cluster 13 cells (**Table S3**), this predicted interaction may mediate homing of circulating CD8 T cell populations to inflamed colonic epithelium.

While multiple therapeutic monoclonal antibodies used to treat IBD have been repurposed to treat steroid-refractory irColitis, the mechanisms by which they induce disease remission are not fully understood (Johnston et al. 2009; Brahmer et al. 2018; Bergqvist et al. 2017; Thomas, Ma, and Wang 2021; Haanen et al. 2017; Puzanov et al. 2017). We therefore sought to define the ligand-receptor pairs disrupted by commonly available irColitis and/or IBD medications including ustekinumab (targeting p40 subunit common to IL-12 and IL-23), vedolizumab (targeting integrin α4β7), and anti-TNFα monoclonal antibodies (**Fig. 7C**). This analysis revealed the ustekinumab was predicted to target interactions between IL-23-producing T cells and immune cells expressing the IL-23 receptor, while vedolizumab was predicted to target α4β7^+^ immune cells binding *FN1* (on mesenchymal cells) and *VCAM1* endothelial cells (**Fig. 7C**). As previously described in IBD (Levin, Wildenberg, and van den Brink 2016), anti-TNFα monoclonal antibodies may act through canonical receptors (*e.g. TNFRSF1A*, *TNFRSF1B*) and non-canonical receptors (*e.g. TRAF2*, *LTBR*) to primarily disrupt the effects of myeloid-derived TNFα on both immune and epithelial cells (**Fig. 7C**). These results provide evidence of complex, putative cell-cell interaction networks that fine-tune immune cell chemotaxis and TCR signaling strength in a manner that may vary along the length of the epithelial crypt axis, and allow for the identification lineage-specific transcriptional pathways targeted by commonly used and potentially novel irColitis therapies.

## Discussion

Here we perform in-depth analysis of paired colon tissue and blood to report the first comprehensive, high-resolution study of immune and non-immune cell perturbations in an IRAE. Through the use of complementary, multimodal approaches – including snRNA-seq, scRNA-seq with paired TCR/BCR and CITE-seq, multispectral microscopy, and secreted factor analysis – we analyzed 299,407 cells/nuclei encompassing 105 cellular populations across paired colon tissue and blood specimens, and identified 17 populations specifically associated with irColitis, including 8 T, 3 mononuclear phagocyte, 4 epithelial, and 2 mesenchymal cell states enriched in patients with irColitis. Our study reveals that irColitis is defined by expanded *IL26*-expressing CD8 Trm cells with Th17-like transcriptional features. In addition, we predict the recruitment and retention of cytotoxic CD8 T cells from blood via altered interactions with inflamed colonic endothelium and epithelium that may contribute to epithelial damage including absorptive defects. Lastly, we use these findings to nominate novel therapeutic drug targets that could be used to mitigate irColitis.

In irColitis, colon mucosal immune cell infiltrates are dominated by expanded CD8 T cells with T_RM_ features. While prior studies have differentiated between CD8 T cells that reside in the lamina propria (i.e. LP CD8 T) or within the epithelium (i.e. IELs) (Isakov et al. 2009; Luoma et al. 2020; Smillie et al. 2019), newer work supports that mucosal CD8 T cells transit between the LP and epithelium and share overlapping TCR clones and transcriptional programs (FitzPatrick et al. 2021; Bartolomé-Casado et al. 2019; Puzanov et al. 2017). scRNA-seq studies have previously identified two transcriptionally distinct populations of intestinal CD8 T cells that are *bonafide*, long-lived Trms defined by divergent expression of the E-cadherin receptors KLRG1 and CD103 (*ITGAE*), respectively (Bottois et al. 2020; Bartolomé-Casado et al. 2019; FitzPatrick et al. 2021). These two previously described cell subsets co-expressing either *KLRG1* and *ITGB2* or *ITGAE* and *KLRB1* were transcriptionally similar to CD8 T cell cluster 3 (*EOMES*, *GZMK*, *KLRG1*) and clusters 6 and 2 (*ITGAE ZNF683*), respectively. TCR analysis showed overlapping TCR clones shared by multiple CD8 T cell populations expressing *ITGAE* and *ZNF683* (clusters 2, 6, 7 and cycling population 4), of which clusters 4, 6, and 7 are enriched in irColitis, suggesting a shared ontologic origin of these CD8 T cell clusters. *ITGAE-*expressing CD8 T cells could be further subdivided into cells expressing *KIR2DL4*, which is a gene associated with T cell exhaustion that is highly expressed in the CD8 T cell cluster 5 (expressing *TIGIT HAVCR2*) and in CD8 T_RM_ cluster 1. Expanded *ITGAE-*expressing CD8 T cells also upregulated the Th17 cytokine genes *IL17A* and *IL26* and are transcriptionally similar to *IL26*-expressing CD8 T cells previously reported in IBD, where IL-26 has been proposed to ameliorate colitis (Smillie et al. 2019; Bottois et al. 2020; Corridoni et al. 2020). Though not observed in our serum Luminex data from patients on anti-PD-1 and/or CTLA-4 therapy, increased peripheral IL-17 has been reported in patients with anti-CTLA-4-associated colitis and has been proposed as a therapeutic target of disease (Callahan MK S. Tandon et al. 2011; Postow, Sidlow, and Hellmann 2018). Clinical trial data, however, suggest that anti-IL17 therapy may worsen bowel inflammation in patients with pre-existing IBD and may be associated with an increased incidence of *de novo* IBD, raising the possibility that IL-17 may be protective in human colitis and, concerningly, that anti-IL-17 therapy may actually worsen human colitis (Hueber et al. 2012; Targan et al. 2016; Fauny et al. 2020; Anonymous 2015). Our data revealed marked diversification of CD8 T_RM_ cells in tissue in the setting of irColitis, which was not mirrored in blood, and raises the possibility that irColitis is caused by an expansion of CD8 T_RM_ populations with diverse epitope specificity.

In addition to CD8 T_RM_ cells, circulating CD8 T cells from blood are also likely to contribute to irColitis pathogenesis. Two CD8 T cell clusters enriched in irColitis, including clusters 3 (*EOMES*, *GZMK*, *KLRG1*) and 11 (*CX3CR1*), expressed genes found in circulating T cells (e.g., *S1PR1*, *S1PR5*, *CCR7*, *KLF2*, *SELL*) and shared TCRs with circulating CD8 T populations previously purported to be tissue recirculating and CX3CR1^+^ intravascular effector memory populations, respectively (Buggert et al. 2020; Gerlach et al. 2016). Neither population shared significant TCR clones with other tissue-derived CD8 T cell populations (including cycling cluster 4 cells) or displayed many differentially expressed genes between cases and controls, suggesting that these populations may be expanded in irColitis due to relocalization to inflamed tissue and not just cellular division. While it is unclear if these two populations (i.e., clusters 3 and 11) emigrated from tissue or blood during colitis, T_RM_ studies in the context of intestinal transplants clearly demonstrate that CD8 T_RM_ cells co-expressing *GZMK* and *KLRG1* (phenotypically similar to cluster 3 CD8 T co-expressing *EOMES* and *GZMK* in our data) are more frequently repopulated from blood than other T_RM_ populations and give rise to long-lived memory cells in the intestinal mucosa (Bartolomé-Casado et al. 2019; FitzPatrick et al. 2021). Furthermore, our cell-cell interaction analysis identified several putative receptor-ligand pairs that may mediate recruitment of these two populations to inflamed endothelium (*ITGAL/ITGB2-ICAM1*, *ITGAL/ITGB2-ICAM2, ITGA4/ITB7*-*MADCAM1*, *CX3CR1*-*CX3CL1*) and retention of CD8 T cluster 3 cells in tissue (*KLRG1-CDH1*, *CXCR3-CXCL9/10/11*). Clusters 3 and 11 both expressed cytotoxic transcriptional programs defined by *GZMK*, *GZMH*, *GZMB*, *GZMA,* and *PRF1,* though future work will need to define the exact contribution of these circulating populations to irColitis pathogenesis. These two populations were also notably more abundant in patients on dual PD-1/CTLA-4 inhibition (e.g. those more likely to develop colitis (Wolchok et al. 2017)) compared to those on anti-PD-1 monotherapy, suggesting that they may play a pathologic role in irColitis.

Our findings further support an active role for CD4 T cells and myeloid cells in irColitis with smaller contributions from B cells. irColitis patients showed marked tissue expansion of Tregs, including those with Th1-like gene signatures (*TBX21*, *CXCR3, IL12RB1/*2), and CD4 T cells expressing Th17 (*IL17A, IL17F, IL26*) and Th1 (*IFNG, CXCR3*) gene programs. Like CD8 T cells, CD4 T cells upregulated *CXCL13* in irColitis. CXCL13-expressing T cells have also been described across many different cancer types including colon cancer, where they comprise larger hubs that are relatively depleted of B cells but contain *CXCL10/11*-expressing myeloid and malignant cells and are predictive of immunotherapy responsiveness (Litchfield et al. 2021; Pelka et al. 2021). Thus, in the context of ICI therapy, *CXCL13* may serve as a homing signal for cell types other than *CXCR5* expressing B cells. Importantly, tissue B cells showed no abundance differences and few differentially expressed genes between irColitis patients and controls. A prior study (Das et al. 2018) reported that patients with IRAEs showed expanded peripheral CD21^Lo^ B cells and plasmablasts, which we did not observe in our data, potentially because the reported study included patients with diverse, extraintestinal IRAEs. We did, however, observe increased transitional B cells in the blood of irColitis patients, a finding also seen in patients with Lupus, where transitional B cells may be an important checkpoint for autoreactive B cells clones (Dieudonné et al. 2019). There has been at least one report of decreased IRAEs in the setting for B cell-depleting therapy (i.e. for lymphoma) (Risbjerg et al. 2020); however, given our observation of irColitis in a patient depleted of B cells and the few differences in tissue B cells seen in our data, B cell depletion is expected to have little impact on irColitis incidence. Unlike in IBD, we observed no skewing from IgA to IgG plasma B cells. We did, however, see marked differences in IgG receptor expression in myeloid cells marked by elevated ratios of activating (*FCGR1A*, *FCGR3A*) to inhibitory receptors (*FCGR2B*), which has been observed in ulcerative colitis and associated with NLRP3 inflammasome activation in myeloid cells (Castro-Dopico et al. 2019). Indeed, our data showed that myeloid cells from irColitis patients upregulated multiple NLRP3 inflammasome components (*CASP1*, *PYCARD*, *GSDMD*, *IL18*), which may drive the expansion of mononuclear phagocytes with an ISG^Hi^ phenotype (MP cluster 6) in active disease. While patients with irColitis displayed few differences in B cell subset abundance and gene expression, B cells may contribute to disease through crosstalk with myeloid cells expressing increased IgG activating receptors.

Importantly, we saw marked differences between tissue and blood immune cells from irColitis patients. While overlapping CD8 TCRs were observed between the tissue and blood of individual patients, the CD8 T cell programs that dominated tissue (e.g., *IL17A, IL26, CXCL13*) were not seen in blood CD8 T cells and secreted factors corresponding to these programs (e.g., *IL17A, IL17F, IFNG, IFNA, CXCL13, IL1A, IL1B, IL10, IL22, CCL3/4/5)* could not be readily detected in patient serum. This is likely because patients on immunotherapy have complex peripheral immune responses shaped by a combination of tumor burden, tumor responder vs non-responder status, and the presence of other IRAEs as seen in our patient cohort (Table S1). These findings underscore the importance of studying irColitis disease pathogenesis in tissue.

Colonic epithelial and mesenchymal cells from irColitis patients displayed a strong ISG signature that included genes implicated in immune cell chemotaxis, TCR signaling, angiogenesis, and epithelial absorptive and anti-microbial defense pathways. Our data suggest that ISG responses likely vary along the length of the epithelial crypt axis where canonical (e.g., *ISG15, ISG20, OAS1, GBP4, STAT1*) and non-canonical (e.g., *CD274*, *CXCL1*, MHC II) ISGs are most strongly induced in epithelial cells expressing top crypt genes (*SELENOP, CEACAM7*). Protein immunofluorescence microscopy confirmed that PD-L1 was preferentially expressed from top crypt epithelial cells closest to the intestinal lumen. These cells displayed striking downregulation of aquaporin genes (*AQP87, AQP8, AQP11*) and solute-carrier genes, including genes associated with IBD in GWAS studies (*SCL22A5*, *SLC22A23*) (Leung et al. 2006; Serrano León et al. 2014). These novel findings in irColitis explain absorptive defects that are likely contributing directly to diarrheal symptom morbidity observed in patients. These transcriptional absorptive defects were coupled with the upregulation of genes associated with apoptotic pathways (e.g.*, CASP8, TNRFSF10A, ZBP1*), a relative depletion of epithelial stem cells, and an increase in TA cells that collectively reflect increased epithelial turnover. Epithelial and mesenchymal cells from irColitis patients also upregulated genes implicated in neutrophil chemotaxis (e.g., *CXCL1, CXCL2, CXCL3, CXCL8*) and growth (*CSF3*), which provide a mechanism for the histopathologic finding of neutrophilic cryptitis observed in some patients with irColitis (Chen et al. 2017). One drawback of our study is that granulocytes, such as neutrophils, were absent from our immune datasets because they were not readily captured by 10X droplet-based chemistry (*data not shown*). We suspect that many of the differentially-expressed gene programs seen in the epithelium of irColitis patients – including those involved in immune signaling (MHC II genes, *CD274*), epithelial absorptive function (*AQP8*), and antimicrobial function (*NOS2, DUOX2*) are the result of indirect immune-mediated tissue damage and not the direct effect of ICI therapy as many of these changes have also been observed in IBD patients (Biton et al. 2018; Nakazawa et al. 2004; Grasberger et al. 2015; Ricanek et al. 2015; Kanai et al. 2003).

Strikingly, the colon mesenchymal compartment displayed a marked increase in endothelial abundance in irColitis patients. Unexpectedly, patients on dual PD-1/CTLA-4 inhibition had increased endothelial abundance compared to those on anti-PD-1 monotherapy, though it is unclear if this vascular remodeling is protective or pathogenic in irColitis. In fibroblasts we observed marked upregulation of *OSMR*, which is a putative IBD risk gene that comprises a pathway implicated in TNF-resistance and upregulated in the colonic mucosa of IBD patients (Smillie et al. 2019; Liu et al. 2015; West et al. 2017). Unlike in IBD, however, we did not see a significant expansion of inflammatory fibroblasts or a depletion of goblet cells (Smillie et al. 2019; Kinchen et al. 2018). This may reflect the early time point in disease that samples were collected (days to weeks after symptoms onset (**Fig. 1B**, **Table S1**), compared to most IBD scRNA-seq datasets where patients have had symptoms for months to years and may have developed mucosal fibrosis.

We have identified several feed-forward loops that likely sustain inflammation in patients with irColitis. These include (1) the CXCR3 chemokine system, which recent studies have shown is required in myeloid cells for anti-tumor immunity in mice following checkpoint blockade (Chow et al. 2019; House et al. 2020), (2) interferon-related signaling via the cGAS-STING pathway, and (3) the upregulation of TCR co-stimulatory receptors (e.g. *TNFRSF9/14/18/25*) and their respective ligands (e.g. *NECTIN2*). Interestingly, colon mucosal T cells from irColitis patients also upregulated TCR co-inhibitory receptor-ligand pairs and genes implicated in T cell exhaustion (*CTLA4-CD80/86, HAVCR2-CEACAM1, PDCD1-CD274/PDCD1LG2)*, cytotoxicity (*IFNG, GZMB, GZMA*), and proliferation (*MKI67*), suggesting the uncoupling of exhaustion and effector T cell gene programs in the presence of checkpoint inhibition. Whether T cell effector function in this context is activated or suppressed is likely a reflection of its local tissue microenvironment, including the presence of TCR receptor co-inhibitory and co-stimulatory receptor ligands.

By globally mapping ligand-receptor interaction networks between tissue and immune cells in irColitis, our results identified several putative, cell-specific mechanisms that could be therapeutically targeted to treat irColitis while helping maintain anti-tumor immunity. While steroids and anti-TNF monoclonal antibodies are first and second-line treatments of irColitis, respectively, long-term prospective trials are lacking to assess the impact of these treatments on anti-tumor immune responses, which are blunted by highly immunosuppressive therapy (Faje et al. 2018; Arbour et al. 2018). Unsurprisingly, the inhibition of many molecular pathways activated in irColitis, such as *CXCR3-CXCL9/10/11* are predicted to have detrimental anti-tumor responses (Chow et al. 2019; House et al. 2020). In contrast, the inhibition of T cell suppressive pathways active in both irColitis and anti-tumoral immunity, such as interactions between *PCSK9-LDLR*, are predicted to worsen colitis. PCSK9 binds LDLR and targets it for lysosomal degradation, thus preventing interacting LDLR and CD3 subunits from being recycled to the cell membrane (Benjannet et al. 2004; Lagace et al. 2006; Yuan et al. 2021). Thus upregulated PCSK9 from colon epithelium is predicted to have a T cell suppressive effect as observed in intratumoral immunity (Yang et al. 2016; Yuan et al. 2021), though it is unclear if this T cell suppressive pathway is also active in inflamed colonic mucosa.

The incidence of IRAEs is expected to increase in the years to come as combination therapy becomes standard of care and new immune-based treatments for tumors (e.g. targeted TIGIT, LAG-3, and TIM-3) are predicted to induce serious IRAEs, including irColitis. In defining the detailed interaction networks between immune and colon epithelial cells, this body of work has set the foundation for undertaking the challenging task of dissociating immunotherapy efficacy from toxicity. This study provides a comprehensive framework to dissect the pathogenic underpinnings of IRAEs and identify more tailored therapeutic candidates that could be used to mitigate tissue-specific immune damage while maintaining anti-tumor immunity.

## Data and Code Availability

Count matrices (sc/snRNA-Seq, TCR and BCR) and related data will be deposited in the GEO database (https://www.ncbi.nlm.nih.gov/geo/) and raw human sequencing data will be deposited in the controlled access repository DUOS (https://duos.broadinstitute.org/), upon publication. Source code for data analysis will be made available on GitHub and a permanent data repository upon publication.

## Acknowledgements

We are deeply grateful to all donors and their families. We acknowledge the contribution of Moshe Biton, Eugene Drokhlyansky, Ramnik Xavier, and Hacohen lab members for their feedback on experimental design and data interpretation. This work was supported by a NIAID grant T32AR007258 (to K.S.) and a NIDDK T32DK007191 (to M.F.T). This work was made possible by the generous support from the National Institute of Health Director’s New Innovator Award (DP2CA247831; to A.C.V.), the Massachusetts General Hospital Transformative Scholar in Medicine Award (to A.C.V.), the Damon Runyon-Rachleff Innovation Award (to A.C.V.), The Melanoma Research Alliance Young Investigator Award (2020A016475; to A.C.V.), the Kraft Foundation Award (to. K.R. and A.C.V.), the Arthur, Sandra, and Sarah Irving Fund for Gastrointestinal Immuno-Oncology (to N.H. and A.C.V.), the Spanish Society of Medical Oncology (SEOM) grant for a 2-year translational project at the MGH Cancer Center (to L.Z.), the Adelson Foundation (to G.M.B), and the American College of Gastroenterology Clinical Research Award (R01AG068390; to H.K.).

## Contributions

M.F.T., K.S., and A.C.V. conceived and led the study; M.F.T. and A.C.V. led experimental design; M.F.T. carried out experiments with assistance from K.M., J.T., P.S., M.N., A.T., and BY.A.; K.S. designed and performed computational analysis, with input and assistance from M.N., N.S., and S.R. M.F.T, L.T.N., J.H.C. designed and performed microscopy experiment, with input and assistance from K.H.X. and V.J.; T.E., K.P. provided input for sc/snRNA-Seq experiments, protocols and data interpretation; M.F.T., L.Z., C.J.P., T.S., R.G., E.P.Y.C., R.J.S., D.J., G.M.B., H. K., M.D. and K.L.R. provided clinical expertise, coordinated and performed sample acquisition and/or administrative coordination; N.H. contributed to biological expertise, study design, and advice; B.L. contributed to computational expertise and advice; A.C.V. managed and supervised the study; M.F.T., K.S., A.C.V. wrote the manuscript, with input from all authors.

## Supplemental Figure Legends

**Figure S1.**
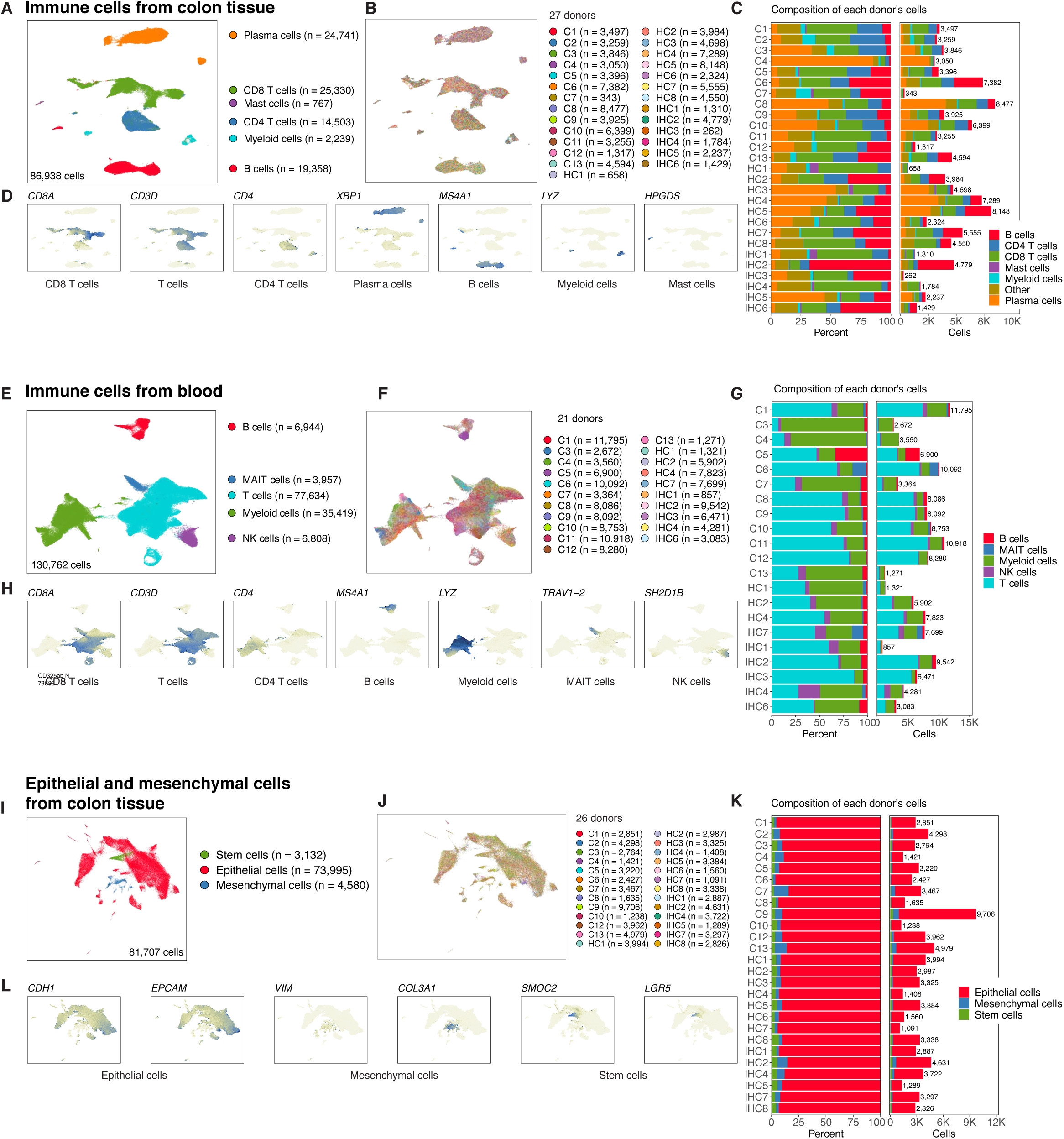
Major cell lineages and donor cell type composition for immune cells from colon tissue, immune cells from blood, and epithelial and mesenchymal cells from colon tissue. **A, E, I.** Major cell lineages depicted with different colors in UMAP embeddings of (**A**) immune cells from colon tissue, (**E**) immune cells from blood, and (**I**) epithelial and mesenchymal cells from colon tissue. **B, F, J.** Donors depicted with different colors in UMAP embeddings of (**B**) immune cells from colon tissue, (**F**) immune cells from blood, and (**J**) epithelial and mesenchymal cells from colon tissue. **C, G, K.** Donor cell type composition for (**C**) immune cells from colon tissue, (**G**) immune cells from blood, and (**K**) epithelial and mesenchymal cells from colon tissue. **D, H, L.** Selected genes that indicate major cell types in UMAP embeddings of (**D**) immune cells from colon tissue, (**H**) immune cells from blood, and (**L**) epithelial and mesenchymal cells from colon tissue.

**Figure S2.**
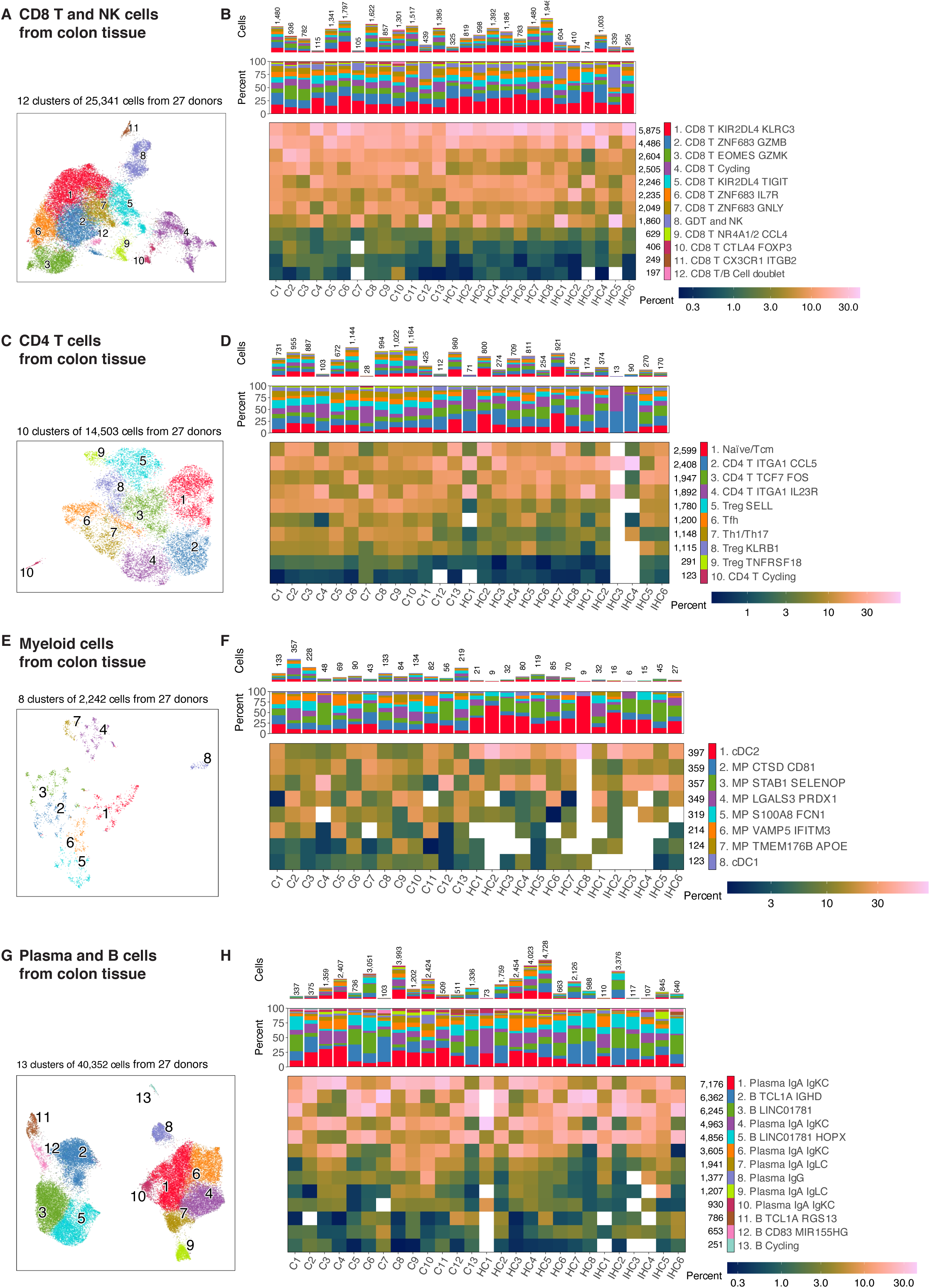
Detailed cell type composition of each donor for immune cells from colon tissue. **A, C, E, G.** Cell clusters depicted with different colors in UMAP embeddings of (**A**) CD8 T/GDT/NK cells, (**C**) CD4 T cells, (**E**) myeloid cells, (**G**) B cells from colon tissue. **B, D, F, H.** Detailed composition of each patient across cell clusters for (**B**) CD8 T/GDT/NK cells, (**D**) CD4 T cells, (**F**) myeloid cells, (**H**) B cells from colon tissue. Heatmap color indicates percent of patient’s cells assigned to each cell cluster.

**Figure S3.**
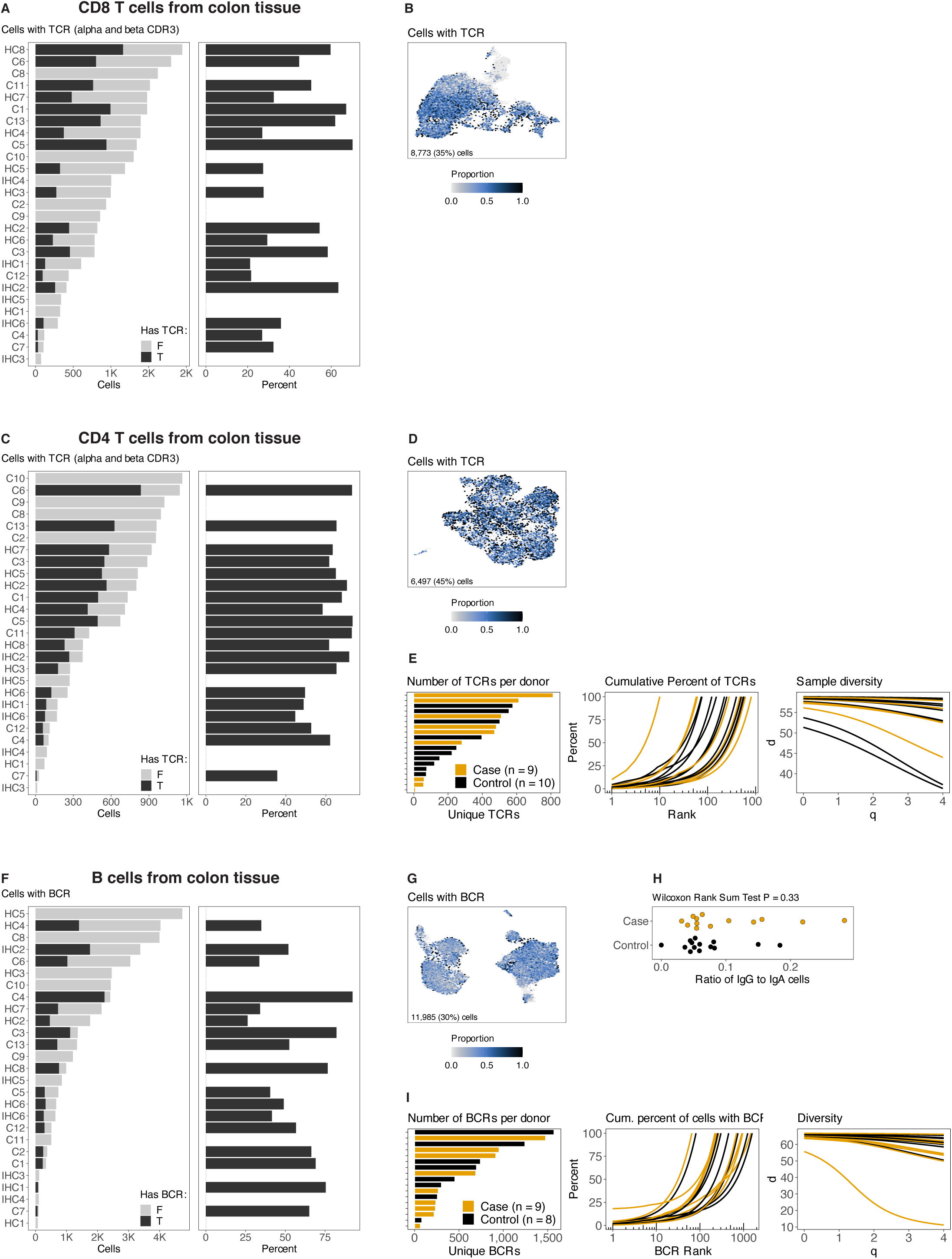
TCR and BCR detection and diversity analysis for immune cells from colon tissue. **A, C, F.** Number of cells from each donor for (**A**) CD8 T cells, (**C**) CD4 T cells, (**F**) B cells, with color indicating whether or not the T cell has a TCR or the B cell has a BCR. **B, D, G.** UMAP embedding with color depicting cells with TCR or BCR for (**B**) CD8 T cells, (**D**) CD4 T cells, (**G**) B cells from colon tissue. **E, I**. TCR analysis of (**E**) CD4 T cells and BCR analysis of (**I**) B cells. Bars indicate the number of unique clones per patient. Middle panel shows one line for each patient, with y-axis indicating cumulative percent of cells with the top N unique TCR clones (or BCR clones). Right-most panel shows the Hill diversity index. **H.** Proportion of IgG and IgA plasma B cells from each donor. Wilcoxon Rank Sum Test p-value = 0.33 for a difference between cases and controls.

**Figure S4.**
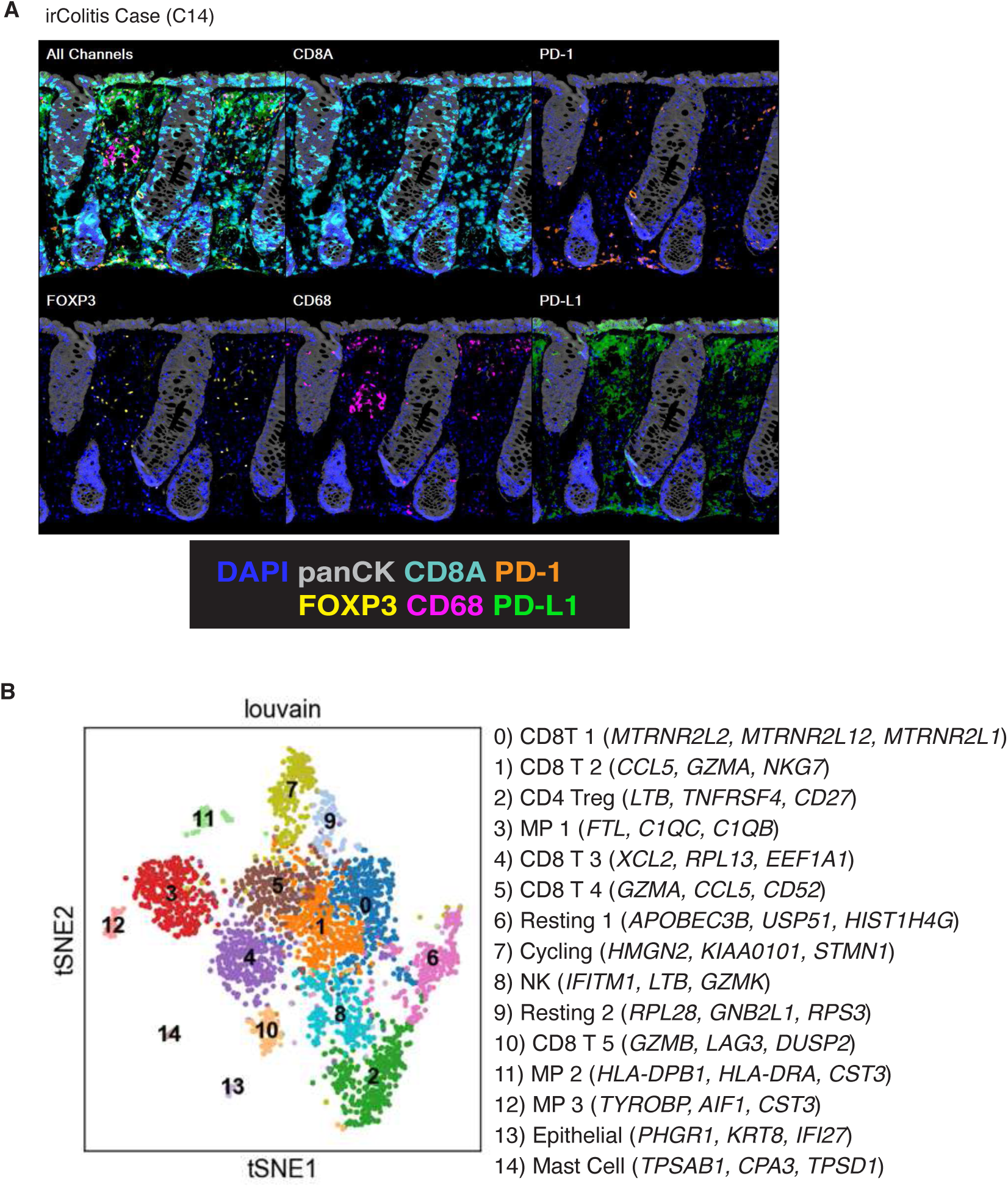
irColitis in a B cell-depleted patient receiving ICI therapy for lymphoma. **A.** Results from multispectral immunofluorescence staining of fixed colon mucosal tissue from C14* with a 7-color panel that included DAPI (blue), panCK (grey), CD8A (aqua), PD-1 (orange), FOXP3 (yellow), CD68 (pink), and PD-L1 (green). **B.** tSNE-embedding of 3,295 CD45^+^-sorted cells from a patient depleted of B cells. Cell cluster identity with top three AUC genes are shown in parentheses.

**Figure S5.**
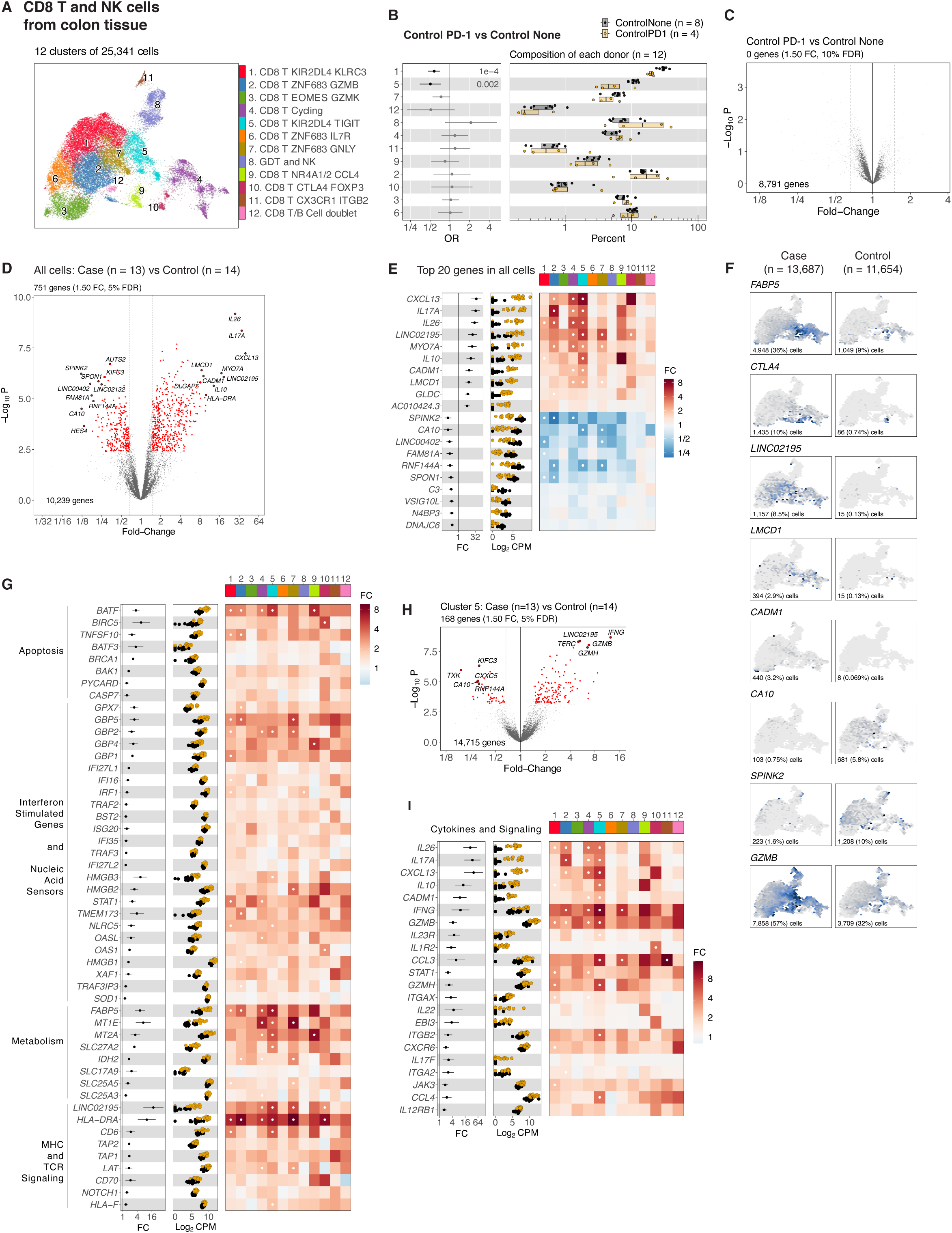
Detailed analysis of CD8 T, GDT, and NK cells from colon tissue. **A.** Cell clusters are depicted with different colors in UMAP embedding. **B.** Left panel shows logistic regression odds ratio (OR) for differential abundance of cells from Controls on PD-1 within each cell cluster. Right panel shows one dot for each patient, indicating the percent of the patient’s cells assigned to each cluster. **C.** Volcano of pseudobulk differential gene expression for all cells, for the contrast of controls on PD-1 versus Controls without therapy. The x-axis indicates the fold-change and y-axis indicates the negative log10 *p*-value reported by limma. **D, H.** Volcano of differential expression for pseudobulk expression with (**D**) all cells or with (**H**) cells in cluster 5, for the contrast of Cases versus Controls The x-axis indicates the fold-change and y-axis indicates negative log10 p-value reported by limma. **E, G, I.** Fold-changes, pseudobulk differential gene expression (log2CPM), and a heatmap with color indicating fold-change between cases and controls for each cell cluster. **F.** Gene expression displayed in UMAP embeddings, stratified by irColitis case and control cells. Number and percent of cells with expression is shown in each panel.

**Figure S6.**
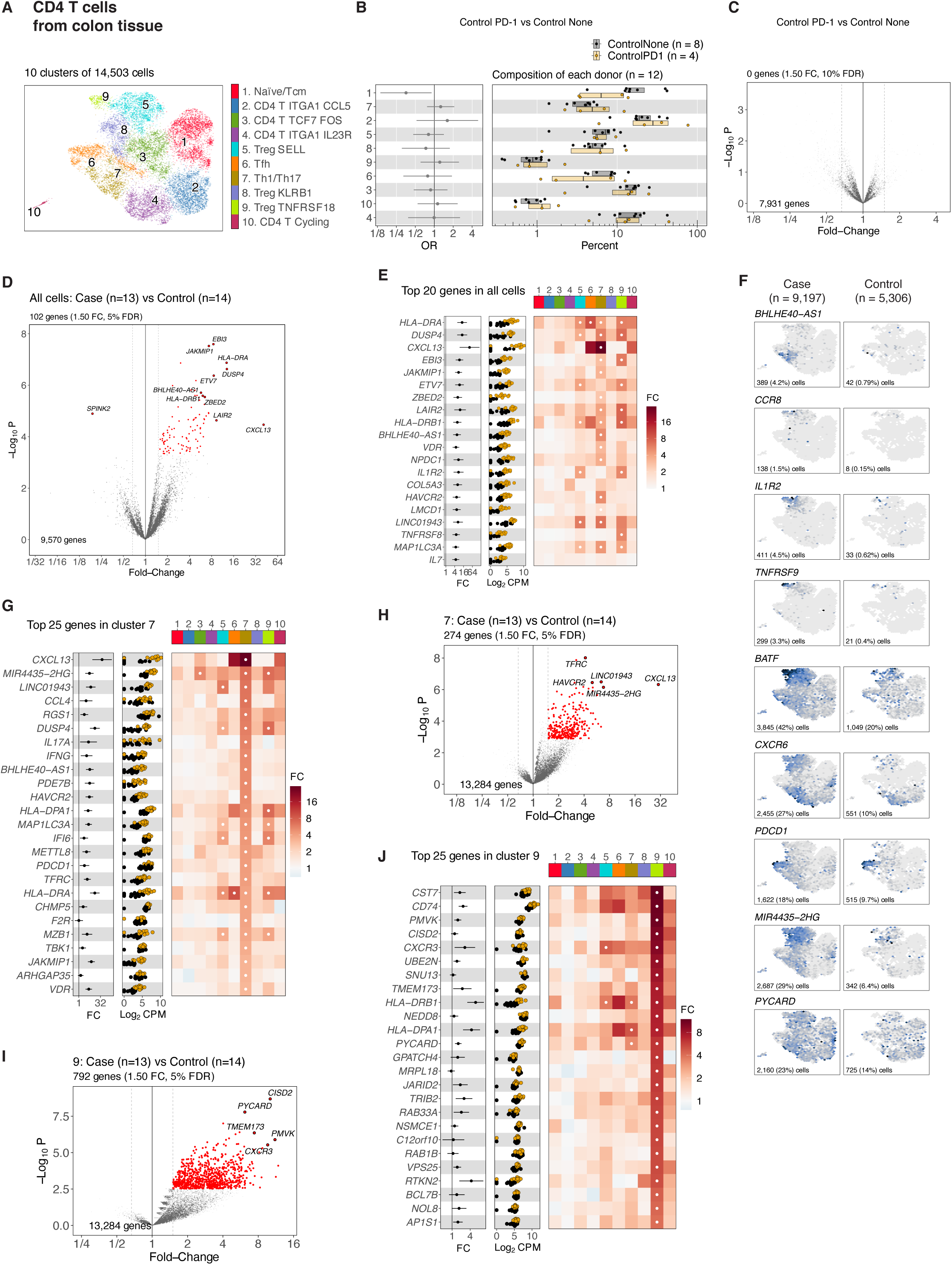
Detailed analysis of CD4 T cells from colon tissue. **A.** Cell clusters are depicted with different colors in UMAP embedding. **B.** Left panel shows logistic regression odds ratio (OR) for differential abundance of cells from controls on PD-1 within each cell cluster. Right panel shows one dot for each patient, indicating the percent of the patient’s cells assigned to each cluster. **C.** Volcano of pseudobulk differential gene expression with all cells, for the contrast of controls on PD-1 versus controls without therapy. The x-axis indicates the fold-change and y-axis indicates the negative log10 p-value reported by limma. **D, H, I.** Volcano of differential pseudobulk expression with (**D**) all cells or (**H**) cells in cluster 7 or (**I**) cells in cluster 9, for the contrast of irColitis cases versus controls. The x-axis indicates the fold-change and y-axis indicates negative log10 *p*-value reported by limma. **E, G, J.** Fold-changes, pseudobulk differential gene expression (log2CPM), and a heatmap with color indicating fold-change between cases and controls for each cell cluster. **F.** Gene expression displayed in UMAP embeddings, stratified by irColitis case and control cells. Number and percent of cells with expression is shown in each panel.

**Figure S7.**
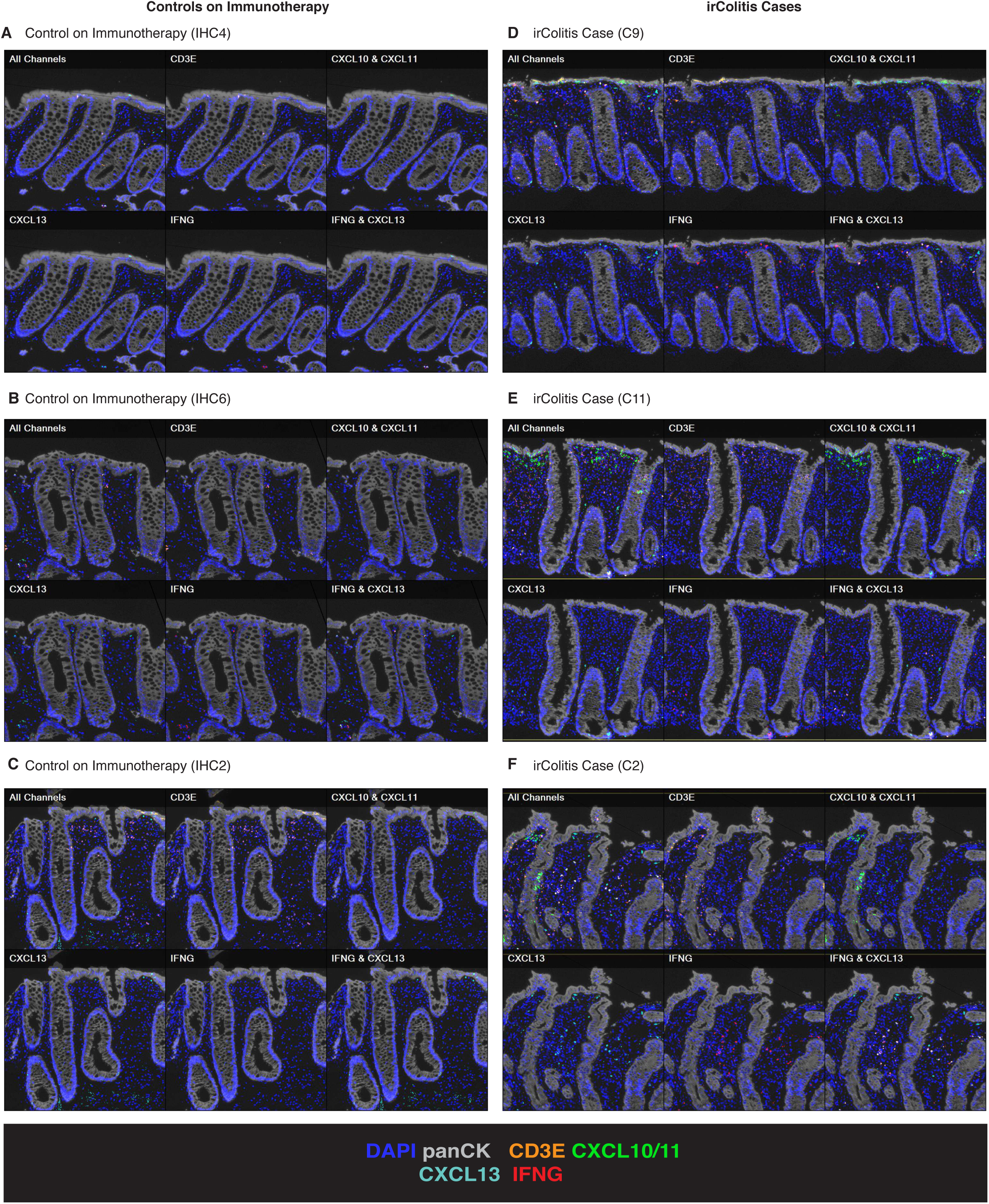
Multispectral microscopy images of irColitis cases and controls for the detection of transcripts for *CD3E*, *CXCL10/11, CXCL13*, and *IFNG*. **A, B, C, D, E, F.** Results from staining colon mucosal tissue from (**A, B, C**) controls and (**D, E, F**) irColitis cases with a 6-color multispectral fluorescence panel depicting the RNA expression level of *CD3E* (orange), *CXCL13* (aqua), *IFNG* (red), and *CXCL10/11* (green), protein immunofluorescence of panCK (grey), and DAPI (blue).

**Figure S8.**
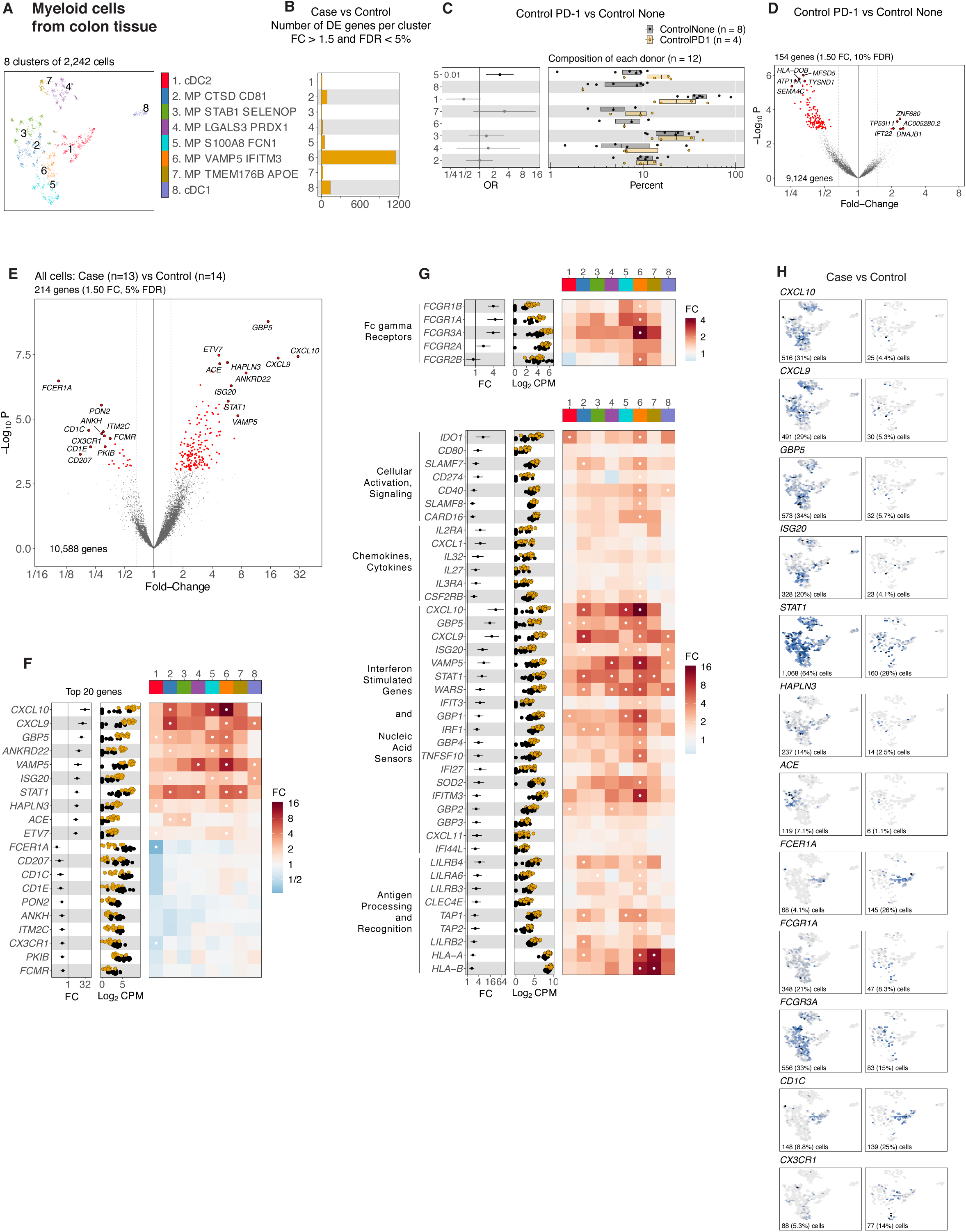
Detailed analysis of Myeloid cells from colon tissue. **A.** Cell clusters are depicted with different colors in UMAP embedding. **B.** Number of differentially expressed genes for each cell cluster (fold-change greater than 1.5, false discovery rate less than 5%). **C.** Left panel shows logistic regression odds ratio (OR) for differential abundance of cells from controls on PD-1 within each cell cluster. Right panel shows one dot for each patient, indicating the percent of the patient’s cells assigned to each cluster. **D.** Volcano of pseudobulk differential gene expression with all cells, for the contrast of controls on PD-1 versus controls without therapy. The x-axis indicates the fold-change and y-axis indicates the negative log10 p-value reported by limma. **E.** Volcano of pseudobulk differential gene expression with all cells, for the contrast of irColitis cases versus controls. The x-axis indicates the fold-change and y-axis indicates negative log10 p-value reported by limma. **F, G.** Fold-changes, pseudobulk differential gene expression (log2CPM), and a heatmap with color indicating fold-change between irColitis cases and controls for each cell cluster. **H.** Gene expression displayed in UMAP embeddings, stratified by irColtiis case and control cells. Number and percent of cells with expression is shown in each panel.

**Figure S9.**
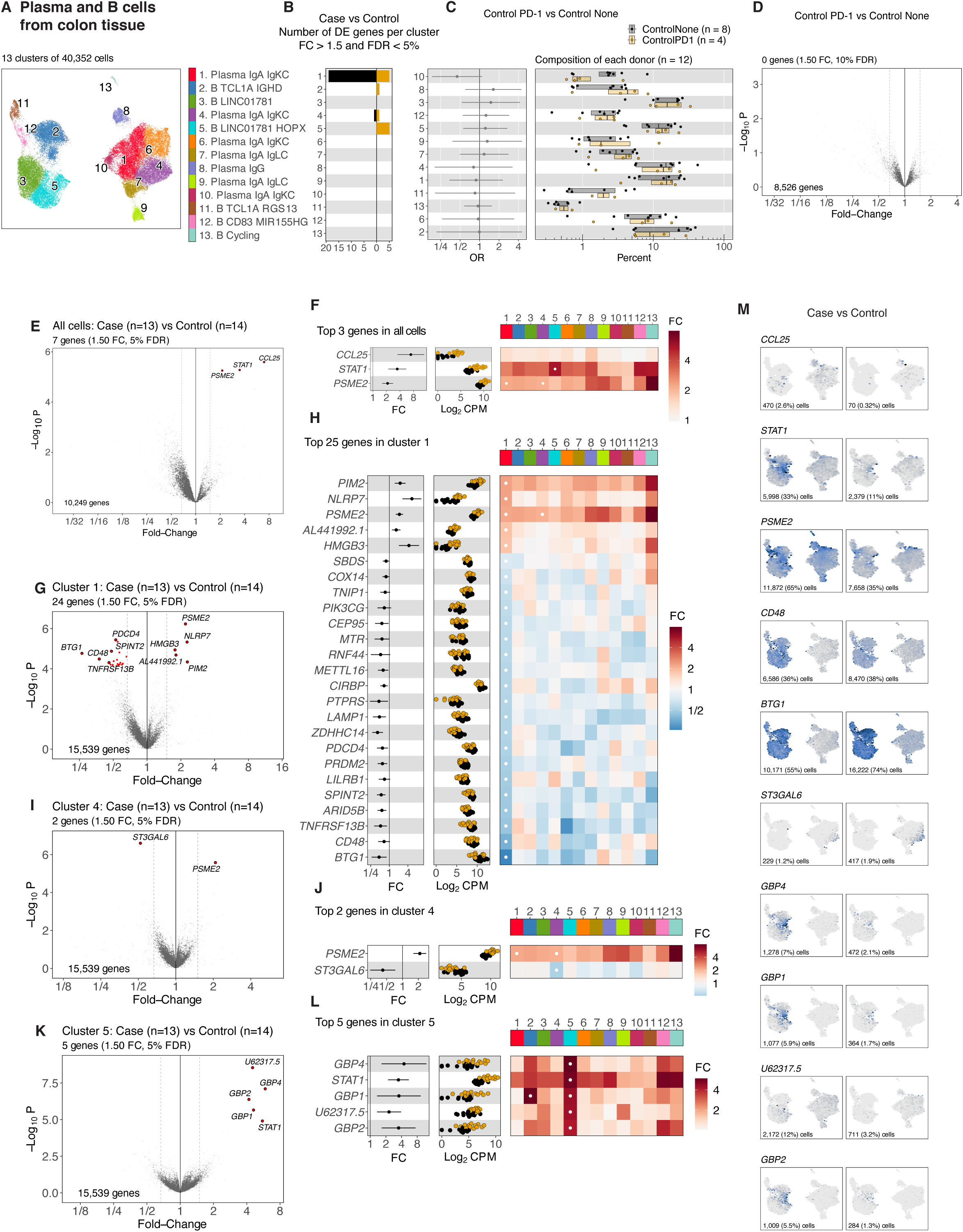
Detailed analysis of B cells from colon tissue. **A.** Cell clusters are depicted with different colors in UMAP embedding. **B.** Number of differentially expressed genes for each cell cluster (fold-change greater than 1.5, false discovery rate less than 5%). **C.** Left panel shows logistic regression odds ratio (OR) for differential abundance of cells from Controls on PD-1 within each cell cluster. Right panel shows one dot for each patient, indicating the percent of the patient’s cells assigned to each cluster. **D.** Volcano of pseudobulk differential gene with all cells, for the contrast of controls on PD-1 versus controls without therapy. The x-axis indicates the fold-change and y-axis indicates the negative log10 p-value reported by limma. **E, G, I, K.** Volcano of differential expression for pseudobulk expression with (**E**) all cells or (**G**) cluster 1 cells or (**I**) cluster 4 cells or (**K**) cluster 5 cells, for the contrast of irColitis cases versus controls. The x-axis indicates the fold-change and y-axis indicates negative log10 p-value reported by limma. **F, H, J, L.** Fold-changes, pseudobulk differential gene expression (log2CPM), and a heatmap with color indicating fold-change between cases and controls for each cell cluster. **M.** Gene expression displayed in UMAP embeddings, stratified by irColitis case and control cells. Number and percent of cells with expression is shown in each panel.

**Figure S10.**
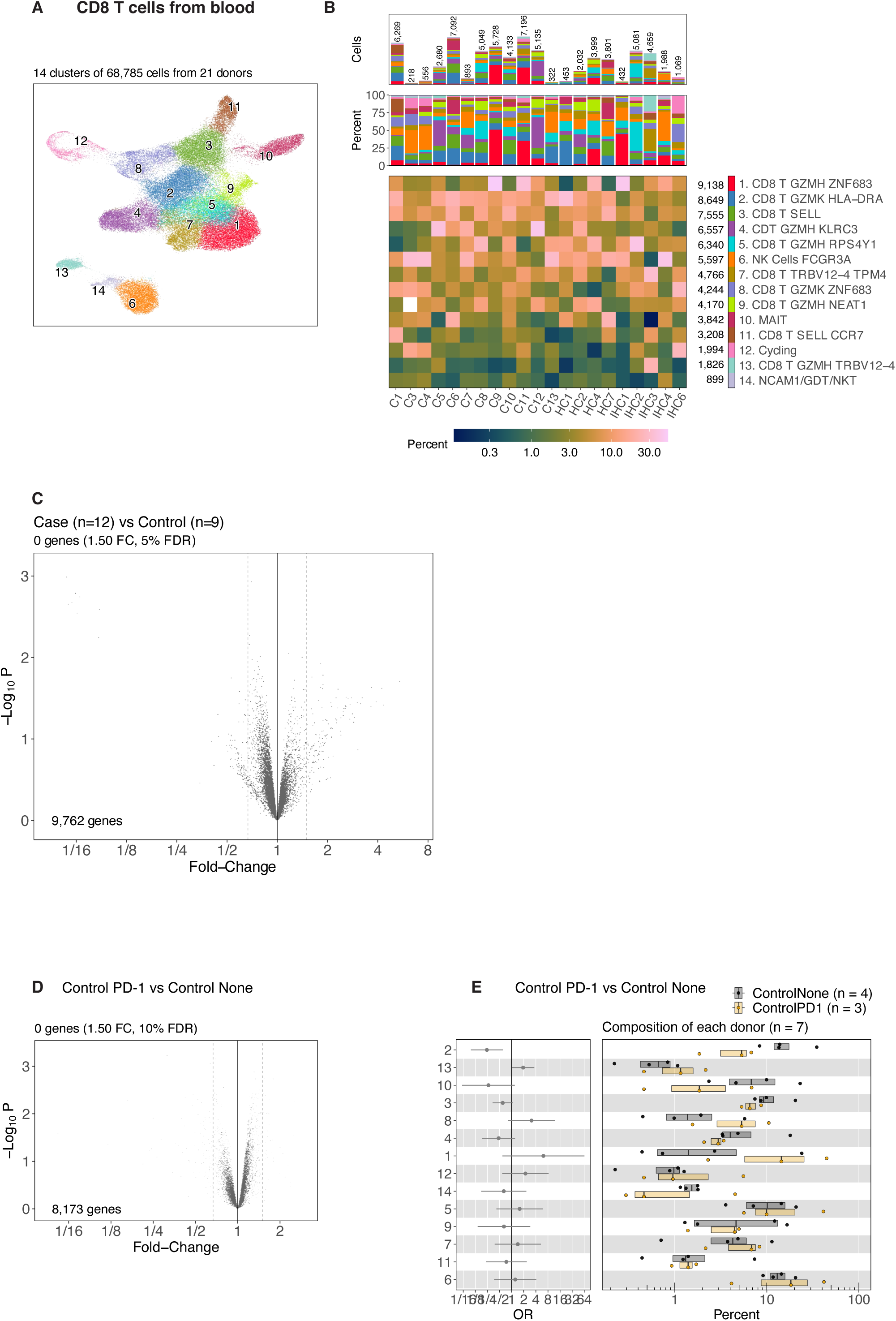
Detailed analysis of CD8 T/GDT/NK/MAIT cells from blood. **A.** Cell clusters are depicted with different colors in UMAP embedding. **B.** Detailed composition of each patient across cell clusters. Heatmap color indicates percent of patient’s cells assigned to each cell cluster. **C, D.** Volcano of pseudobulk differential gene expression with all cells for the contrast of (**C**) irColitis cases versus controls or (**D**) controls on PD-1 versus Controls on no therapy. The x-axis indicates the fold-change and y-axis indicates the negative log10 p-value reported by limma. **E.** Left panel shows logistic regression odds ratio (OR) for differential abundance of cells from controls on PD-1 versus Controls on no therapy within each cell cluster. Right panel shows one dot for each patient, indicating the percent of the patient’s cells assigned to each cluster.

**Figure S11.**
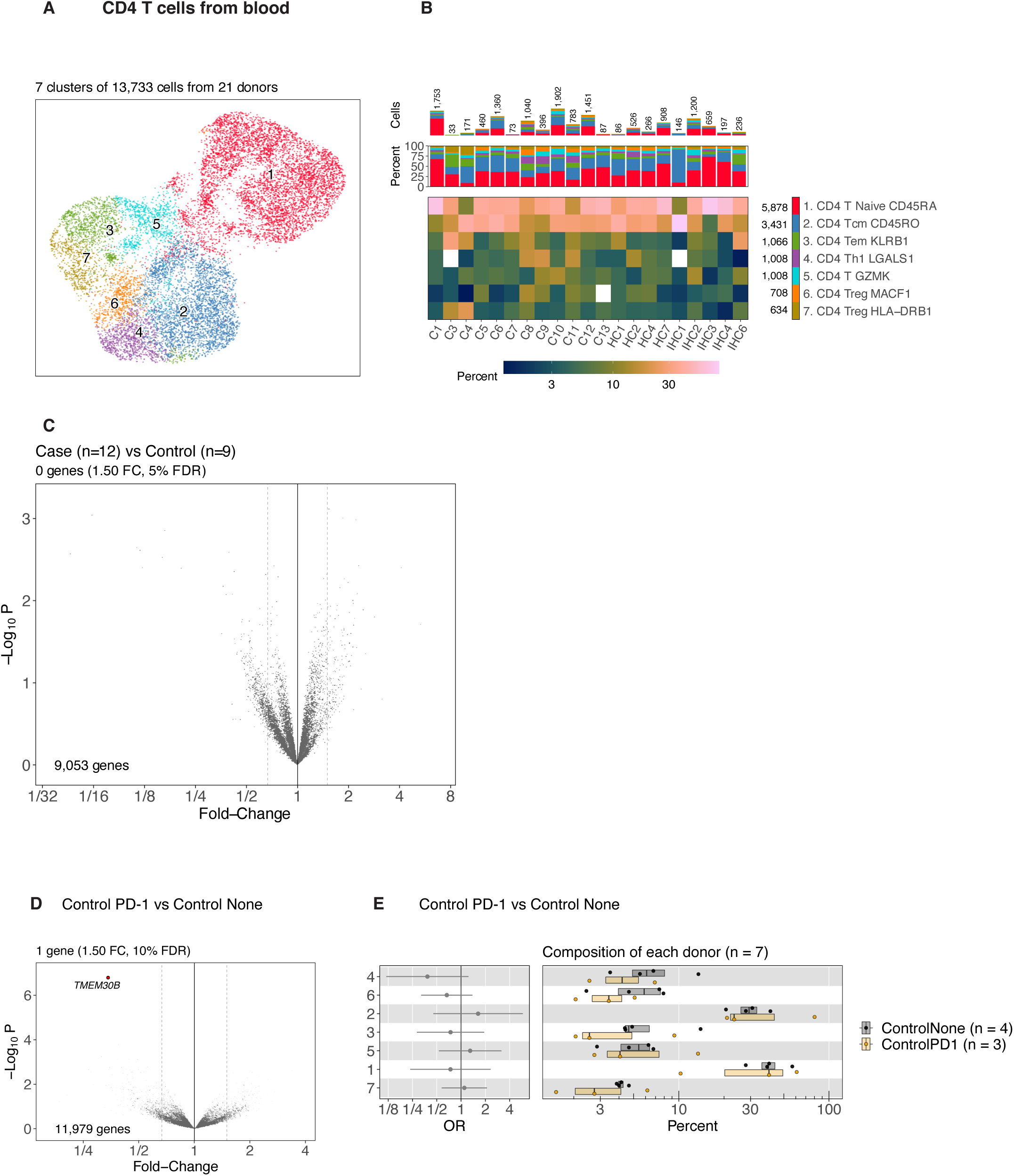
Detailed analysis of CD4 T cells from blood. **A.** Cell clusters are depicted with different colors in UMAP embedding. **B.** Detailed composition of each patient across cell clusters. Heatmap color indicates percent of patient’s cells assigned to each cell cluster. **C, D.** Volcano of pseudobulk differential gene expression with all cells for the contrast of (**C**) irColitis cases versus controls or (**D**) controls on PD-1 versus Controls on no therapy. The x-axis indicates the fold-change and y-axis indicates the negative log10 p-value reported by limma. **E.** Left panel shows logistic regression odds ratio (OR) for differential abundance of cells from controls on PD-1 versus Controls on no therapy within each cell cluster. Right panel shows one dot for each patient, indicating the percent of the patient’s cells assigned to each cluster.

**Figure S12.**
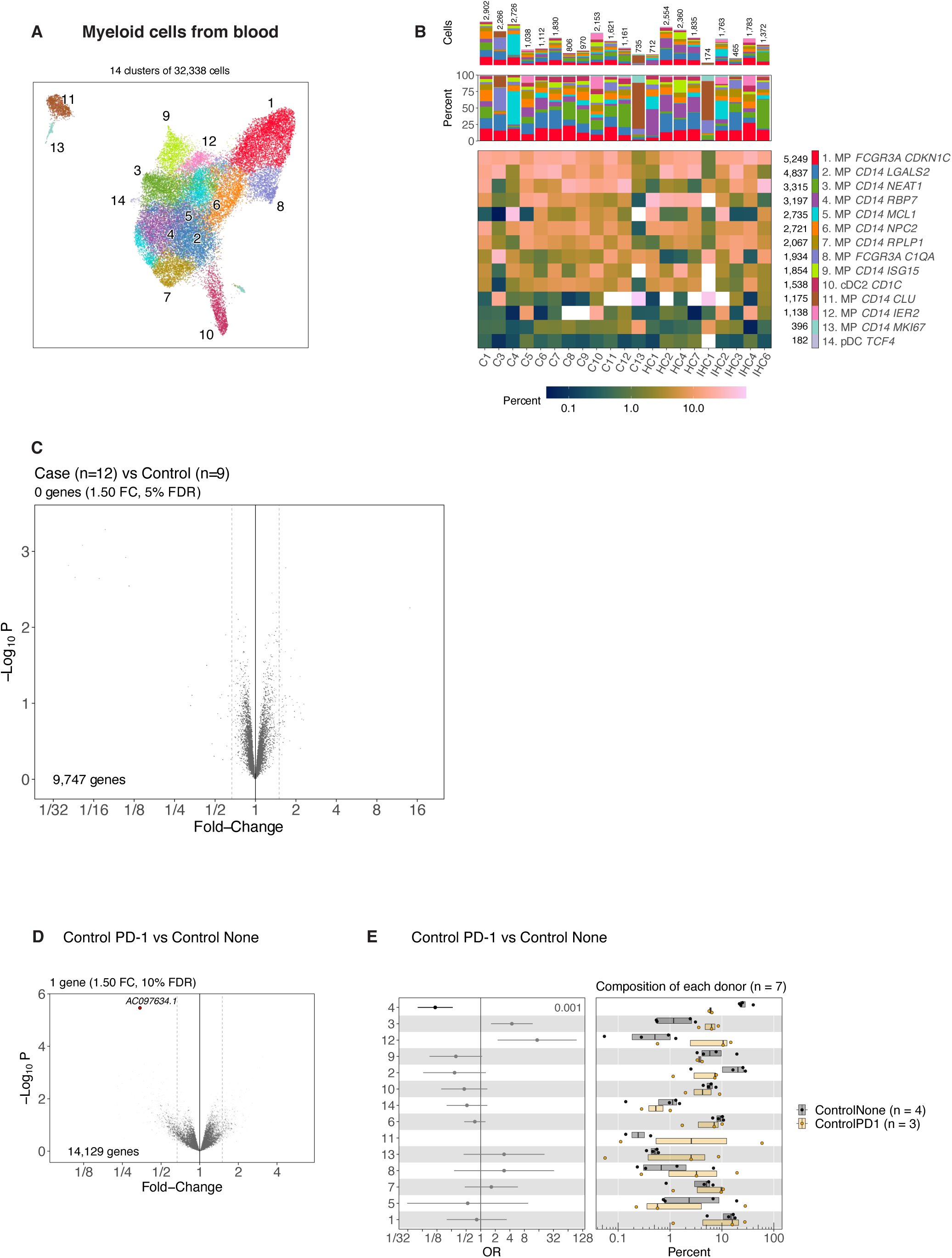
Detailed analysis of Myeloid cells from blood. **A.** Cell clusters are depicted with different colors in UMAP embedding. **B.** Detailed composition of each patient across cell clusters. Heatmap color indicates percent of patient’s cells assigned to each cell cluster. **C, D.** Volcano of pseudobulk differential gene expression with all cells for the contrast of (**C**) irColitis cases versus controls or (**D**) controls on PD-1 versus Controls on no therapy. The x-axis indicates the fold-change and y-axis indicates the negative log10 p-value reported by limma. **E.** Left panel shows logistic regression odds ratio (OR) for differential abundance of cells from controls on PD-1 versus Controls on no therapy within each cell cluster. Right panel shows one dot for each patient, indicating the percent of the patient’s cells assigned to each cluster.

**Figure S13.**
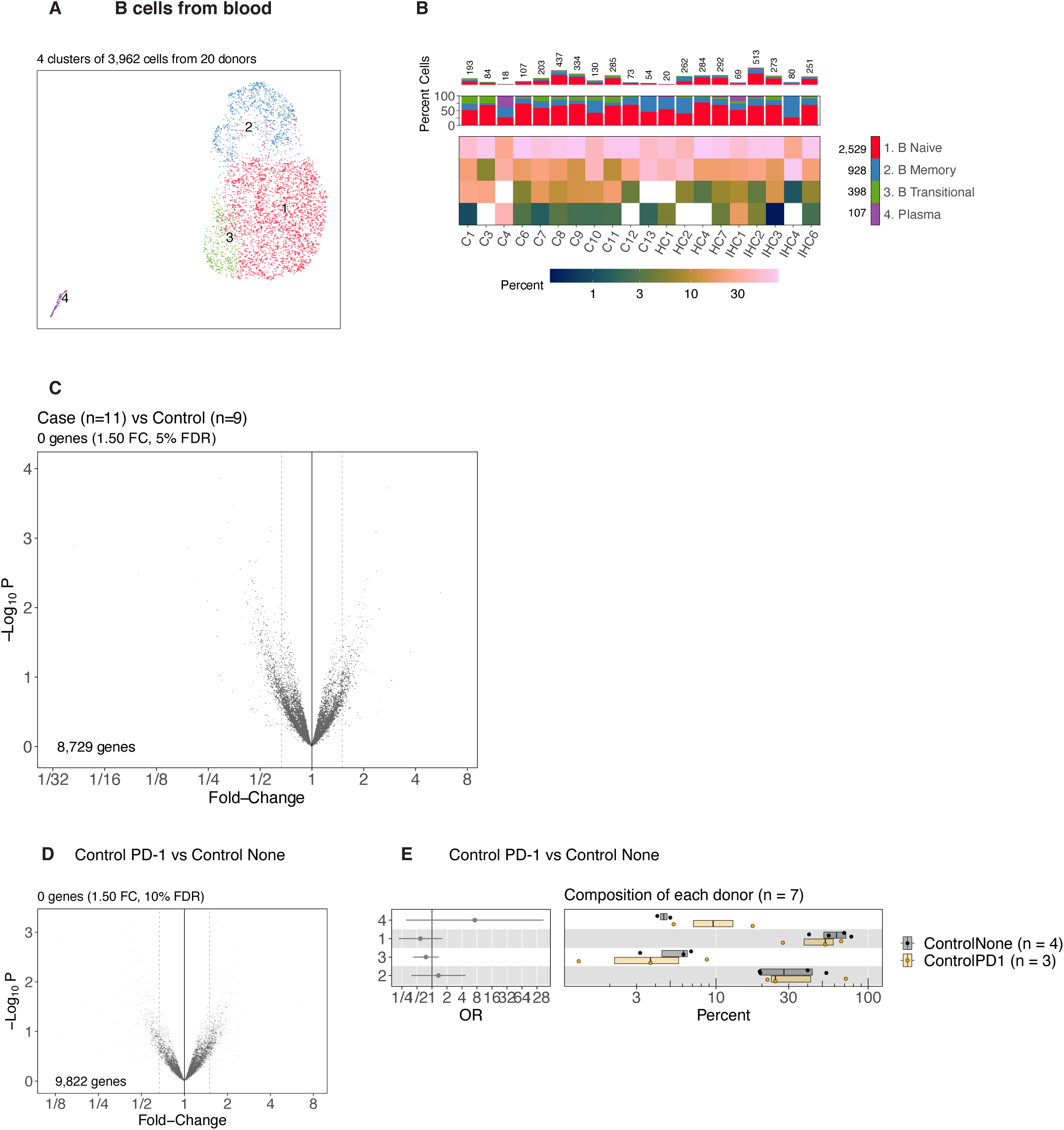
Detailed analysis of B cells from blood. **A.** Cell clusters are depicted with different colors in UMAP embedding. **B.** Detailed composition of each patient across cell clusters. Heatmap color indicates percent of patient’s cells assigned to each cell cluster. **C, D.** Volcano of pseudobulk differential gene expression with all cells for the contrast of (**C**) irColitis cases versus controls or (**D**) controls on PD-1 versus Controls on no therapy. The x-axis indicates the fold-change and y-axis indicates the negative log10 p-value reported by limma. **E.** Left panel shows logistic regression odds ratio (OR) for differential abundance of cells from controls on PD-1 versus Controls on no therapy within each cell cluster. Right panel shows one dot for each patient, indicating the percent of the patient’s cells assigned to each cluster.

**Figure S14.**
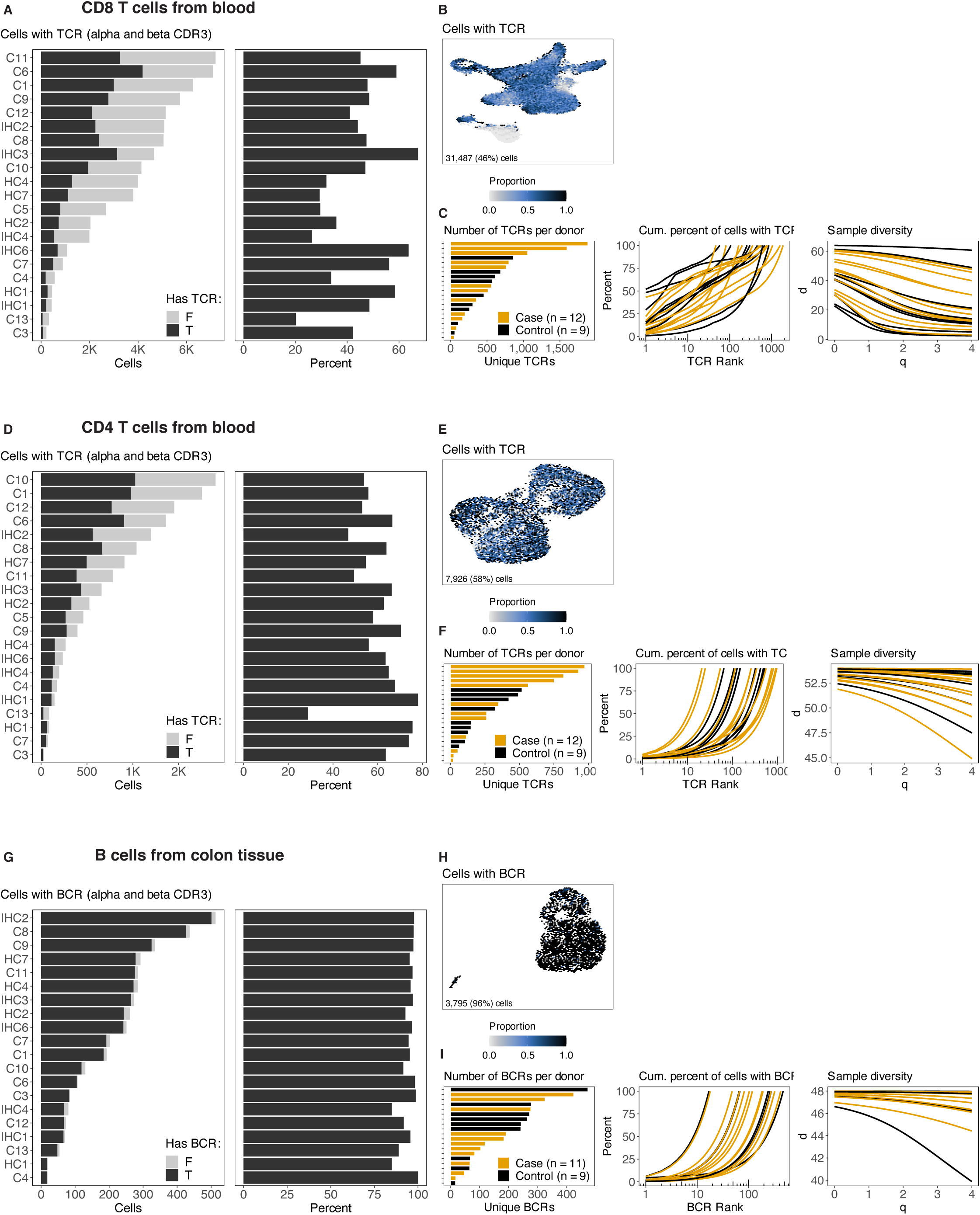
TCR and BCR detection and diversity analysis for immune cells from blood. **A, D, G.** Number of cells from each donor for (**A**) CD8 T cells, (**D**) CD4 T cells, (**G**) B cells, with color indicating whether or not the T cell has a TCR or the B cell has a BCR. **B, E, H.** UMAP embedding with color depicting cells with TCR or BCR for (**B**) CD8 T cells, (**E**) CD4 T cells, (**H**) B cells from colon tissue. **C, F, I**. TCR analysis of (**C**) CD8 T cells, (**F**) CD4 T cells, and (**I**) BCR analysis of B cells. Bars indicate the number of unique clones per patient. Middle panel shows one line for each patient, with y-axis indicating cumulative percent of cells with the top N unique TCR clones (or BCR clones). Right-most panel shows the Hill diversity index.

**Figure S15.**
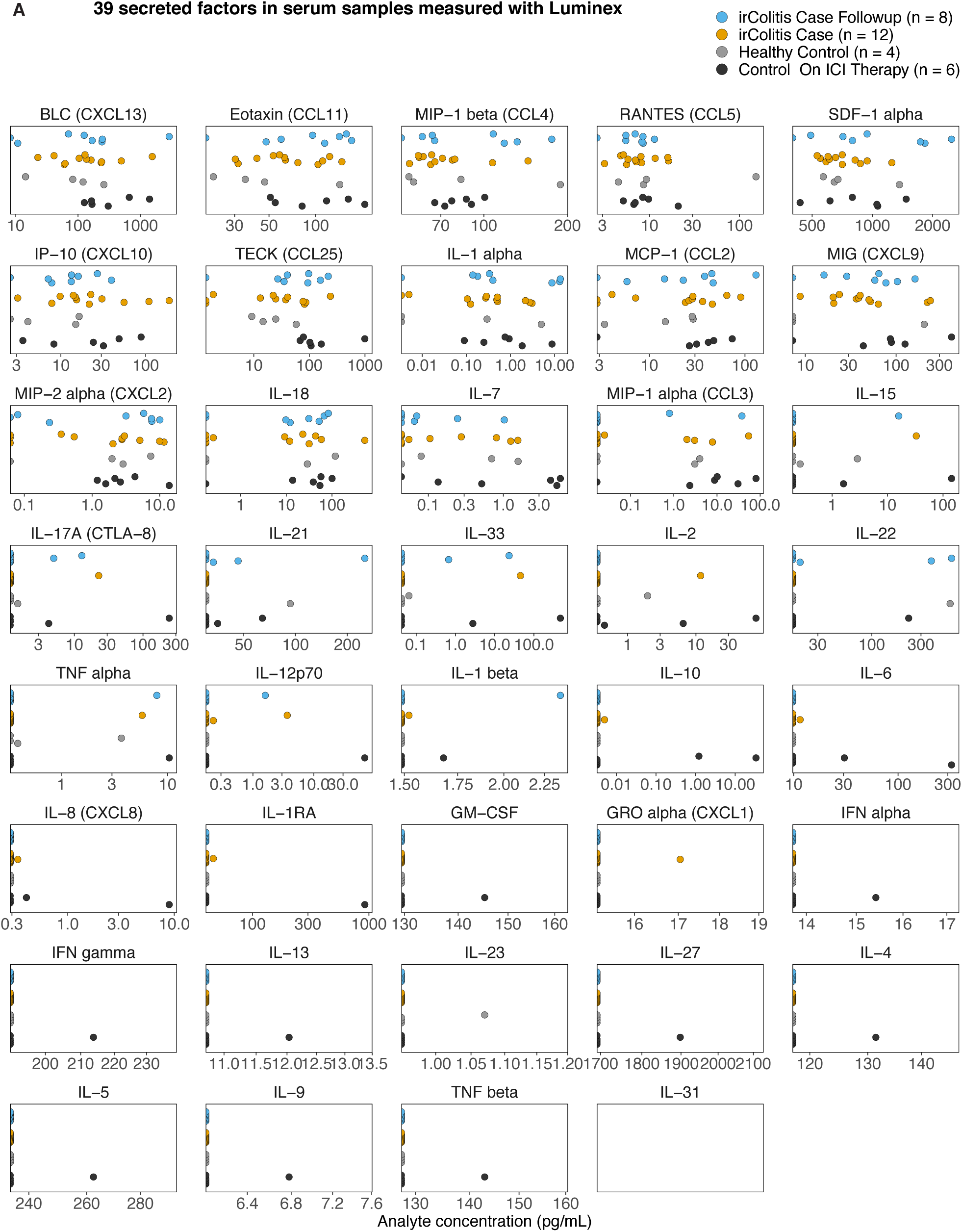
irColitis case vs control analysis of 39 secreted factors in serum blood samples measured with Luminex. Concentration of 39 secreted factors (pg/mL) for 30 patient samples in four categories (irColitis Case Followup, irColitis Case, Healthy Control, Control on ICI therapy).

**Figure S16.**
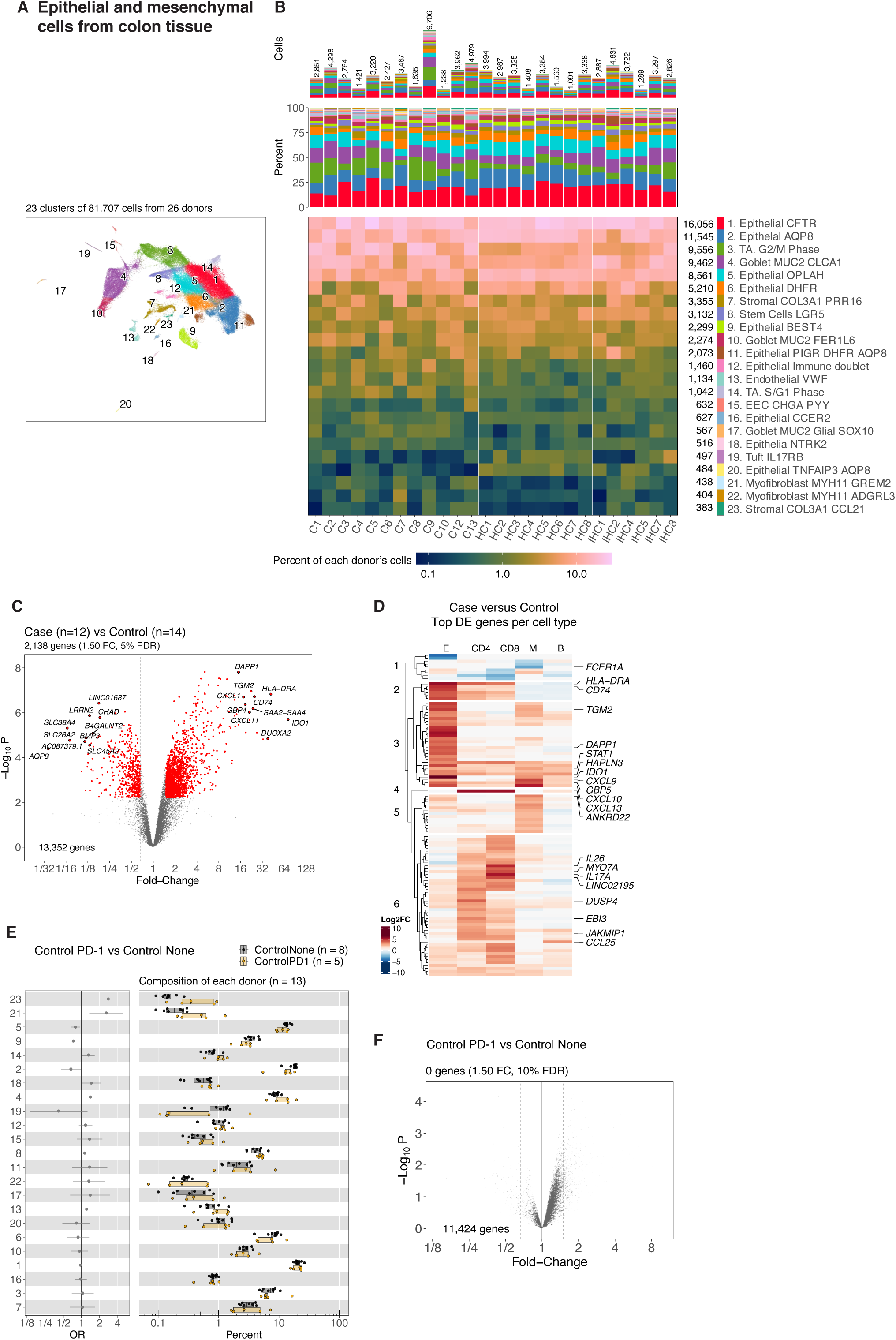
Detailed analysis of epithelial and mesenchymal cells from tissue. **A.** Cell clusters are depicted with different colors in UMAP embedding. **B.** Detailed composition of each patient across cell clusters. Heatmap color indicates percent of patient’s cells assigned to each cell cluster. **C, F.** Volcano of pseudobulk differential expression with all cells for the contrast of (**C**) Cases versus Controls or (**F**) Controls on PD-1 versus Controls on no therapy. The x-axis indicates the fold-change and y-axis indicates the negative log10 p-value reported by limma. **D.** Top differentially expressed genes for each major cell lineage. Color indicates the log2 fold-change between cases and controls for each gene and for each major cell lineage. **E.** Left panel shows logistic regression odds ratio (OR) for differential abundance of cells from Controls on PD-1 versus Controls on no therapy within each cell cluster. Right panel shows one dot for each patient, indicating the percent of the patient’s cells assigned to each cluster.

**Figure S17.**
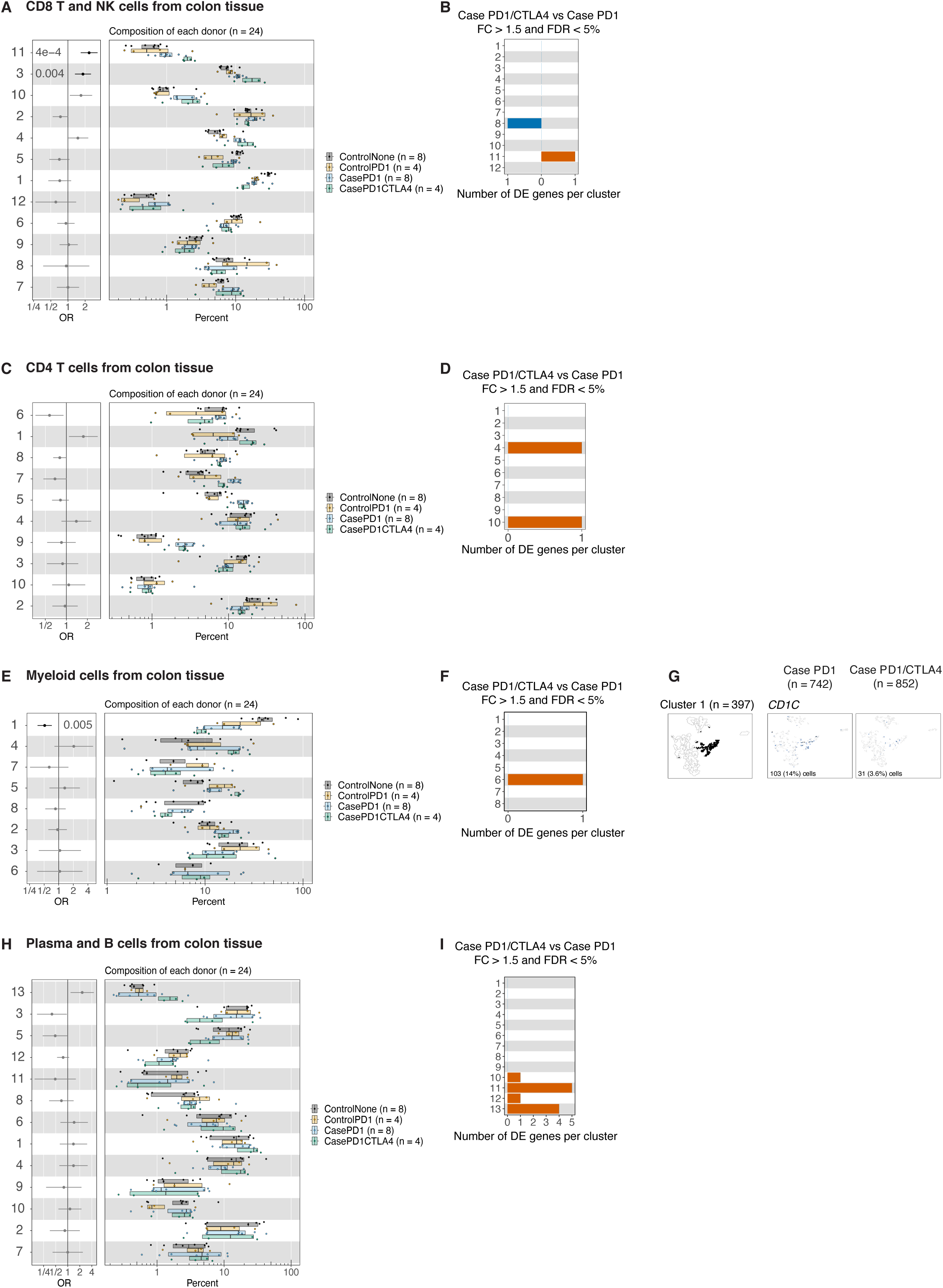
Cell type abundance and differential gene expression analysis of colon tissue immune cells from patients on anti-PD-1 vs anti-PD-1/CTLA-4 therapy. **A, C, E, H**. Left panel shows logistic regression odds ratio (OR) for differential abundance of cells from Cases on PD-1/CTLA-4 versus Cases on PD-1 within each cell cluster from (**A**) CD8 T/GDT/NK cells, (**C**) CD4 T cells, (**E**) Myeloid cells, or (**H**) B cells. Right panel shows one dot for each patient, indicating the percent of the patient’s cells assigned to each cluster. Likelihood ratio test *p-*values are shown for clusters with false discovery rate less than 5%. **B, D, F, I.** Number of differentially expressed genes for each cell cluster (fold-change greater than 1.5, false discovery rate less than 5%) from (**B**) CD8 T/GDT/NK cells, (**D**) CD4 T cells, (**F**) Myeloid cells, or (**I**) B cells. **G.** Cells labeled by assignment to cluster 1 on a UMAP embedding. Gene expression (log2CPM) for *CD1C* (the one differentially expressed gene for myeloid cells) across cells from Cases on PD-1 or PD-1/CTLA-4.

**Figure S18.**
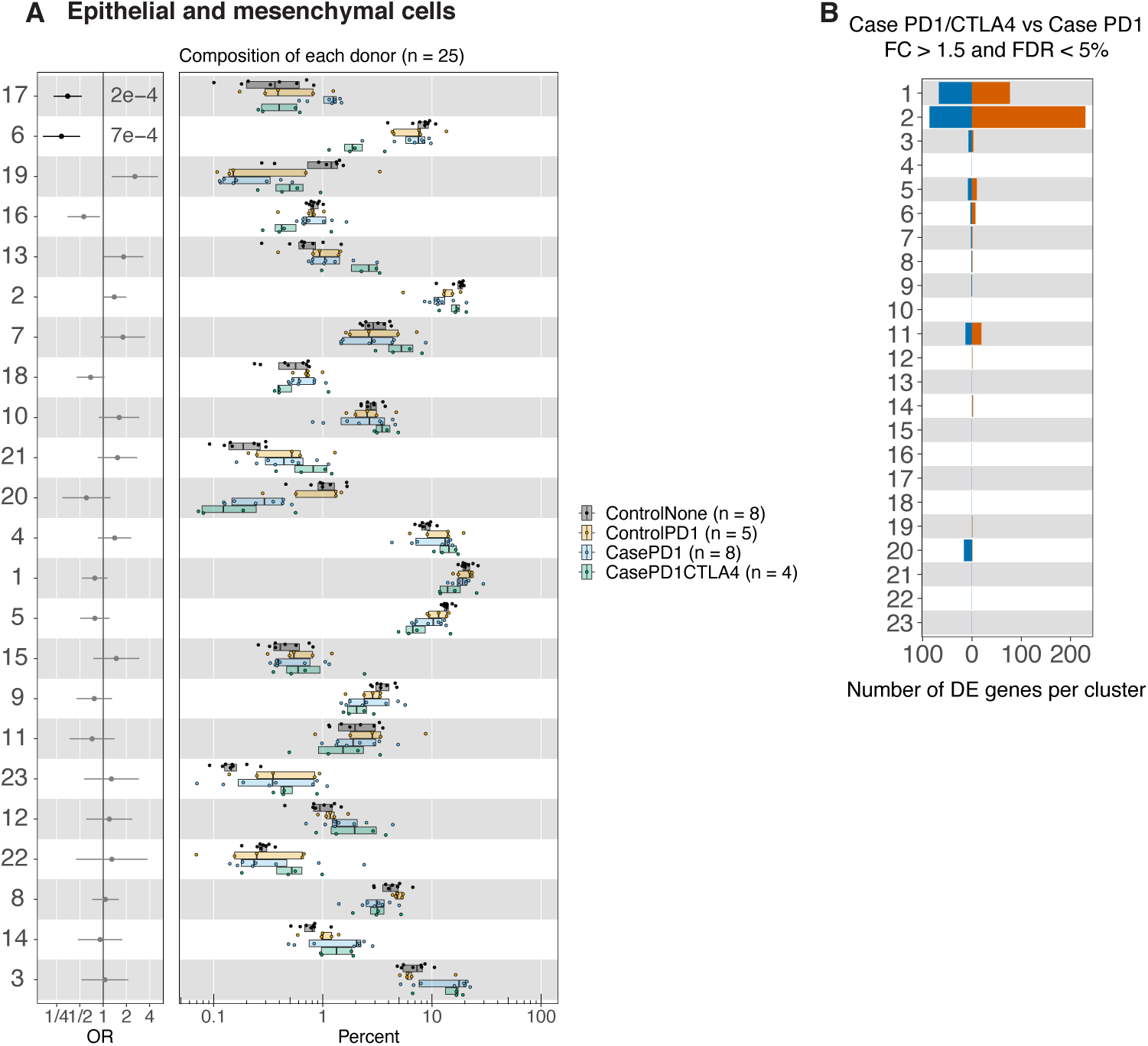
Cell type abundance and differential gene expression analysis of colon tissue epithelial and mesenchymal cells from patients on anti-PD-1 vs anti-PD-1/CTLA-4 therapy. **B.** Left panel shows logistic regression odds ratio (OR) for differential abundance of cells from Cases on PD-1/CTLA-4 versus Cases on PD-1 within each cell cluster. Right panel shows one dot for each patient, indicating the percent of the patient’s cells assigned to each cluster. Likelihood ratio test p-values are shown for clusters with false discovery rate less than 5%. **C.** Number of differentially expressed genes for each cell cluster (fold-change greater than 1.5, false discovery rate less than 5%).

**Figure S19.**
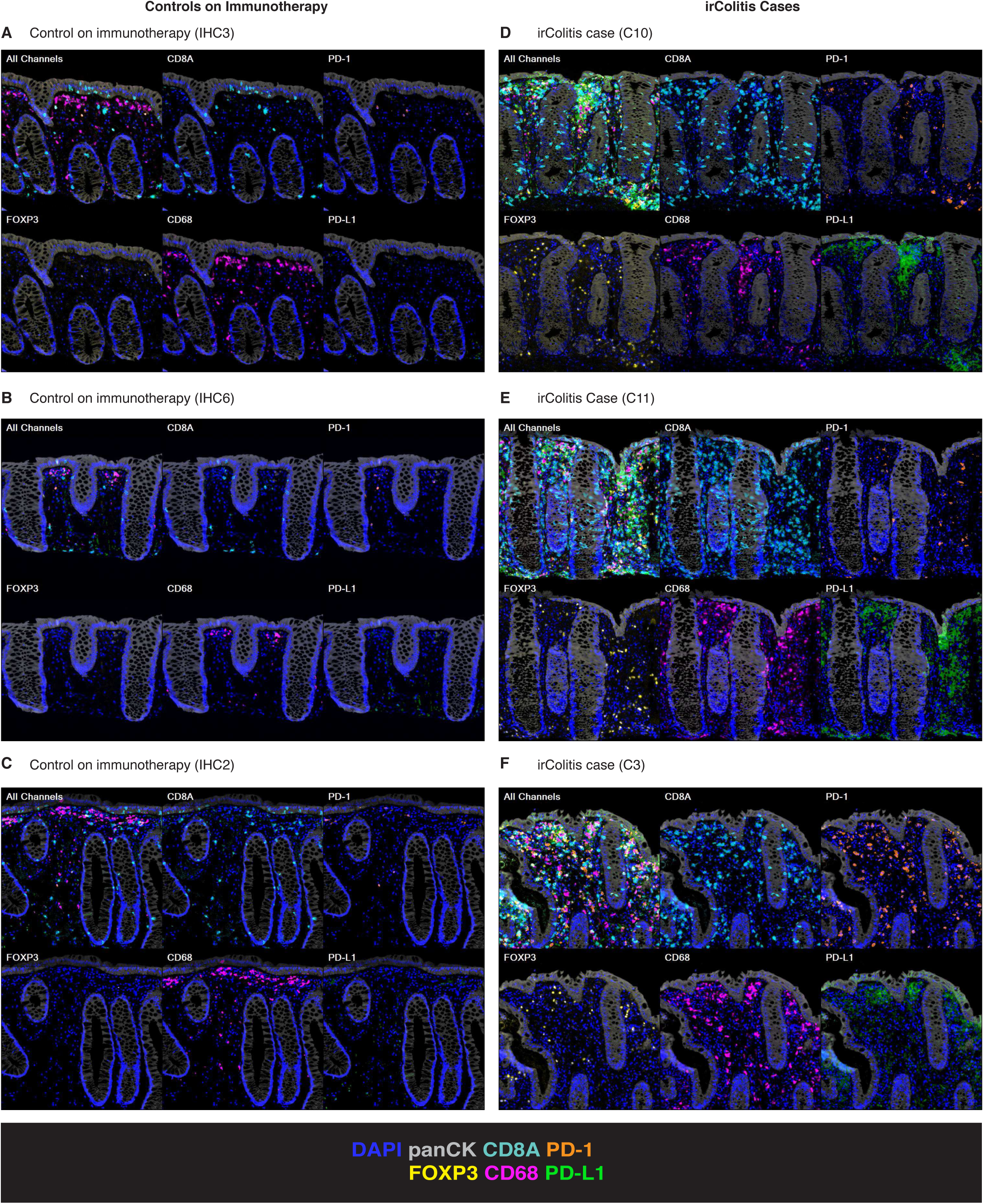
Multispectral immunofluorescence staining of PD-1, PD-L1, and immune cell lineages in colon mucosal tissue biopsies. **A, B, C, D, E, F**. 7-color panel that included DAPI (blue), panCK (grey), CD8A (aqua), PD-1 (orange), FOXP3 (yellow), CD68 (pink), and PD-L1 (green) across (**A, B, C**) controls and (**D, E, F**) irColitis cases.

**Figure S20.**
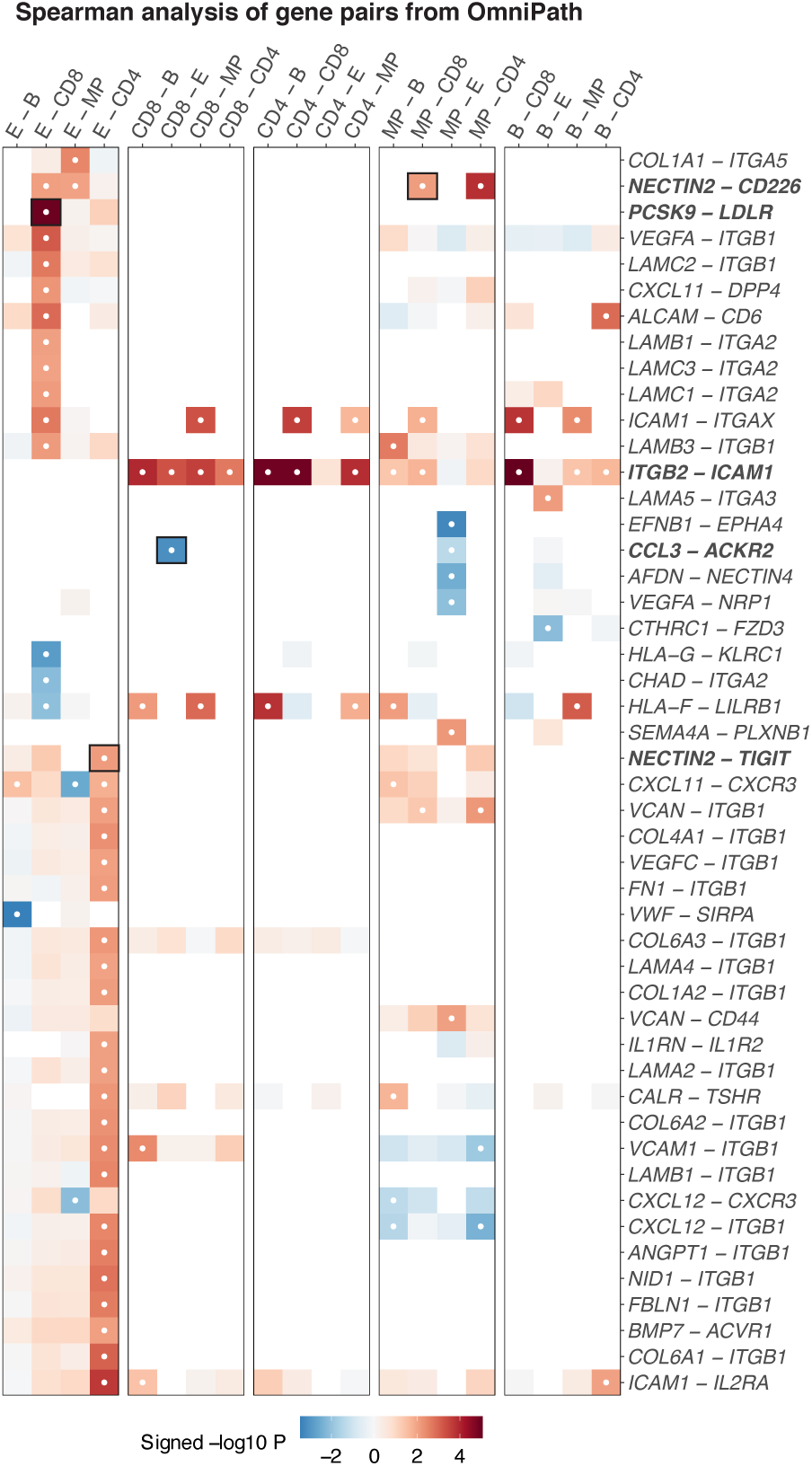
Spearman analysis of ligand receptor gene pairs for colon tissue cell types. Spearman correlation of percent of cells with gene expression for pairs of genes, for each pair of major cell lineages. Heatmap color depicts the Spearman *p-*value signed by the sign of the Spearman correlation coefficient for each pair of genes. White dots indicate FDR <5%. Columns indicate pairs of different cell lineages (E = epithelial, B = B cell, MP = mononuclear phagocyte cell, CD8 = CD8 T cell, CD4 = CD4 T cell). Pairs of genes in boldface are shown in panel Fig. 7C.

## Supplemental Tables

**Table S1.** Patient profiles and summary of which assays were applied to each patient sample

**Table S2.** Cluster names for all cell clusters from colon tissue and blood

**Table S3.** Cell cluster marker gene differential expression statistics for one-versus-all cell comparisons, for all cell clusters from tissue and blood

**Table S4.** Cell cluster differential abundance analysis statistics for three contrasts: Case vs Control, Case PD-1/CTLA-4 vs Case PD-1, Control PD-1 vs Control None

**Table S5.** Differential gene expression analysis statistics for case versus control, for all cell types and clusters

**Table S6.** Number of CD8 T cell TCRs in blood, tissue, and both

**Table S7.** Luminex measurements of 39 analytes from 30 serum samples

**Table S8.** Differential gene expression analysis statistics for Case PD-1/CTLA-4 vs Case PD-1 and for Control PD-1 vs Control None, for all cell types and clusters

**Table S9:** Cell-cell communication results for differential expression analysis of sums of gene pairs

**Table S10:** Cell-cell communication results for Spearman correlation of percent of cells with gene expression

**Table S11.** Sequencing information for scRNA-seq, TCR, and BCR data for immune cells from colon tissue

**Table S12.** Sequencing information for scRNA-seq, CITE-seq, TCR, and BCR data for immune cells from blood

**Table S13.** Catalog numbers of hashtag antibody clones from Biolegend

**Table S14.** Descriptions of CITE-seq (TotalSeq-C) antibodies from Biolegend

**Table S15.** Sequencing information for single-nuclei RNA-seq data for epithelial and mesenchymal cells from colon tissue

**Table S16.** Names and versions of all software packages used for data analysis

## Methods

### Patient selection, endoscopic tissue acquisition, blood and serum collection, PBMC isolation

Endoscopic colon biopsies and blood were obtained from irColitis patients, patients on immunotherapy without colitis, patients with chronic diarrheal symptoms, and healthy controls at Massachusetts General Hospital (MGH). Informed consent was obtained from all patients in accordance with protocols obtained from the Mass General Brigham and/or DANA- Farber/Harvard Cancer Center Institutional Review Board (DFCI/HCC 11-181 and 13-416, Mass General Brigham 2015P001333). Subjects included both males and females spanning ages 30 – 81. Patients were excluded if they were on immunosuppressive medications (other than chronic low-dose prednisone ≤ 10 mg daily for non-colitis indication), antibiotics, had clinical evidence of an active infection including infectious colitis, biopsy-proven ischemic colitis, or pre-existing microscopic colitis, Crohn’s disease, or ulcerative colitis. All cases of irColitis were confirmed by a clinical pathologist. Clinical metadata for each patient can be found in **Table S1**. Tissue biopsies were obtained during colonoscopy or flexible sigmoidoscopy with standard endoscopic biopsy forceps and were taken from the descending colon, sigmoid colon, and rectum (unless otherwise indicated, **Table S1**). For each patient, 6 tissue biopsies from each endoscopic region were placed in HypoThermosol medium (Stemcell Technologies 07935) and were transported on ice to the laboratory for further processing.

Blood was collected from patients undergoing colonoscopy no more than 60 minutes before their endoscopic procedure when tissue biopsies were collected. For selected patients with irColitis, follow-up blood samples were collected at an indicated time of routine clinical blood draw. Blood was collected in EDTA and heparin-coated serum vacutainer tubes (BD 366643, 668660) and processed within 4 hours of collection. PBMCs were isolated from EDTA collection tubes using Ficoll-Paque density gradient centrifugation and frozen at -80°C in a 1:1 mixture of cells suspended in RPMI with 2% (v/v) human AB serum and CryoStor (BioLife Solutions 210102). Serum vacutainers were positioned upright for at least 30 minutes at room temperature, centrifuged at 2100 *g*, and supernatant was stored at -80° C.

### Preparation of single cell suspensions from colon biopsies

2-3 Biopsies collected from multiple subregions of the colon were pooled together in equal proportion into phenol-free RPMI media (ThermoFisher 11835055) containing 1 mg/ml cold active protease from *Bacillus licheniformis* (MilliporeSigma P5380), 5 mM CaCl2, and 0.1 mg/ml DNAse I (MilliporeSigma DN25). Biopsies were cut into ≤ 1 mm pieces with standard laboratory tissue dissection scissors and incubated on a spinning rotor at 4°C for 40 minutes with tissue resuspension every 10 minutes using a P1000 pipette. Reaction was quenched by adding 10% (v/v) human AB serum (MilliporeSigma H4522). Digestion products were then passed through a 70 µM nylon mesh filter to give uniform cell suspensions, centrifuged at 350 *g* for 10 minutes, and resuspended in phenol-free RPMI. Cells were counted with a hemocytometer yielding at least 200,000 viable cells/biopsy from patients with irColitis and at least 100,000 viable cells/biopsy from patients without colitis. A portion of the cell suspension was then aliquoted for further CD45^+^ cell selection.

### Tissue biopsy flow cytometry, immune cell isolation, and preparation of tissue immune cells for 10x scRNA-seq

Colon mucosal CD45^+^ immune cells were isolated from single cell suspensions using either bead-based selection or sorting by flow cytometry (**Table S11**). For bead-based selection, single cell suspensions of enzymatically-digested colon mucosal biopsies were depleted of dead cells using Annexin V conjugated beads (Stemcell 17899). Red blood cells were then depleted from the suspension using antibodies directed against glycophorin A (Stemcell 18352). CD45^+^ cells were then positively selected using CD45 antibodies conjugated to magnetic beads (Biolegend MojoSort CD45 nanobeads 480029). Viable cells were counted on a hemocytometer. At least 20,000 live immune cells were isolated from each patient for downstream analysis.

For selection of CD45^+^ cells by flow cytometry, single cell suspensions of enzymatically digested biopsies were brought up in phenol-free RPMI with 2% (v/v) human AB serum and were incubated on ice for 30 minutes with the following antibodies: CD66b-FITC (1:100, Biolegend 305104), EpCAM-PE (1:100, Biolegend 324206), CD45-APC (1:150, Biolegend 304012), CD3 PerCP-Cy5.5 (1:150, Biolegend 300328), and CD235a PE-Cy7 (1:150, Biolegend 349112). Cells were washed once and resuspended in phenol-free RPMI with 2% (v/v) human AB serum containing DAPI (ThermoFisher 62248). Single-color controls were performed with these same antibodies using BD CompBeads (BD Biosciences 552843). Live, singlet CD66b^−^, CD235a^−^, EpCAM^−^, CD45^+^ cells were sorted into phenol-free RPMI with 2% (v/v) human AB serum. All sorting was performed on Sony SH800 or MA900 Cell Sorters. At least 20,000 immune cells were sorted from each patient. Data were analyzed using FlowJo 10.4.2 (BD Life Sciences).

Isolated CD45^+^ cells were centrifuged and resuspended at a concentration of ∼500 - 1000 cells/µl in RPMI with 2% (v/v) human AB serum in preparation for loading on the 10X Chromium instrument.

### PBMC CD45^+^ and CD8^+^ T cell enrichment, cell hashing, CITE-seq staining

PBMCs cryopreserved and stored at -80°C from eight patient samples at the time (for a total of 23 samples; **Table S12**) were thawed at 37°C, transferred to individual 15 ml conical tubes with 10x volumes of RPMI with 10% (v/v) heat-inactivated FBS (Sigma) followed by centrifugation at 300 x g for 7 minutes. The cell pellet was resuspended in FACS buffer [ 1x PBS (ThermoFisher Scientific, 10010023), 2.5% (v/v) heat-inactivated FBS, 2mM EDTA] and transferred to a 96-well U-bottom plate (Corning Costar Assay Plate, 3788). Single cell suspensions were depleted of dead cells and red blood cells using a bead-based Annexin-V-conjugated bead kit (Stemcell 17899) and glycophorin A-based antibody kit (Stemcell 01738), respectively, with modifications made to the manufacturer protocol to decrease sample volume size to 150 µl. Resulting live cells were counted on the Bio-Rad TC20 automated Cell Counter with trypan blue, and 250,000 cells were resuspended in 100 µl of FACS buffer containing TruStain FcX blocker (Biolegend 422302) and MojoSort CD45 Nanobeads (Biolegend 480030). Individually titrated working solutions containing one of eight hashtag antibodies were added to each of the eight respective patient samples (Biolegend; **Table S13**) and incubated on ice for 30 minutes followed by three magnet-based washes in FACS buffer. Live cells were counted on the Bio-Rad counter with trypan blue, and 60,000 cells from each of the eight patient samples were combined together for a total of 500,0000 cells per pool. The same approach delineated above was employed to hash CD8^+^ T cells enriched from each patient, with the caveat that CD8^+^ enrichment was achieved by incubating 1-2 million PBMCs with CD8 Mojobeads (Biolegend 480108) for 15 minutes on ice. The remainder of the downstream protocol remained the same.

CD45^+^ and CD45^+^ CD8^+^ cell suspensions respectively containing 500,000 cells from eight patients were subsequently filtered through a 40 µM strainer and centrifuged at 300 x g for 7 minutes, placed on a magnet, and resuspended in 25 µl of TotalSeq-C antibody cocktail (Biolegend; **Table S14**). Cells were incubated on ice for 30 minutes and washed 4 times in 1.5 ml FACS buffer. The cell pellet was resuspended in RPMI with 10% (v/v) FBS, filtered again through a 40 µM strainer, and the live cells were counted and resuspended at a concentration of 1200 cells/µl. A total of 50,000 cells were loaded per 10X channel to generate downstream single-cell gene expression, TCR, BCR, and surface protein CITE-seq libraries (**Table S12**).

### Single nuclei isolation from colon tissue biopsies

Endoscopic colon biopsies were collected as indicated above. Individual biopsies intended for nuclei isolation were placed into cryo-vials, flash frozen on dry-ice, and stored at -80°C. Nuclei were extracted at a later date using dounce homogenization and lysis in a tween-based buffer as previously described (Drokhlyansky et al. 2020). In brief, flash frozen biopsies were thawed, rinsed in PBS, and dounce homogenized with a 2 ml Dounce Tissue Grinder (Sigma D8938) 20 times with pestle A and 20 times with pestle B. Tissue homogenization was performed in a lysis buffer containing 10% (w/v) tween, ST buffer (146 mM NaCl, 1 mM CaCl_2_, 21mM MgCl_2_, 10 mM Tris-HCl pH 9), and RNAse inhibitor (Takara 2313, diluted 1:1000). Homogenate was then passed through a 40 µM filter to remove debris, centrifuged at 500 x *g*, resuspended in ST buffer with RNAse inhibitor, and filtered through a 10 µM filter. Nuclei were counted with bright field microscopy and resuspended at a concentration of ∼500 – 1000 nuclei/µl in ST buffer in preparation for loading on the 10X Chromium instrument (**Table S15**).

### Droplet-based scRNA-seq

Single immune cell or nuclei suspensions from colon tissue and hashed CD45^+^ and CD45^+^CD8^+^ PBMC were generated as indicated above (**Tables S11, S12, and S15**). An input of 12,000 single cells or nuclei from tissue was added to each channel with a recovery goal of approximately 4,000 single cells or nuclei. An input of 50,000 hashed PBMC samples was loaded on the 10X Chromium instrument for blood immune cell analysis for a recovery goal of 30,000 single cells. Suspensions were then loaded along with reverse transcriptase reagents, 3’ or 5’ gel beads, and emulsification oil onto separate channels of a Single Cell A Chip, which was loaded into the 10X Genomics Chromium Controller instrument to generate emulsions. Emulsions were transferred to PCR strip tubes for immediate processing and reverse transcription. Library preparation was performed according to manufacturer’s recommendations.

All tissue-derived immune cell and nuclei gene expression libraries (**Tables S11, S12, and S15**) were generated with the Chromium Single Cell 3’ (V2, 10X genomics PN-120237) or 5’ (V1, 10X Genomics PN-1000006) kits according to the manufacturer’s protocols. All hashed PBMC single cell libraries were generated with the Chromium Single Cell 5’ (V1.1, 10X Genomics PN-1000020) together with the 5’ Feature Barcode library kit (10X Genomics PN-1000080). For immune cells libraries generated with the Chromium Single Cell 5’ V1 or 5’ V1.1 kits, PCR-amplified cDNA was used for TCR and BCR enrichment with the Chromium Single Cell V(D)J Enrichment kit (10 Genomics PN-1000005 and PN-1000016). cDNA and library quality were evaluated using an Agilent 2100 Bioanalyzer. Gene expression libraries were sequenced on an Illumina Nextseq instrument and TCR/BCR-enriched libraries were sequenced on an Illumina MiSeq instrument. All hashed and CITE-seq PBMC libraries were sequenced on an Illumina Novaseq instrument.

### Luminex

Serum secreted protein measurements were obtained using a custom Luminex ProcartaPlex Multiplex kit (Thermo Fisher Scientific) to detect the following 39 analytes: CXCL13, CCL11, GM-CSF, CXCL1, IFN-α, IFN-γ, IL-1a, IL-1b, IL-1RA, IL-2, IL-4, IL-5, IL-6, IL-7, CXCL8, IL-9, IL-10, IL-12p70, IL-13, IL-15, IL-17A, IL-18, IL-21, IL-22, IL-23, IL-27, IL-31, IL-33, CXCL10, CCL2, CXCL9, CCL3, CCL4, CXCL2, CCL5, SDF-1a, CCL25, TNF-α, and TNF-β. Secreted proteins were detected per manufacturer’s protocol. In brief, human serum stored at -80°C was thawed, centrifuged at 1000xg, and all supernatants were diluted 1:2. All samples were aliquoted in duplicate to the same plate to avoid batch effects. Standard curves were prepared by serial dilutions of standards corresponding to secreted factor patterns supplied by the manufacturer. Plate was read with a FLEXMAP 3D instrument (Luminex) via xPONENT software. Cloud-based Procartaplex Analysis Software was used to calculate standard curves and mean fluorescence intensity (MFI) for each detected protein, and analyte concentrations (pg/mL) were averaged across duplicates. We tested for differences in irColitis cases versus controls for each analyte with a two-tailed t-test on log10 (concentration).

### RNA and Immunofluorescence Microscopy

5 µm sections were cut from formalin-fixed paraffin-embedded endoscopically-collected colon mucosal tissue blocks onto SuperFrost plus slides (Fisher Scientific 1255015). Slides were baked at 65°C for 2 hours before use. Protein immunofluorescence staining for CD8A, CD68, FoxP3, PD-1, PD-L1, and PanCK was performed using the Motif PD-1/PD-L1 panel (Akoya Biosciences OP-000001) on a Leica Bond Rx instrument (Leica Biosystems) per manufacturer’s protocol. Mixed RNAscope *in situ* hybridization and antibody antigen retrieval and staining was performed with RNAscope probes to *CXCL13*, *CXCL10/11*, *IFNG*, and *CD3E* and antibody to panCK on a Leica Bond Rx instrument using the RNAscope LS multiplex Fluorescent v2 Assay combined with Immunofluorescence protocol (Advanced Cell Diagnostics (ACD) 322818-TN) as described previously (Pelka et al. 2021). The only two variations from the written protocol were (1) an open wash dispense after the peroxide step and (2) DAPI (Sigma D9542) was dispensed twice at the end of the protocol at a concentration of 1 µg/mL. RNAscope probes included the following: CXCL13 (RNAscope LS 2.5 Probe-Hs-CXCL13, ACD 311328), CXCL10 (RNAscope LS 2.5 Probe-Hs-CXCL10-C2, ACD 311858-C2), CXCL11 (RNAscope LS 2.5 Probe-Hs-CXCL11-C2. ACD 312708-C2), IFNG (RNAscope LS 2.5 Probe-Hs-IFNG-C3, ACD 310508-C3), and CD3E (RNAscope LS 2.5 Probe-Hs-CD3E-C4, ACD 553978-C4). Immunofluorescence was performed using mouse anti-PanCK AE1/AE3 antibody (Agilent, M3515) and secondary Opal polymer HRP Ms + Rb (Akoya ARH1001EA). RNA probes and panCK antibody were detected with the following Opal Fluorophores pairings: CXCL13 Opal 480 (Akoya FP1500001KT), CXCL10/11 Opal 520 (Akoya FP1487001KT), CD3E Opal 620 (Akoya FP1495001KT), IFNG Opal 690 (Akoya FP1497001KT), and panCK Opal 780 (Akoya FP1501001KT). Slides were rinsed with water (Fisher 23-751628) prior to coverslipping (Fisher 12-544C) with mountant (Life Technologies P36961).

All slides were imaged using a Vectra Polaris microscope (Akoya Biosciences). Raw Vectra Polaris Images for each slide were unmixed with inForm software (Akoya Biosciences) using an algorithm built on a library of fluorescence spectra measured using single fluorophore-labeled control slides. Unmixed multi-layer image TIFFs from single fields of view were then stitched together into a single multilayer TIFF using Halo software (Indica Labs).

### Computational data analysis

All R software package names and versions used for data analysis can be found in **Table S16**.

### Pre-processing of single cell RNA-seq data

We used 10X Genomics Cell Ranger (version 3.1.0) to pre-process raw single-cell RNA-seq data, including demultiplexing FASTQ reads, aligning reads to the human reference genome (GRCh38, version 3.0.0 from 10X Genomics), and counting the unique molecular identifiers (UMIs) to produce a count matrix with one row for each gene and one column for each cell. We used R scripts to aggregate the data and filter out poor quality cells that had fewer than 500 genes detected (with at least 1 read) or more than 30% of reads from 13 mitochondrial genes (MT-ND6, MT-CO2, MT-CYB, MT-ND2, MT-ND5, MT-CO1, MT-ND3, MT-ND4, MT-ND1, MT-ATP6, MT-CO3, MT-ND4L, MT-ATP8).

### Normalizing count data

After excluding poor quality cells, we normalized the sequencing depth of each cell by dividing each cell’s counts by the total counts in that cell, resulting in a matrix where the entries represent the proportion of a cell’s reads allocated to each gene (i.e. values in the range [0,1]). To estimate a library size for each dataset, we summed the total counts in each cell, and then we took the median as the library size for the dataset. Next, we multiplied the proportions by the library size to get a count matrix that was normalized for sequencing depth. Finally, we transformed the normalized count matrix with log2(1 + count). We referred to this log-transformed quantity in the figures as log2CPM.

### Principal component analysis

Briefly, we selected overdispersed genes, centered and scaled each gene, computed principal components, and ran the molecular cross validation (MCV) algorithm to select the number of components for downstream analysis (Batson, Royer, and Webber 2019).

To identify overdispersed genes, we first used the raw counts matrix to compute a mean with Matrix::rowMeans() and standard deviation with proxyC::rowSds() for each gene, excluding genes that are detected in fewer than 50 cells. Next, we fit a local 2nd degree polynomial regression with stats::loess() with the formula “log10(sd) ∼ log10(mean)” to model the relationship between mean and variance for each gene. We computed the residual variance for each gene, and selected the top 80% of genes with the greatest residual variance.

With the selected genes, we centered and scaled each gene to have mean equal to zero and variance equal to one. We used RSpectra::svds() to compute the truncated singular value decomposition (SVD) of the scaled expression matrix. Then, we multiplied the left singular vectors by the singular values to obtain principal component scores for each cell.

To select the optimal number of principal components, we followed the molecular cross validation algorithm. For a raw count matrix, we randomly split the counts for each cell, resulting in two count matrices A and B whose sum was equal to the original matrix. With the centered and scaled matrix A’, we computed principal components as described above. Then, we reconstruct matrix A’ from the top K PCA scores and loadings (UV^T^) and estimated a reconstruction loss with the mean squared error. We also reconstructed matrix B’ from the PCA scores that were computed on matrix A’ to estimate a molecular cross validation (MCV) loss. We recorded the losses for each choice of top K principal components between 2 and 80. We repeated this procedure three times, resulting in three estimates of the MCV loss for each choice of the top K. Then, we chose the K that minimized the average MCV loss. These K principal components were used in downstream analyses.

### Cell clustering and two-dimensional embedding

We used the Harmony algorithm to align PCA scores across batches of data (Korsunsky, Millard, et al. 2019), and then we created a nearest neighbor network of cells by connecting each cell to the 50 cells with the least Euclidean distance in the space of the harmonized PCA scores. We ran the Leiden algorithm (Traag, Waltman, and van Eck 2019) from the leidenalg Python package on the nearest neighbor network to identify cell clusters (10 iterations). We ran the UMAP algorithm (McInnes, Healy, and Melville 2020) with uwot::umap() to embed the network in two dimensions (spread = 1, min_dist = 0.25).

### Cell type abundance analysis

We fit a logistic regression model with lme4::glmer() to determine whether the cells that belong to a given cluster were enriched or depleted from cases (Fonseka et al. 2018). For each cluster, we defined a boolean vector Y with one value for each cell that is true if the cell belonged to the cluster and false otherwise. Then, we fit a null model *Y ∼ 1 + (1|donor)* where *(1|donor)* indicates a random effect that gives each donor a different intercept. Next, we fit an alternative model *Y ∼ 1 + case + (1|donor)* where *case* is a boolean vector that is true if the cell belongs to a case donor. We used the likelihood ratio test with stats::anova() to compare the two models and record the *p-*values. We use the Benjamini & Hochberg (1995) method to compute false discovery rate (FDR) for the *p-*values and report results with FDR less than 5 percent as statistically significant. In the figures, we report the odds ratio (OR) and 95% confidence interval for each cluster.

For the comparison of controls on PD-1 versus controls without ICI therapy, we only used the cells that belonged to control donors. Here, we fit the null model *Y ∼ 1 + (1|donor)*. The alternative model is *Y ∼ 1 + drug + (1|donor)* where *drug* is a binary indicator with levels *PD1* and *None*.

For the comparison of dual therapy (anti-PD-1 and anti-CTLA-4) versus single therapy (anti-PD-1), we used only the cells that belong to case donors on single or dual therapy. Here, we fit the null model *Y ∼ 1 + sex + (1|donor)* where *sex* is a binary indicator with two levels *M* and *F*. The alternative model is *Y ∼ 1 + sex + drug + (1|donor)* where *drug* is a binary indicator with levels *PD1* and *PD1CTLA4*.

### Differential gene expression analysis

We created pseudobulk expression (L. Lun, Bach, and Marioni 2016) for the cells in a cluster for each donor such that the pseudobulk matrix had one row for each gene and one column for each cluster from each patient. We normalized the pseudobulk counts to log2CPM as described in the previous section. Then we use limma::lmFit() to test for differential gene expression with the log2CPM pseudobulk matrix (Ritchie et al. 2015). We also use presto::wilcoxauc() to compute the area under receiver operator curve (AUROC or AUC) for the log2CPM value of each gene as a predictor of the cluster membership for each cluster (Korsunsky, Nathan, et al. 2019).

For the comparison of dual therapy (anti-PD1 and anti-CTLA4) versus single therapy (anti-PD1), the pseudobulk expression was created by summing over all cells in a lineage (Tissue CD8, Tissue CD4, Tissue Myeloid, Tissue B cell, Tissue Non-Immune) rather than summing over clusters within each cell lineage. The linear model fit to each gene is *Gene ∼ 0 + case_drug + sex* where *case_drug* is a categorical variable with four levels *ControlNone*, *ControlPD1*, *CasePD1*, *CasePD1CTLA4* and *sex* is a binary indicator with levels *M* and *F*. We then use limma::contrasts.fit() to test three contrasts *ControlPD1 - ControlNone*, *CasePD1 - ControlPD1*, and *CasePD1CTLA4 - CasePD1*. We reported the contrast *CasePD1CTLA4 - CasePD1* in the figures. We excluded spurious genes with low mean expression or detection in less than 2% of cells.

### Differential expression analysis of sums of gene pairs

We obtained gene pairs from OmniPath (Türei et al. 2021) with OmnipathR::import_intercell_network() and then filtered to the pairs with the following criteria: consensus_score_intercell_source >= 3, curation_effort >= 1, n_resources >= 4. After filtering, we had 1,827 gene pairs that were likely to represent ligand-receptor pairs. Next, we looped through each pair of cell types (Tissue CD8, Tissue CD4, Tissue Myeloid, Tissue B, Tissue Non-Immune) and looped through each gene pair, and summed the pseudobulk log2CPM values for the two genes. Finally, we used limma::lmFit() to fit the model *Gene ∼ 1 + case* where *Gene* is the sum of the log2CPM values of the gene pair and *case* is a binary indicator with levels *Case* and *Control*.

### Spearman correlation of percent of cells with gene expression

We computed the percent of cells with non-zero expression of each gene for each cell lineage (Tissue CD8, Tissue CD4, Tissue Myeloid, Tissue B, Tissue Non-Immune) in each donor. Then, we computed the Spearman correlation coefficient and *p-*value for each gene pair for each pair of cell types with Hmisc::corr(). This tested for correlation across all of the donors, so the result indicates whether the percent of cells in cell type 1 for gene 1 is correlated with the percent of cells in cell type 2 for gene 2. We used the same list of 1,827 gene pairs as in the previous section.

### TCR and BCR diversity analysis

We defined a unique TCR sequence as the concatenation of four pieces: TCRɑ V-gene, TCRɑ CDR3 amino acid sequence, TCRβ V-gene and TCRβ CDR3 amino acid sequence. Clones missing any of these features were not included in the analyses. Next, we used alakazam::alphaDiversity() to compute the Hill numbers of order q (effective number of species) (Gupta et al. 2015). For BCR analysis, we defined a unique BCR clone as the heavy-chain CDR3 amino acid sequence and light chain CDR3 amino acid sequence. Clones that had a recovered heavy chain CDR3 amino acid sequence and either a lambda or kappa light chain CDR3 amino acid sequence were used for the analyses.

### Analysis of TCR sharing between tissue and blood

To estimate whether TCR clones observed in blood CD8 T cells are enriched or depleted from a particular tissue CD8 T cell cluster (or vice versa), we fit a logistic regression model with lme4::glmer() (Bates et al. 2015). For each cluster, we defined a boolean vector Y with one value for each cell that is true if the cell belonged to the cluster and false otherwise. Then, we fit a null model *Y ∼ 1 + (1|donor)* where *(1|donor)* indicates a random effect that gives each donor a different intercept. Next, we fit an alternative model *Y ∼ 1 + tcr_shared + (1|donor)* where *tcr_shared* is a boolean vector that is true if the cell has a TCR that is observed in both blood and tissue cells. We used the likelihood ratio test with stats::anova() to compare the two models and recorded the *p-*values. We computed false discovery rate (FDR) (Benjamini and Hochberg 1995) for the p-values and reported results with FDR less than 5 percent as statistically significant. In the figures, we reported the odds ratio (OR) and 95% confidence interval for each cluster.

